# Genome reorganisation and expansion shape 3D genome architecture and define a distinct regulatory landscape in coleoid cephalopods

**DOI:** 10.1101/2025.08.29.672809

**Authors:** Thea F. Rogers, Jessica Stock, Natalie Grace Schulz, Gözde Yalçin, Simone Rencken, Anton Weissenbacher, Tereza Clarence, Darrin T. Schultz, Clifton W. Ragsdale, Caroline B. Albertin, Oleg Simakov

## Abstract

How genomic changes translate into organismal novelties is often confounded by the multi-layered nature of genome architecture and the long evolutionary timescales over which molecular changes accumulate. Coleoid cephalopods (squid, cuttlefish, and octopus) provide a unique system to study these processes due to a large-scale chromosomal rearrangement in the coleoid ancestor that resulted in highly modified karyotypes, followed by lineage-specific fusions, translocations, and repeat expansions. How these events have shaped gene regulatory patterns underlying the evolution of coleoid innovations, including their large and elaborately structured nervous systems, novel organs, and complex behaviours, remains poorly understood. To address this, we integrate Micro-C, RNA-seq, and ATAC-seq across multiple coleoid species, developmental stages, and tissues. We find that while topological compartments are broadly conserved, hundreds of chromatin loops are species- and context-specific, with distinct regulation signatures and dynamic expression profiles. CRISPR-Cas9 knockout of a putative regulatory sequence within a conserved region demonstrates the role of loops in neural development and the prevalence of long-range, inter-compartmental interactions. We propose that differential evolutionary constraints across the coleoid 3D genome allow macroevolutionary processes to shape genome topology in distinct ways, facilitating the emergence of novel regulatory entanglements and ultimately contributing to the evolution and maintenance of complex traits in coleoids.

## Introduction

Deep conservation of macrochromosomal synteny, traceable to the last common ancestor of animals and beyond, is paralleled by the persistence of numerous subchromosomal syntenic units^1–6^. While whole chromosomes are often constrained by meiotic and gamete-balancing processes, the evolutionary forces that maintain local gene linkages over hundreds of millions of years remain more elusive. Many of these syntenic blocks exhibit divergent genomic properties, including variable gene density and co-expression levels (or lack thereof)^7,8^, but the overarching principles of their evolutionary and regulatory constraints are unclear.

Emerging data from several species are shedding light on the complex 3D organisation of the genome, revealing interacting chromatin domains, commonly referred to as compartments or topologically associating domains (TADs), as key units of genome architecture^9–12^. These domains are thought to constrain regulatory interactions^13^ and influence gene expression, with syntenic breaks frequently aligning with compartment boundaries^14–17^. However, whether compartments themselves are conserved across species remains controversial, with conservation appearing highly method-dependent and subject to multiple confounding variables (reviewed in^18^).

At more localised or structurally simpler scales of genome organisation, chromatin loops bring enhancers and promoters into proximity, often via loop extrusion mechanisms, and may contribute to gene regulation in a context-dependent manner^12^. Their roles in developmental, tissue-, and cell-specific regulation are still being investigated. From an evolutionary perspective, the persistence of certain loops, within or beyond compartments, may result in constraints on regulatory architecture, as suggested by models such as the bystander or co-expression hypothesis, which propose that gene linkages are maintained to coordinate transcriptional programs^2–7^. However, the extent to which chromatin loops are conserved varies across animals^19^, and the evolutionary timing and stability of loop formation in relation to large-scale genomic processes remain unclear. Recent studies have highlighted how chromosomal fusions can reshape 3D genome folding and influence recombination landscapes, offering new evolutionary perspectives^20^. Alternatively, it has been proposed that loop extrusion serves more general biophysical roles in genome organisation, such as preventing the formation of spurious chromatin structures and maintaining global accessibility to diffusible regulatory proteins, rather than contributing primarily to specific, targeted regulatory functions^21^. Thus, studying the evolution of 3D genome architecture, from large-scale chromatin domains to fine-scale looping interactions, has the potential to address long-standing questions about the nature of genomic constraints and gene regulation.

A key to understanding such patterns of 3D genome organisation lies in examining the emergence and persistence of evolutionary changes over macroevolutionary timescales, including chromosomal and sub-chromosomal rearrangements, as well as genome expansion and contraction^11,22^. Few systems currently provide the comparative resolution needed to explore these dynamics.

Coleoid cephalopods (squid, cuttlefish and octopus) represent a powerful and emerging model in this context (Fig. 1A, B). The coleoids are an ancient clade (∼ 450 Ma)^23^ that have evolved the largest invertebrate brains and several lineage-specific innovations, including novel organs and complex behaviours^24–28^. Initial analyses of genome topology have identified compartmental structures with CTCF motifs enriched at TAD boundaries^8^. Several studies have also documented an extensive ancient coleoid genome rearrangement event (Fig. 1C)^29,30^. Furthermore, Decapodiformes (squid and cuttlefish) and Octopodiformes (octopus and vampire squid) lineages also exhibit distinct evolutionary trajectories with respect to genome organisation: although both lineages underwent genome expansion, they did so independently and through activity of different repeat superfamilies^29,31,32^. Moreover, Octopodiformes show evidence of extensive chromosomal fusions^33^. As a result, coleoid genome sizes vary considerably, for example, approximately 2.7 Gb in some octopuses, and over 5 Gb in squid and cuttlefish species (Fig. 1D). These genomic features have been implicated in the evolution of cephalopod innovations^8,29,30^, but the mechanisms by which changes in genome architecture give rise to organismal novelties through gene regulation remain poorly understood. Coleoid cephalopods therefore offer a compelling model to investigate how ancient macroevolutionary processes influence the structure, maintenance, and evolvability of regulatory landscapes.

**Figure 1.**
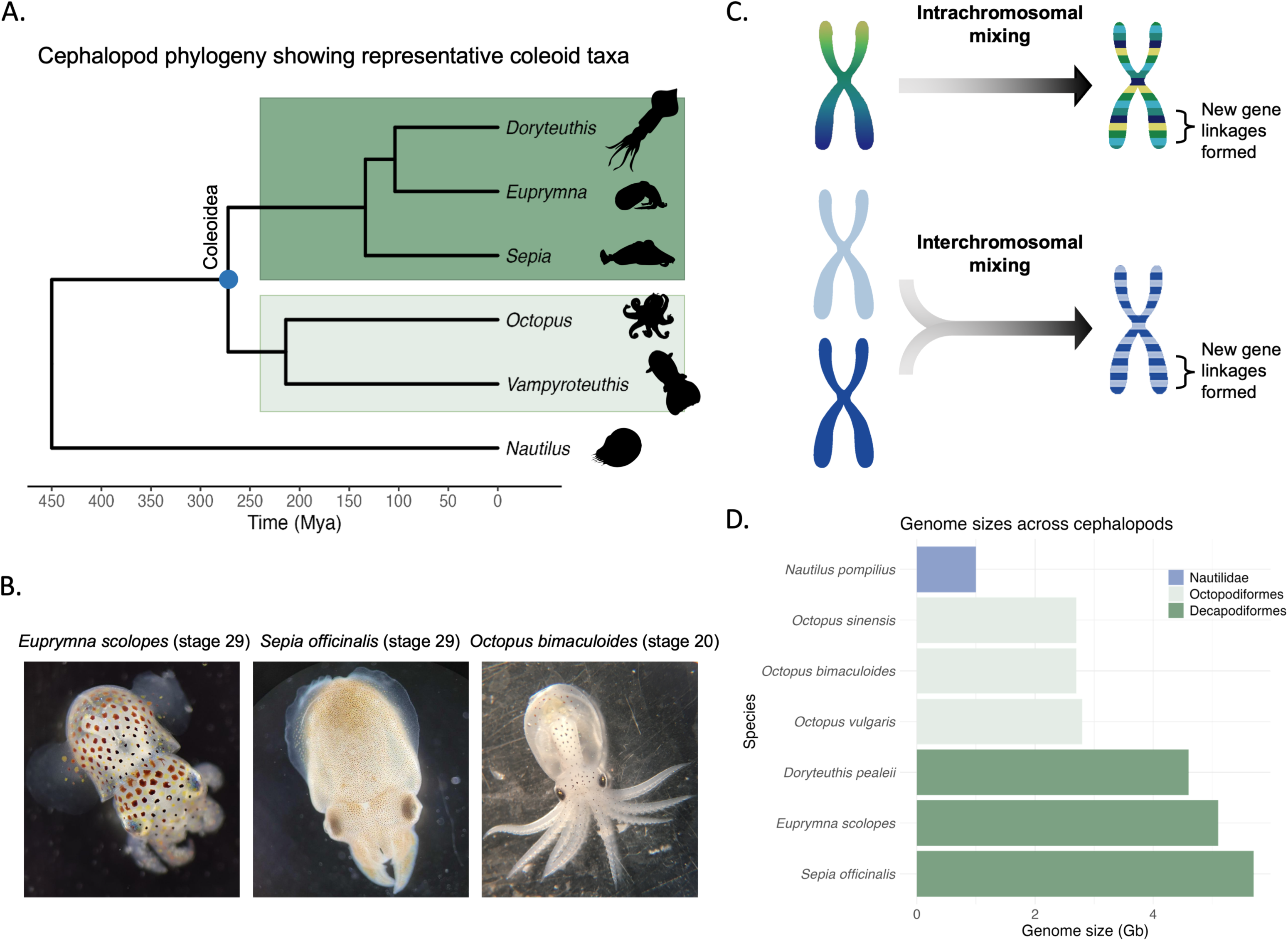
The subclass Coleoidea have undergone large-scale genome rearrangements and significant genome expansions compared to their molluscan ancestors. (A) Phylogenetic relationships among major cephalopod lineages, including representative species from Decapodiformes (dark green) and Octopodiformes (light green), as well as *Nautilus*, a non-coleoid cephalopod species representing the sister clade Nautiloidea^23,30,101^. (B) Representative photos of cephalopod embryos used in this study, showing the developmental stages used for most comparisons: *E. scolopes* (stage 29), *S. officinalis* (stage 29), and *O. bimaculoides* (stage 20), the final stage before hatching in all species. (C) Schematic showing two modes of genomic reorganisation that generated new gene linkages during the ancient chromosomal rearrangement event in the coleoid ancestor: intrachromosomal mixing (within a single chromosome) and interchromosomal mixing (between different chromosomes). (D) Genome sizes across selected coleoid species with chromosome-scale genomes. The nautiloid ancestor *N. pompilius* is shown for comparison.

In this study, we present a comparative analysis of 3D genome architecture across three coleoid species spanning the key lineages within the group: *Euprymna scolopes* and *Sepia officinalis* (Decapodiformes), and *Octopus bimaculoides* (Octopodiformes). By integrating tissue- and developmental stage-specific Micro-C with ATAC-seq, RNA-seq, gene synteny, and conserved non-coding element (CNE) analyses, we uncover a complex pattern of chromatin compartmentalisation that is largely conserved between species. However, compartmental interactions that arose following coleoid genome rearrangements exhibit distinct functional and topological properties compared with those present in the coleoid ancestor. In contrast, chromatin loops show extensive variation across species, tissues, and developmental stages, and are enriched for dynamic regulatory features. Together, these findings reveal a hierarchy of regulatory constraints acting at different levels of 3D genome organisation. Moreover, our results suggest that many novel interactions arise from fusion events, a process particularly pronounced in the octopodiform lineage. Once brought into proximity, these regions may become embedded within nested regulatory architectures, subject to increasing evolutionary constraint, a process we refer to as ‘regulatory entanglement’. By this, we mean the progressive accumulation of interdependencies among genomic elements, shaped by chromosomal rearrangement, spatial proximity and regulatory interactions^22,34^. Such regulatory entanglement can make genes, non-coding elements, and the 3D genome landscape structurally and functionally interwoven, thereby restricting evolutionary trajectories and shaping long-term phenotypic outcomes. Such scenario can explain local synteny retention: a constraint driven by the interplay between genome topology and regulatory dependencies, resulting in hypothesised regulatory entanglements^22,34^. This model suggests that 3D chromatin architecture itself plays a central role in preserving syntenic blocks. Our findings highlight the potential for 3D genome structures to act as synapomorphic traits across lineages, as well as regulatory substrates that guide macroevolutionary trajectories and drive regulatory innovation.

## Results and discussion

### Establishment and diversification of the coleoid genome architecture at coding and non-coding levels

Coleoid chromosomal complements are distinct from other molluscs^30^ and an in-depth syntenic assessment has been lacking. For this, we conducted synteny analysis on orthologous coding genes to identify homologous chromosomes across several coleoid genomes. This analysis confirms general conservation of chromosomal synteny for the three focal species, *E. scolopes*, *S. officinalis*, and *O. bimaculoides* (Fig. 2)^30^. While *S. officinalis* and *E. scolopes* chromosomes showed almost one-to-one correspondence, their homology to *O. bimaculoides* was more complex, indicative of the proposed secondary modifications (fusions and translocations) to the octopod karyotypes^33^ (Fig. 2A). To further quantify the degree of fusions or translocations within chromosomes, we calculated how many 1 Mb, 10 Mb, and 50 Mb windows have single or multiple homologies between the species. For 10 Mb windows, we find 351 instances of pairwise *E. scolopes* to *O. bimaculoides* homologous regions. Additional 80 *O. bimaculoides* regions had a mapping to this shared set, i.e. resulting in two or more correspondences to a single *E. scolopes* regions (Methods). Conversely, there were 686 additional squid regions that corresponded to single regions in *O. bimaculoides.* This suggests that, on average, almost every *O. bimaculoides* 10 Mb homologous region corresponds to two distinct *E. scolopes* regions. This pattern of fragmentation, where single *O. bimaculoides* regions often correspond to two or more *E. scolopes* regions, is consistent across window sizes, though the number of detected events scales with the size of the window. These results point to a high degree of within-chromosomal mixing in the Octopodiformes.

**Figure 2.**
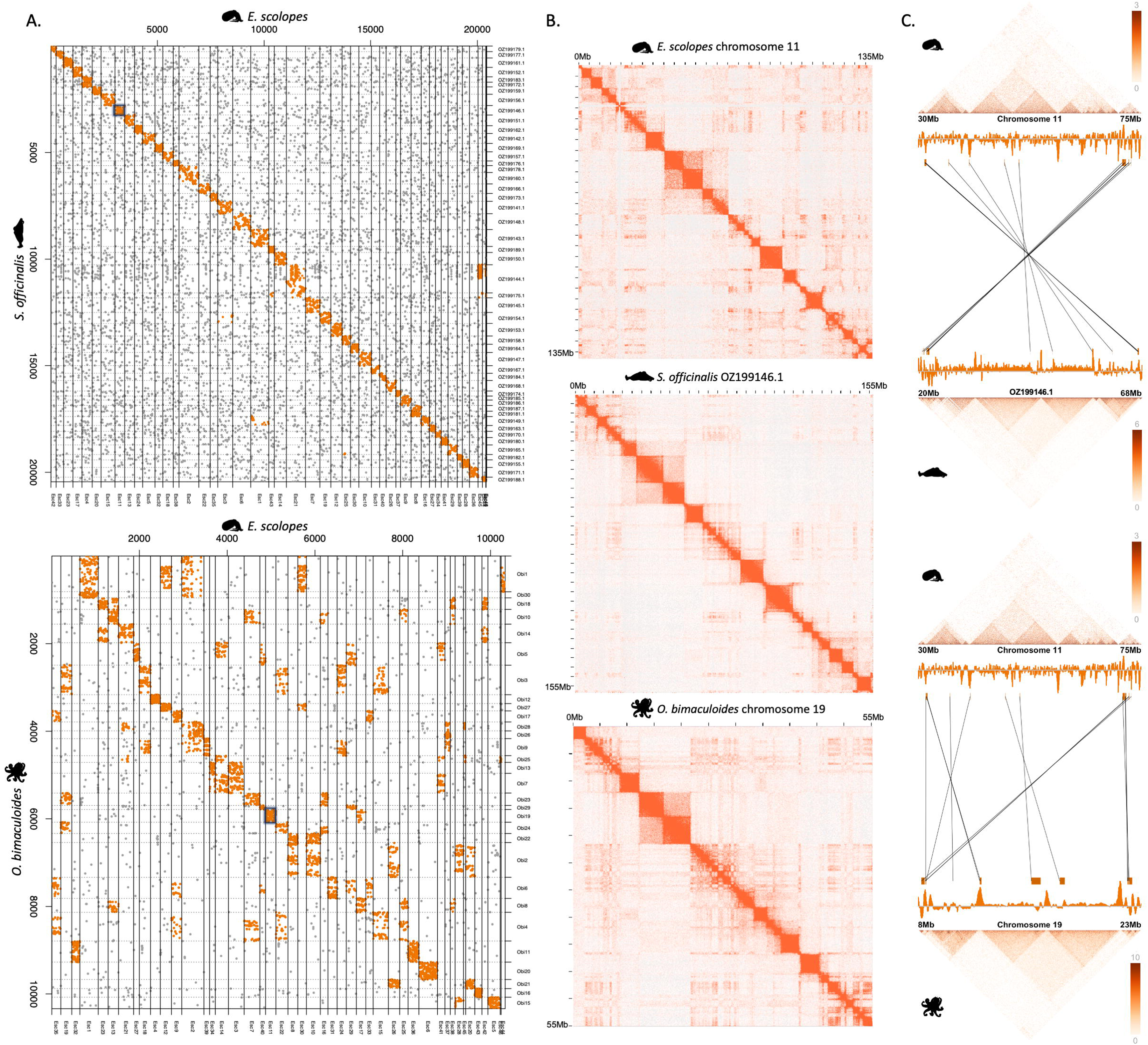
Genome topology and chromosome evolution across coleoids. (A) Dot plots of orthologous gene content comparing *E. scolopes* chromosomes to *S. officinalis* (top) and *O. bimaculoides* (bottom). *O. bimaculoides* and *E. scolopes i*n particular show extensive chromosomal rearrangements, including large-scale inversions (visible as reverse-diagonal synteny blocks), and chromosomal fusions and fissions, as shown by several one-to-many and many-to-one relationships across species. (B) Micro-C contact maps for chromosomes highlighted by a black square in panel A; chromosome 11 in *E. scolopes*, OZ199146.1/chromosome 6 in *S. officinalis*, and chromosome 19 in *O. bimaculoides*, showing 3D genome organisation patterns at 150 kb resolution in the Decapodiform species and 100 kb resolution in *O. bimaculoides.* Compartment-like structures are visible across all species. (C) Cross-species comparisons of Micro-C contact maps and synteny relationships for orthologous chromosome regions, highlighting one-to-one orthologous genes between *E. scolopes*, *S. officinalis*, and *O. bimaculoides*. Genes are only included in the plot if they are a one-to-one orthologs in all species and located within the regions pictured. Insulation scores at a 350 kb window are represented by orange tracks and indicate domain boundaries. Note that the crossed orthology lines in the top panel reflect opposite assembly orientations between the two species rather than a true inversion; gene order within this region is collinear.

To further complement this analysis, we assessed genomic alignment lengths and positions between these species. In total, we found 78 Mb aligned between *E. scolopes* and *S. officinalis* and 19 Mb aligned between *E. scolopes* and *O. bimaculoides* (excluding repetitive regions, Methods). 11 Mb and 4 Mb of the aligned regions, respectively, did not contain any coding sequences and thus comprise putative CNEs. To quantify evolutionary conservation of these regions, we calculated their dispersion across chromosomes. Dispersion is measured as the number of translocated regions across homologous chromosomes and correlates with evolutionary distance^35^. Between *E. scolopes* and *S. officinalis* we found 150,610 and 110,440 total alignments (coding and non-coding) of 100bp or longer on homologous and non-homologous chromosomes, respectively. Between *E. scolopes* and *O. bimaculoides*, 35,273 alignments were on homologous chromosomes and 29,948 alignments were on non-homologous chromosomes, respectively, showing a similar dispersion rate with the absolute numbers scaling with the genetic distance. Interestingly, the dispersion rate for CNEs was higher, with only 3,185 alignments on homologous chromosomes and 22,462 on non-homologous chromosomes for *E. scolopes* and *S. officinalis*, and 528 alignments on homologous chromosomes compared to 6,182 alignments on non-homologous chromosomes for *E. scolopes* and *O. bimaculoides*. This suggests that across coleoids, there are more relaxed constraints on the translocation of putative regulatory non-coding sequences compared with coding regions, likely due to their smaller size, modular function, and reduced sensitivity to disruption relative to protein-coding genes.

While these results highlight the different evolutionary modalities in Octopodiformes compared to Decapodiformes, they cannot provide for a quantitative assessment of how different levels of genome organisation evolved in these clades. For this purpose we applied a recently established evolutionary topology approach on 15 cephalopod species to study multi-scale changes across coleoid cephalopod genomes (multi-locus topology^34^). Using pair-wise distances and manifold projection, this approach allows us to identify localisations of coding and non-coding genes across all chromosomal distance scales. Coding gene pairwise distances showed several disconnected clusters for both Octopodiformes and Decapodiformes (Fig. S1A). Interestingly, the number of clusters on these topological maps were similar between the two clades and in the range of the decapodiform chromosome number, supporting its more ancestral state^33^. However, octopodiform genomes showed more connectivity between the orthologous genes, indicative of the proposed additional chromosomal fusion-with-mixing within the octopodiform lineage, suggesting higher degree sub-chromosomal mixing (regulatory entanglement) and a more derived state. This is supported by the observed higher entropic mixing of these fused chromosomes (Supplementary Note 1). We hypothesise that this more entangled genome architecture in Octopodiformes imposes topological and regulatory constraints that limit the capacity for further genome expansion in this lineage, in contrast to the more extensive genome expansion observed in Decapodiformes. Interestingly, these analyses also show that unlike coding genes that are scrambled across chromosome and sub-chromosomal topological areas, CNEs are less mixed within the Octopodiformes compared to the coding genes, as evidenced by a line-pattern (as opposed to ‘cloud’, Fig. S1B, C). In Decapodiformes, the CNEs are, on the other hand, more scrambled than the coding genes. While this pattern may partly reflect the sparser octopodiform sampling, with fewer octopodiform species than decapodiform species and therefore fewer CNEs spanning deeper evolutionary nodes, it remains consistent with the more regulatory-entangled state of the Octopodiformes after enhanced chromosomal mixing of more ancient genomic regions such as the conserved orthologous genes (Fig. 1A).

Together, this data provide quantification of gene and CNE movements across coleoid chromosomes, revealing a pattern that is highly dependent on both the phylogenetic distance and the evolutionary history of chromosomal fusions and translocations. These data reveal a more derived and mixed (entangled) nature of octopodiform genomes, which will further be assessed below.

### Topological properties of coleoid genomes

To account for inconsistencies in compartment calling, we evaluated 3D genome organisation directly from the normalised Micro-C interaction signal. This approach captures both multi-scale compartmentalisation and nested domain architecture. Overlaying these topological maps onto genomic coordinates revealed conservation of sub-chromosomal regions across species (Fig. 2B, C). To quantify this conservation, we calculated pairwise interaction intensities (derived from the Micro-C matrices; see Methods) between intrachromosomal gene pairs in the coleoids. As expected, when calculating all possible gene pair combinations within chromosomes, the majority of gene pairs (53,538) were not in conserved coleoid interactions. We identified 1,592 gene pairs that interacted (KR normalised interaction strength ≥10, Fig. S2) in all coleoid species, 952 gene pairs interacting exclusively in Decapodiformes, 3,634 gene pairs interacting only in *O. bimaculoides*.

To further assess the roles of patterns of pairwise chromatin interaction types, we assessed their co-expression (Fig. 3A), expression (Fig. S3A-C, Supplementary Note 2, Table S3, S4), and functional enrichment properties (Supplementary Note 3, Fig. S4). In *E. scolopes*, gene pairs interacting across all three coleoid species exhibited the highest correlation in gene expression across *E. scolopes* tissues, followed by those interacting in Decapodiformes only. Gene pairs interacting exclusively in *O. bimaculoides* (i.e., not interacting in *E. scolopes*, where expression is measured) showed the second weakest correlation, while those not in conserved coleoid interactions displayed the weakest correlation (Fig. 3A). All comparisons were statistically significantly different (Wilcoxon test with Benjamini-Hochberg [BH] correction), with adjusted P values ranging from 3.72 × 10⁻²³ (no conserved interactions vs. interacting across coleoids) to 0.042 (interacting across coleoids vs. interacting Decapodiformes only), indicating that co-expression strength scales with the degree of evolutionary conservation difference between interaction categories in each comparison. Conserved chromatin interactions were also found to be associated with significantly higher (Fig. S3B, C, Table S3, Supplementary Note 2) and broader (Table S4, Supplementary Note 2) expression patterns, as well as core housekeeping functions (Supplementary Note 3, Fig. S4). These results suggest that gene pairs maintained in conserved spatial proximity across species are more likely to be functionally coordinated and under strong evolutionary constraints, possibly reflecting shared regulatory mechanisms and core biological roles.

**Figure 3.**
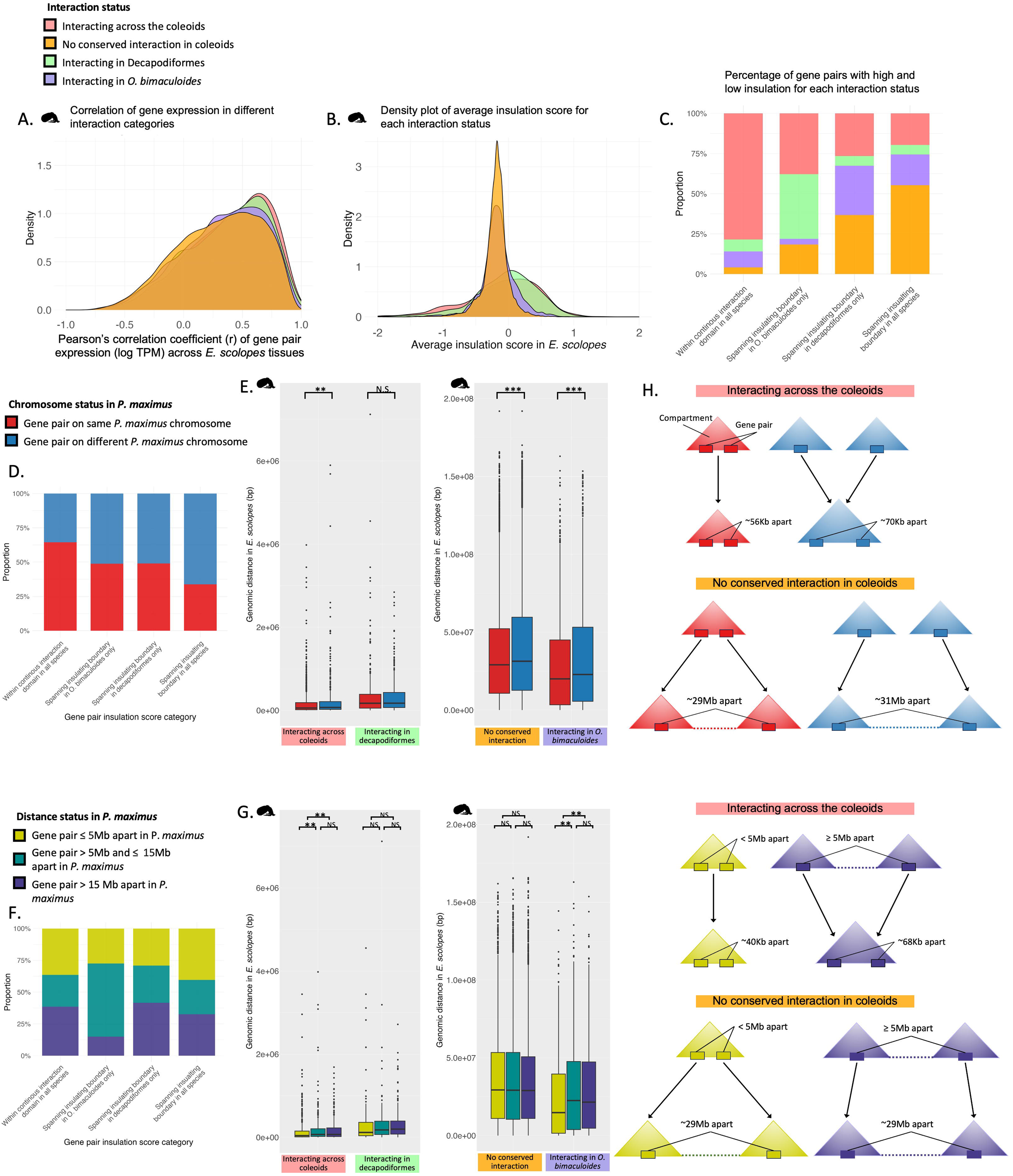
Co-expression, regulatory insulation and intergenic distances for gene pairs across different interaction categories, split by *P. maximus* chromosomal origin. (A) Pearson’s correlation of gene expression across 19 tissues in *E. scolopes* for gene pairs grouped by interaction status. Gene pairs interacting across coleoids show the highest co-expression, and all pairwise comparisons were statistically significant (BH-corrected Wilcoxon test, P < 0.05). (B) Density plot of average insulation score in *E. scolopes* for each interaction category. Insulation scores were computed from *E. scolopes* Micro-C data at 100 kb resolution using a 350 kb window. All pairwise comparisons were statistically significant (BH-corrected T test, P < 0.05). (C) Proportion of gene pairs within compartments (insulation score ≥ 0.2) and located across boundaries (insulation score ≤ −0.2) insulation scores for each interaction group. (D) Proportion of gene pairs within compartments and located across boundaries separated by their ancestral status in *P. maximus*. (E) Genomic distances in *E. scolopes* between gene pairs grouped by ancestral chromosome status in *P. maximus* and interaction status. (F) Proportion of gene pairs within compartments and located across boundaries across ancestral *P. maximus* genomic distance categories. (G) Genomic distances in *E. scolopes* between gene pairs grouped by interaction status. (H) Schematics illustrating typical distances and compartment configurations for gene pairs interacting across coleoids versus those not in conserved coleoid interactions, given their ancestral*P. maximus* chromosome status, as suggested by the patterns observed in panel E (top) and G (bottom). Boxplot significance values are based on BH-corrected Wilcoxon tests: NS = non significant, * = P *<* 0.05, ** = P *<* 0.005, *** = P *<* 0.0005.

When examining the average insulation score between gene pairs (Fig. 3B for *E. scolopes*; for *S. officinalis* and *O. bimaculoides*, see Fig. S5), we found that the vast majority of pairwise comparisons between interaction categories were significantly different (BH-corrected T test, P < 0.05). Gene pairs with no conserved interaction in the coleoids frequently had an insulation score of zero or just below, with little variation around these values. This suggests that these regions lack a distinct topological structure. In contrast, interacting gene pairs exhibited positively skewed insulation scores, indicating that they are often located within chromatin compartments. However, the broader range of scores, including negative values, suggests that some interacting gene pairs overlap with compartment boundaries. Consistent with this, when we examined the proportion of gene pairs with high or low insulation scores across species and interaction categories, we found that the vast majority of gene pairs with no conserved interaction in the coleoids were associated with insulation scores ≤ 0.2, whereas interacting gene pairs were much more likely to be found in regions with higher insulation (≥ 0.2; Fig. 3C). Furthermore, where insulation boundaries are absent, intra-compartment mixing is stronger. Consistent with this, mixing entropy (Supplementary Note 1) is higher for conserved interacting gene pairs (Supplementary Note 1), indicating greater mixing within compartments and fewer cross-boundary contacts.

In the two coleoid species with available gene annotations, *E. scolopes* and *O. bimaculoides*, interacting gene pairs consistently showed higher gene coverage between them than pairs not in conserved coleoid interactions. For example, the median gene coverage within gene pairs not in conserved coleoid interactions was 33.0% in *E. scolopes* and 33.1% in *O. bimaculoides*, significantly higher than other interacting pair categories, ranging from 37.7% to 46.2% (Table S5). This enrichment suggests that interacting genes are more likely to reside in gene-dense regions, which may facilitate coordinated regulation, and further implies that these interactions are under stronger evolutionary constraint, likely reflecting their functional importance. Additionally, genomic regions between interacting pairs showed depletion of long interspersed repeats in all species, and have higher proportion of simple repeats, suggesting constraint against transposon-driven genomic expansion (Fig. S6).

### Chromosomal origin and genomic rearrangement drive divergence in gene pair proximity and regulatory architecture

To further investigate the relationship between chromatin interactions and the ancestral coleoid genome reorganisation, we examined whether interacting gene pairs were associated with chromosome rearrangements by assessing the chromosomal locations of the orthologous gene pairs in *P. maximus.* Patterns of chromosomal origin were consistent between *P. maximus* and *Nautilus pompilius* (Fig. S7), supporting the use of *P. maximus* as an appropriate outgroup for capturing the genomic changes that resulted from the genome rearrangement event after the coleoids diverged from the nautiloids. This analysis revealed that ancestral chromosomal origin further shaped expression patterns (Fig. S3D-F, Table S6, Supplementary Note 2), co-expression (Supplementary Note 2) and functional enrichment (Fig. S8, Supplementary Note 3) for interacting gene pairs, although there was no significant difference in gene coverage or tissue specificity (Supplementary Note 2). Furthermore, we found that gene pairs not in conserved coleoid interactions (Fig. S9A), as well as those that had lower insulation scores between them (Fig. 3D, S10A) were more likely to originate from different *P. maximus* chromosomes than interacting gene pairs and those with higher insulation scores. This is consistent with the notion that compartment boundaries frequently coincide with synteny breakpoints, regions of the chromosome that are more likely to break and reorganise^14–17^.

Across interaction categories, when examining genomic distance between gene pairs, we found that gene pairs interacting across the coleoids, despite large-scale genomic changes, tend to remain in close spatial proximity (Fig. 3E, G, S11, Table S7). In contrast, lineage-specific or species-specific interactions often form longer distance interactions, potentially resulting from compartment fusions during genome rearrangement or from regions more prone to genome expansion due to somewhat relaxed regulatory constraints. As expected based on their lack of interaction, gene pairs not in conserved coleoid interactions are consistently positioned over very large genomic distances.

Within each interaction category (interacting across coleoids, in Decapodiformes only, in *O. bimaculoides* only), the genomic distance between gene pairs varied significantly, depending on whether the orthologous genes were located on the same or different *P. maximus* chromosomes (Fig. 3E, S11A, Table S7A). In all species, interacting gene pairs that were located on the same *P. maximus* chromosome were significantly closer together than those located on different chromosomes. This suggests that interactions between genes that have remained syntenic since the ancestral molluscan genome are under stronger topological constraints, maintaining their proximity despite large-scale genome rearrangements. Alternatively, or in addition, interacting gene pairs may become incorporated into larger regulatory domains when brought onto the same chromosome through chromosomal fusions, which, after mixing, can result in entangled regulatory states, thereby favouring their maintenance over deep evolutionary timescales. Interestingly, this pattern was also significant in the same direction for gene pairs not in conserved coleoid interactions, though the effect was less pronounced. This may reflect the fact that gene pairs not in conserved coleoid interactions that originate from different *P. maximus* chromosomes have had less time for intrachromosomal mixing following genome rearrangement, leading to a residual signal of their ancestral chromosomal organisation. An exception to this pattern was observed for decapodiform gene pair distances in decapodiform-specific interactions, which showed similar genomic distances regardless of their ancestral chromosomal origin. As these interactions are likely lineage-specific, this pattern may reflect the larger interaction distances in the more expanded genomes of this clade, or, if they are ancestral coleoid interactions that have been lost in *O. bimaculoides*, moderately relaxed constraints on the regulatory linkages of these gene pairs. Together, our results suggest that the majority of regulatory interactions and topological compartments at sub-chromosomal levels were maintained following genome rearrangements in the coleoids, whereas many gene pairs without conserved coleoid interactions remained unlinked in their regulation once they were transferred to the same chromosome.

Generally, longer distances between intrachromosomal gene pair orthologs in *P. maximus* serve as a proxy for a higher likelihood that coleoid gene pairs were unlinked in their regulation prior to the ancient chromosomal rearrangement event (Fig. 1C). Topological organisation of coleoid gene pairs across different *P. maximus* distance categories were mostly evenly distributed (Fig. 3F, S9B, Table S7B). However, gene pairs interacting across the coleoids were significantly closer together across all profiled coleoid species if their orthologs in *P. maximus* were ≤5 Mb apart (Fig. 3G, S11B). Conversely, beyond this threshold, there were no significant differences in genomic distance between gene pairs interacting across the coleoids. This suggests that while proximity within 5 Mb may increase the likelihood of maintaining a regulatory interaction,interactions occurring at greater distances in the pre-rearrangement species do not appear to impact the conservation of spatial proximity. Consequently, coleoid-interacting gene pairs located more than 5 Mb apart in *P. maximus* are likely to represent post-rearrangement linkages rather than ancestral regulatory interactions. For other interaction categories, there were no significant differences in genomic distance between gene pairs in any coleoid species across *P. maximus* distance categories, with the exception of *E. scolopes* gene pairs interacting only in *O. bimaculoides*. Furthermore, there were no significant differences in the genomic distances between gene pairs not in conserved coleoid interactions across *P. maximus* distance groups. This suggests that, regardless of how closely genes were linked in *P. maximus*, gene pairs without conserved interactions are equally affected by genome rearrangement and expansion, likely because their linkages are not functionally important for gene regulation. Indeed, median genomic distances for these gene pairs were remarkably similar across *P. maximus* distance groups (Table S7B), implying that they may evolve under consistently relaxed regulatory constraints. Consequently, the subtle but significant differences observed across categories of interacting gene pairs and ancestral chromosome organisation likely reflect meaningful, divergent evolutionary processes rather than random variation.

Together, our results highlight different evolutionary scenarios of compartment evolution (as measured by interacting gene pair distributions), through genome rearrangement and expansion (summarised in Fig. 3H). These models provide a conceptual framework for understanding how ancestral genomic location and evolutionary changes in genome architecture influence the persistent spatial organisation.

### Chromatin loops represent evolutionarily dynamic structures associated with regulatory element enrichment

To investigate the role of genome topology in forming specific functional interactions, we assessed chromatin loop complement across coleoid species. In whole-embryo samples we identified 329, 912, and 663 chromatin loops in *E. scolopes*, *S. officinalis*, and *O. bimaculoides* respectively, based on combined 50 kb and 100 kb loop calls. After quantifying loops with genes at both anchors and removing duplicates containing identical gene sets, 156 gene-associated loops in *E. scolopes*, 402 in *S. officinalis*, and 388 in *O. bimaculoides* remained. This indicates that a large proportion of loops lacked genes at one or both ends, potentially reflecting the presence of cis-regulatory elements (CREs) such as enhancers, silencers, transcriptional repressors, or insulators, which may mediate both activation and repression of gene expression through chromatin looping^36–40^. Genes at loop anchors were found to be enriched for dynamic regulatory functions such as responses to stimuli and hormone signalling (Supplementary Note 4, Fig. S12), as well as cadherin genes (Supplementary Note 4, Fig. S13).

Across species, only a minority of gene-associated loops overlapped with gene pairs from the four interaction categories described above. Approximately 5% of loops contained gene pairs in conserved interactions (i.e. those interacting across the coleoids, or restricted to interacting across the Decapodiformes or in *O. bimaculoides,* combined) while ∼ 1% overlapped with gene pairs not in conserved interactions. Because the four gene pair categories were restricted to intrachromosomal orthologs shared across all three coleoid species and the outgroup *P. maximus*, they represent evolutionarily conserved interaction potential. This suggests that most chromatin loops are not formed around deeply conserved genes, but may instead reflect novel regulatory configurations involving genes that are scattered across different chromosomes in other species or recently evolved genes.

Loops containing genes, detected at 50 kb and 100 kb resolutions, were considered conserved across species if they included at least one orthologous gene overlapping loop each anchor, irrespective of gene orientation. Across *E. scolopes*, *S. officinalis* and *O. bimaculoides*, we found just three loops conserved across species (Fig. S14). We identified 27 conserved loops shared exclusively between *E. scolopes* and *S. officinalis* (Fig. S15), 6 between *E. scolopes* and *O. bimaculoides* (Fig. S16), and 8 between *S. officinalis* and *O. bimaculoides*. This small proportion of conserved loops suggests a high degree of loop turnover. The higher number of conserved loops shared between *E. scolopes* and *S. officinalis* is consistent with their more recent common ancestry compared to the deeper divergence between decapodiform species and *O. bimaculoides* (Fig. 1A). Interestingly, it is not only the presence of conserved loops that is maintained across species, but also their internal structural features. For example, we observe conservation of complex loop architectures such as ’double’ loops (e.g. Fig. S14C, S16F), ’volcano’-shaped interaction patterns^41^ (e.g. Fig. S15A, B, S16A, B), and ‘double compartment’-like structures within loops (e.g. Fig. S14B). This suggests that even small, subtle differences in local chromatin topology can be functionally relevant and under evolutionary constraint.

Genes located in conserved chromatin loops exhibited distinct expression patterns across *E. scolopes* tissues, with clustering revealing both broadly and tissue-specifically expressed genes (Fig. S17). Genes in loops conserved across all three species were consistently highly expressed in neural tissues and in the white body, a cephalopod-specific hematopoietic and immune organ^42^. This suggests that these deeply conserved loops may regulate genes involved in cephalopod-specific innovations. Genes in loops conserved between *E. scolopes* and either *O. bimaculoides* or *S. officinalis* showed more variable patterns, potentially reflecting lineage-specific roles.

Motif enrichment analysis of loop anchor regions in whole-embryo samples across species revealed both shared and species-specific patterns (Tables S8-S10). All three species showed strong enrichment for low-complexity and repeat-associated motifs, including poly-A/T tracts and GA/CA-rich sequences, likely reflecting the high abundance of LINEs in *S. officinalis* and *E. scolopes*, and SINEs in *O. bimaculoides*. Across species, motifs associated with key regulatory families such as Homeobox (Tgif1/2, Nkx3.1), Zinc finger (Maz, ZNF148), bHLH (SCL, Tcf12), Forkhead (Foxa2), and MYB were significantly enriched (hypergeometric test, BH-adjusted P < 0.05), suggesting a role for transcriptional control at loop anchors. CTCF-like motifs were also detected in all three species, supporting the idea that loop extrusion may be a conserved feature of cephalopod 3D genome organisation (as proposed in^8^).

We also tested whether CNEs were enriched within loop anchor regions across whole-embryo samples of the three coleoid species (Supplementary Note 5). Across all species, we observed consistent enrichment of CNEs in loop anchors compared to background regions. These findings indicate that loop anchors, where regulatory contacts are often established, are highly enriched for evolutionarily conserved putative regulatory elements. This suggests that although chromatin loops are evolutionarily dynamic and frequently rewired across species, they tend to recruit deeply conserved regulatory elements, repurposing ancestral genomic features to drive lineage-specific patterns of gene regulation.

Loops also showed significantly lower gene coverage between them than genome-wide gene pairs (Supplementary Note 5). This may reflect their roles as dynamic regulatory elements subject to fewer constraints, allowing them to emerge in gene-sparse regions that are more permissive to structural change. Gene sparsity may also facilitate the accumulation of repetitive elements between anchors, contributing to the low conservation of loops across species.

In all three *E. scolopes* developmental stages with available ATAC-seq data, loop anchor regions showed significantly greater chromatin accessibility than background regions (Supplementary Note 5). These findings are consistent with previous studies and indicate that loop anchors represent focal points of elevated chromatin accessibility, supporting the hypothesis that loops function as organisational hubs for transcriptional control^19,43^. Together with our GO, CNE, and motif enrichment results, this supports a model in which loop anchors are functionally enriched loci that contribute to dynamic gene regulation patterns.

### Chromatin loops show distinct scaling properties depending on their evolutionary history

To test whether conserved chromatin loop size scales with genome size across species, we calculated the linear regression slope between genome size and loop size for each loop conserved in at least two species. A Wilcoxon test on the per-loop slopes showed a statistically significant positive median slope (P = 0.014), indicating that loop size tends to increase linearly with genome size. These results suggest a general trend of loop length scaling with genome size among conserved loops (Fig. S18), with the exception of a few cases. This is consistent with these loops being ancestral coleoid structures affected by but retained during species-specific genome expansions.

Next, we compared all chromatin loop sizes across whole-embryo samples of *E. scolopes*, *S. officinalis*, and *O. bimaculoides*, expanding the analysis beyond only conserved loops to include all loops with at least one gene at each of their anchor points. Unlike the patterns we observed for the conserved loops, despite having the smallest genome, *O. bimaculoides* exhibited the largest and most variable loop sizes, while *E. scolopes* and *S. officinalis* showed consistently smaller loops with narrower distributions (Fig. 4A). Median loop sizes were smallest in *S. officinalis*, followed by *E. scolopes*, with *O. bimaculoides* having the largest median (medians = 1.3 Mb, 1.6 Mb, 2.9 Mb respectively). All pairwise comparisons were statistically significant. This pattern suggests that loop size is negatively correlated with genome size, and may reflect species-specific differences in genome rearrangement rather than overall genome expansion. In particular, the large loops in *O. bimaculoides* may result from its extensive history of chromosomal fusions, representing entangled regulatory states, potentially requiring longer-range contacts between regions that were previously on separate chromosomes. This is supported by the previously outlined comparative genomics and mixing analyses. Interestingly, this pattern parallels the fusion-followed-by-mixing scenario we observed for gene pairs from different *P. maximus* chromosomes, where residual signals of longer intergenic distances reflect their ancestral chromosomal organisation, but here occurring at a shorter evolutionary timescale.

**Figure 4.**
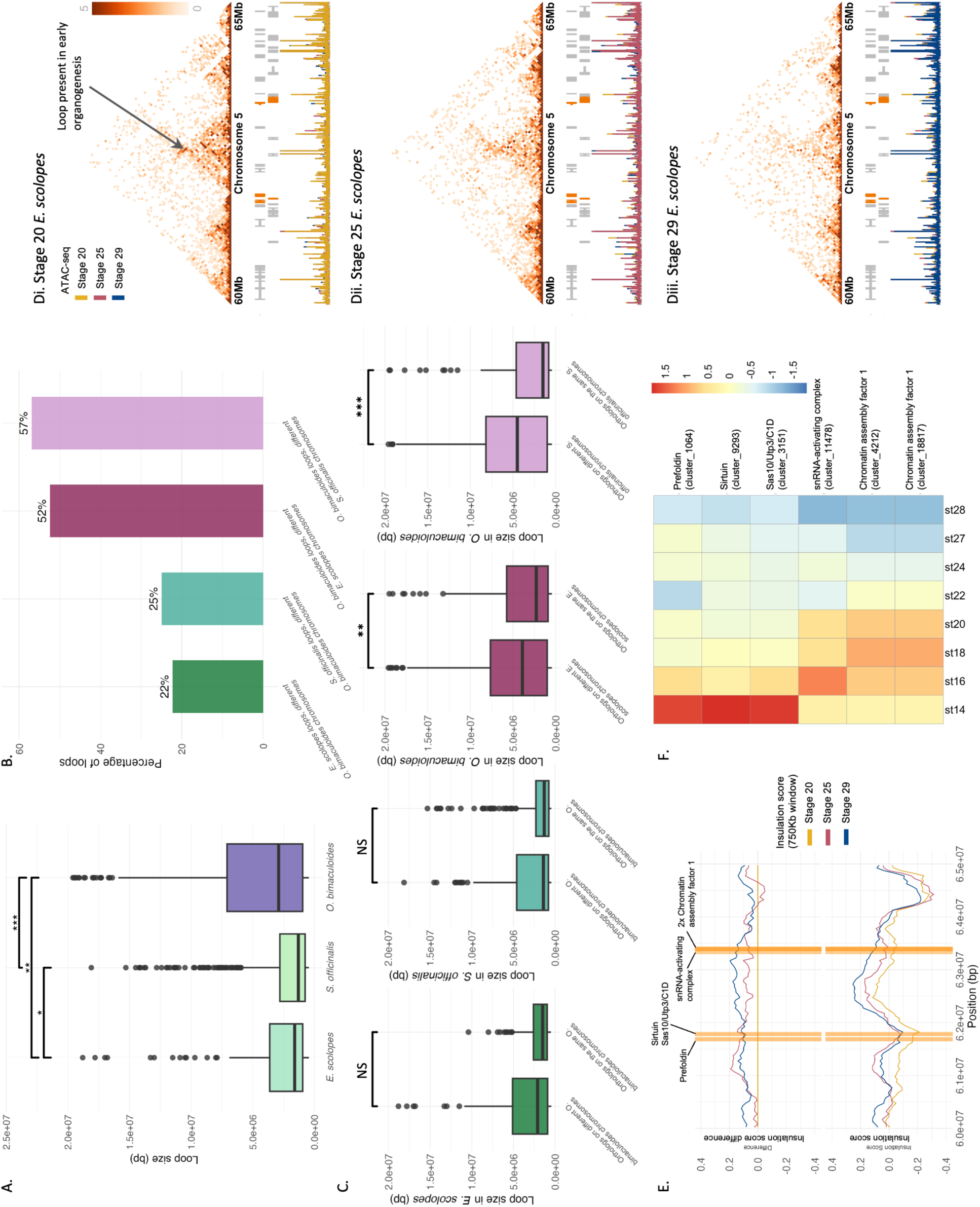
Chromatin loop configuration is shaped by lineage-specific chromosomal rearrangements and context-specific regulation. (A) Boxplots of loop sizes across species for loops containing genes at loop anchors detected in *E. scolopes* whole-embryo samples. Boxplot significance values are based on BH-corrected Wilcoxon tests: * = P *<* 0.05, ** = P *<* 0.005, *** = P *<* 0.0005. (B) Comparison of loop sizes in *E. scolopes* and *O. bimaculoides* based on whether or not genes at loop anchors contain orthologs on different chromosomes in the other coleoid genome. (C) Percentage of loop anchor genes in *E. scolopes, S. officinalis* and *O. bimaculoides* with orthologous genes located on different chromosomes in the other coleoid species’ genome. Boxplot significance values are based on BH corrected Wilcoxon tests: NS = non significant, * = P *<* 0.05, ** = P *<* 0.005, *** = P *<* 0.0005. (D) Micro-C contact maps for a 5Mb region of chromosome 5 for *E. scolopes* across three developmental stages: (Di) stage 20 (early organogenesis), (Dii) stage 25, and (Diii) stage 29 (pre-hatching). A significant reduction in loop strength was observed between stage 20 and 25 (FDR = 0.0032) and between stage 20 and 29 (FDR = 0.0094), as identified by Mustache’s differential loop calling approach. The loop present at stage 20 is indicated, and is missing at the later developmental stages. Normalised ATAC-seq (from^8^) signal tracks show chromatin accessibility at each stage. (E) Insulation scores and the difference in insulation scores around conserved loop anchors across developmental stages in *E. scolopes*. Line plots show average insulation scores at a 750 kb window across stage 20, 25, and 29. Highlighted regions correspond to genes at loop anchors. ‘2x Chromatin assembly factor 1’ indicates 2 genes with the same name in that location. (F) Heatmap showing gene expression patterns of genes located at conserved loop anchors across 8 developmental stages in *E. scolopes*. Loop anchor genes include Prefoldin, Sirtuin, Sas10/Utp3/C1D, snRNA-activating complex (SNAPc), and two subunits of Chromatin Assembly Factor 1 (CAF-1).

To explore whether this increase in loop size is indeed linked to changes in gene organisation, we examined the chromosomal origin of genes located at loop anchors. Specifically, we assessed whether loop-associated genes were located on the same or multiple different chromosomes in the other coleoid species and in *P. maximus*. A substantial proportion of loops in all species contained genes whose orthologs were located on different chromosomes in the other species, but this was particularly pronounced in *O. bimaculoides*, where over 50% of loops contained genes from different decapodiform chromosomes (Fig. 4B), and 70% from different *P. maximus* chromosomes (Fig. S19). These results align with our observations and corroborate previous findings on widespread chromosomal fusions in octopuses^33^. Furthermore, this contrasts with our earlier finding that ∼ 40% of conserved interacting gene pairs spanning different *P. maximus* chromosomes (Fig. S9), suggesting that chromosomal fusions more strongly shape loop formation than conserved gene pair interactions.

Additionally, we compared gene-associated loop sizes in each species based on whether the genes at loop anchors were located on the same or multiple different chromosomes in the other coleoid species. In all three species, loops connecting genes from different chromosomes in the other coleoid species tended to be larger than those connecting genes from the same chromosome (Fig. 4C). This trend was especially marked in *O. bimaculoides*, where the difference was statistically significant, consistent with the idea that extensive chromosomal fusions in the octopodiform genomes have created longer-range regulatory contacts. In *E. scolopes* and *S. officinalis*, the difference in loop size between same- and different-chromosome gene pairs was less pronounced and not statistically significant, suggesting that larger loops are most strongly associated with the extensively fused *O. bimaculoides* genome. Notably, loop size did not correlate with the intergenic distance between orthologous gene pairs on the same chromosome in the other species (R² < 0.01 across all comparisons) indicating that the increased size of intrachromosomal loops is not simply due to ancestral gene spacing but is likely a direct consequence of chromosomal rearrangement events. Together, these results further support the dynamic nature and more recent evolutionary origin of chromatin loops in the coleoids.

### Chromatin loop formation is highly context-specific

Since chromatin loop conservation across species is minimal, we next examined both the full set of detected loops (Fig. S20) and those specifically associated with genes (Fig. S21) across developmental stages and tissues. Chromatin loop abundance varied substantially in both contexts. A representative example of a significantly differential chromatin loop and its associated expression patterns across development is shown in Fig. 4D–F, with additional details provided in Supplementary Note 6. In *E. scolopes*, stage 25 embryos exhibited substantially more loops than either stage 20 or stage 29 (Fig. S20Ai, S21Ai). As stage 25 marks the final stages of organ differentiation and functional maturation during late organogenesis^44^, this elevated loop abundance may reflect the widespread changes in gene expression occurring at this stage, as well as the regulatory complexity needed to establish stable, mature expression programs. In *O. bimaculoides*, neural tissues, i.e. the arm crown, eyes and optic lobes, and the central brain, showed similar numbers of loops, while internal organs had about one third as many (Fig. S20Bi, S21Bi), consistent with the heightened regulatory demands of neural tissues^31^. In contrast, *S. officinalis* showed a distinct pattern; the eyes had by far the highest number of loops, followed by the optic lobes, with the arm crown, central brain, and internal organs exhibiting substantially fewer (Fig. S20Ci, S21Ci). These contrasting tissue-specific patterns between *O. bimaculoides* and *S. officinalis* suggest that the regulatory role of chromatin loops may be distributed differently across neural and peripheral tissues in these species, possibly reflecting lineage-specific adaptations in gene regulation or differences in tissue complexity and function. Supporting this interpretation, in *S. officinalis*, retinal determination genes are expressed specifically in the eye and optic lobe during embryogenesis, linked to the formation of local regulatory architectures driving neural complexity^45^. Furthermore, the cuttlefish optic lobe continues to mature into adulthood^46^, paralleling ongoing increases in regulatory and circuit complexity.

Pairwise comparisons across all developmental stage (Fig. S20Aii, S21Aii) and tissue (Fig. S20Bii, S20Cii, S21Bii, S21Cii) combinations revealed no statistically significant overlap or depletion in loop usage, regardless of whether loops contained genes (super exact test, BH-adjusted P > 0.05). This underscores the dynamic and context-dependent nature of chromatin loop architecture, further supported by the distinct GO and motif enrichment patterns observed across tissue-specific samples (Fig. S22, S23, S24, Supplementary Note 4).

Together, these findings suggest that chromatin looping is not only poorly conserved across evolutionary time but is also highly modular within species, with distinct regulatory interactions likely tailored to the specific transcriptional needs of developmental stages or individual tissues (Fig. 4A-C, Supplementary Note 6). This context-specific organisation may enable precise temporal and spatial control of gene expression during complex processes such as organogenesis and tissue-specific functions, and may be facilitated by relaxed evolutionary constraints that allow greater plasticity in loop formation and usage. This is in contrast to the more stable compartments as measured by the interacting gene pair approach or TAD predictors (Supplementary Note 5, Fig. S25).

To gain deeper insight into how context-dependent loop formation relates to transcription, we examined expression patterns of genes within developmental stage-specific loops in *E. scolopes*. Genes in stage-specific loops tend to be more highly expressed at their corresponding developmental stage and exhibit lower expression at other stages (Fig. S26A). However, these overall trends are not always statistically significant, suggesting more complex regulatory dynamics. To investigate this further, we visualised gene expression within individual loops using heatmaps (selected example in Fig. 4F, Supplementary Note 6, heatmaps across all developmental stage-specific loops in Fig. S26B). These heatmaps reveal diverse expression patterns across developmental stages. In some cases, loops are associated with coordinated upregulation of genes at a specific stage, while in others, loops correspond to downregulation. Most frequently, we observe mixed patterns, with some genes upregulated and others downregulated within the same loop. Consistent with this, genes within stage-specific loops exhibited low co-expression across developmental RNA-seq data, with median pairwise Pearson’s correlation coefficients of 0.0441, 0.203, and 0.200 for *E. scolopes* stages 20, 25, and 29, respectively. Therefore overall, context-specific loop-associated genes exhibit complex expression dynamics, suggesting an interplay between diverse regulatory elements, such as enhancers and repressors, operating within the same 3D chromatin context.

### Knockout of putative regulatory sequence reveals long-range meta-loop-like interactions

Our genome-wide analyses indicated that although most chromatin loops are evolutionarily dynamic and highly context-specific, a small subset are deeply conserved across coleoid whole-embryo samples. Furthermore, loops are enriched for accessible chromatin, CNEs, TF binding motifs, and expression patterns associated with neural gene regulation, suggesting potential roles in orchestrating transcriptional programs essential for nervous system development. To functionally test whether conserved loops harbour regulatory elements critical for gene expression, we used CRISPR-Cas9 to disrupt a candidate regulatory region within one such conserved loop in *Euprymna berryi*, a close relative of *E. scolopes* with established genetic tools.

To this end, we targeted a ∼1 kb region within an intron of the gene Quaking B, located in a conserved coleoid chromatin loop (Fig. 5A), which contains genes upregulated across neural tissues and developmental stages (Fig. S27B, C) but is extensively rearranged in ancestral molluscs, suggesting a derived regulatory architecture in coleoids. Quaking B was chosen as the host gene to identify a putative intronic regulatory sequence due to the availability of sufficiently long non-repetitive non-coding regions within it and evidence for its role during neural development^47–49^. Furthermore, the targeted intronic region is conserved among Decapodiformes and also has a conserved region shared with *bimaculoides* located ∼ 150 bp upstream (Fig. 5B). The repetitive nature of coleoid genomes make it very difficult to target specific sites and the three chosen guides (Fig. 5B) were the only possible knock-out scenario without off-targets. The deletion was confirmed by PCR (Fig. S28A, B) and genotyping. The deleted region includes multiple predicted transcription factor (TF) binding motifs, several of which are associated with neural development and gene regulation (Supplementary Note 7, Table S11).

**Figure 5.**
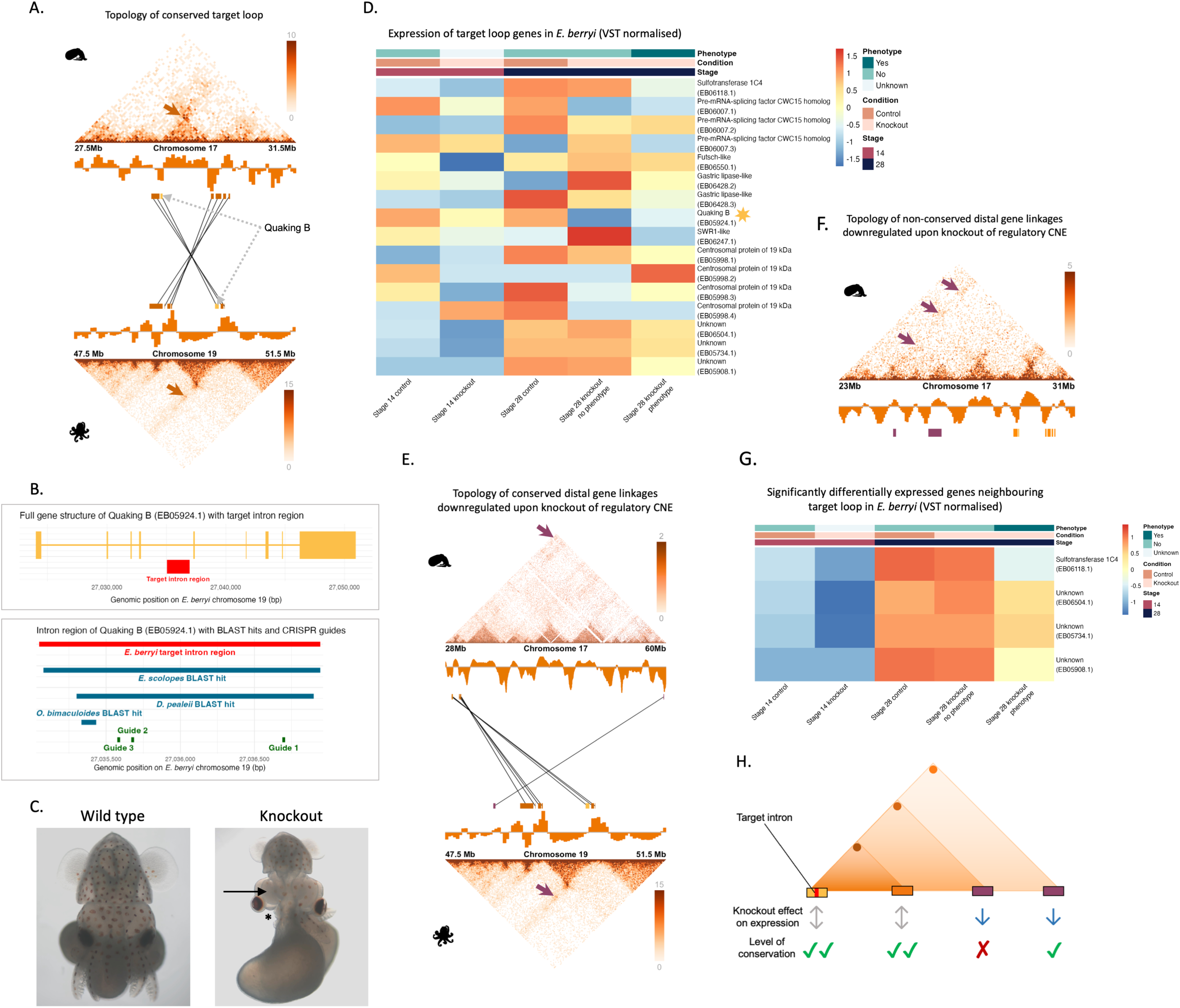
CRISPR knockout of a putative regulatory region reveals long-range meta-loop-like interactions and transcriptional effects in coleoid embryos. (A) Micro-C contact matrices showing the conserved chromatin loop between *E. scolopes* (top) and *O. bimaculoides* (bottom). The gene (Quaking B) containing the target intron is highlighted in yellow and indicated with a grey dotted arrow, with other genes within the loop shown in orange. Black lines indicate orthology between loop genes across species. Contact maps are shown at 50 kb resolution for *E. scolopes* and 25 kb for *O. bimaculoides*. To improve clarity, background genes were removed from the gene annotations overlaid on the contact maps. Gene colouring, orthology, and resolution are consistent with panels D and E, where applicable. Note that the crossed orthology lines here and in B reflect opposite assembly orientations between the two species rather than a true inversion; gene order within this region is colinear. (B) Schematic representing the CRISPR targeting of the intron region in Quaking B. The top panel depicts the gene structure of Quaking B (EB05924.1) in yellow and the intron region targeted by CRISPR highlighted in red in *E. berryi*. The bottom panel shows a detailed view of the targeted *E. berryi* intron region, showing conserved BLAST hits from *E. scolopes*, *O. bimaculoides*, and *D. pealeii* aligned to the *E. berryi* genome. The red bar indicates the *E. berryi* target intron region, blue bars show orthologous BLAST hits in other coleoid species, and green boxes mark the CRISPR guide sites. The knockout region spans from guide 1 to guide 3. (C) Representative images of control (left) and knockout (right) *E. berryi* embryos at stage 28, showing morphological defects, including smaller optic lobes (arrow) and smaller or missing arms (star), following deletion of the target regulatory region. (D) Heatmap of expression levels (VST-normalised) for all transcripts of genes located within the conserved loop across control and knockout *E. berryi* embryos at stages 14 and 29. Quaking B, the gene with the target intron, is indicated by a yellow star. Despite shared topology, gene expression was variable and not significantly altered after BH correction (Wald test). Genes from top to bottom of the heatmap correspond to those from downstream to upstream of the loop in the *E. berryi*, but right to left along the loop in the *E. scolopes* contact map shown in (A). Transcripts with zero variance in expression were removed from the heatmap. (E) Contact matrices and topology of distal interactions (purple arrows) conserved across *E. scolopes* and *O. bimaculoides* that were significantly downregulated upon knockout in *E. berryi*. Significantly downregulated genes are depicted in purple. (F) Contact map of a region *E. scolopes* showing topological positioning of downstream downregulated genes in *E. berryi* knockout, most of which lack strong BLAST hits and are unannotated in *O. bimaculoides*, suggesting they are clade-specific novel genes. Downregulated genes are represented in purple and their topological linkage to the target intron is indicated by purple arrows. Three genes are depicted, though it appears as only two because the two on the right overlap visually. (G) Heatmap of (VST normalised) expression of significantly (Wald test with BH correction) downregulated genes downstream of the target intron in knockout embryos, showing consistent downregulation at stage 29. Sulfotransferase 1C4 is the downregulated gene depicted in purple in (D) and the three unknown genes are those shown in purple in (E). (H) Schematic summary of knockout results and conservation status across coleoids. This model illustrates how conserved regulatory introns can regulate distal genes through long-range chromatin looping, and how these loops, despite being conserved, may undergo topological mixing of their contacts over evolutionary time, resulting in the rewiring of regulatory networks. Grey arrows represent variable changes in gene expression following knockout of the regulatory element, while blue arrows indicate significant downregulation. Green ticks denote conserved interactions in both gene order and contact size; the single green arrow shows partial conservation (i.e., conserved interaction but with differences in region size or gene order); the red cross marks an interaction not conserved in other coleoid lineages.

Among the 25 CRISPR embryos used for RNA-sequencing, all showed signs of developmental delay. Five embryos exhibited a clear reduction in brain size, particularly in the optic lobe. An additional five were severely malformed with disrupted body structures and reduced neural tissue, while six appeared smaller overall with notably shorter arms. Nine did not show an overt phenotype. Thus, depending on classification criteria, approximately one third to two thirds of embryos displayed brain-specific morphological defects, consistent with developmental impairment of neural structures (Fig. 5C, S28C, D).

To assess transcriptional consequences, we conducted whole-body RNA-seq at stages 14 and 28. Stage 14 was selected as Quaking B, the target gene, exhibits peak expression at this developmental stage (Fig. S27B). PCA plots (Fig. S29) showed separation between control and knockout samples based on gene expression at both stage 14 and stage 28. At stage 28, samples with a recorded phenotype cluster separately from those without, suggesting a stronger transcriptional response associated with the observed phenotype. Although genes within the same compartment and loop exhibited expression variability (Fig. 5C), no statistically significant changes were detected. The most pronounced downregulation within the compartment was observed in the host gene Quaking B itself at stage 14 and stage 28, although this was not significant after BH correction.

Strikingly, transcriptional effects were also observed at long range. Upstream of the target site, a single significantly downregulated gene conserved in *O. bimaculoides* was identified, located 14.5 Mb away in *E. berryi* and 30.5 Mb in *E. scolopes.* Although these genes are more distant in *E. scolopes*, they are positioned much closer to the core conserved compartment in *O. bimaculoides* (1.6 Mb) and they appear to be topologically linked in all three species (Fig. 5D, Fig. S30). Downstream of the *E. berryi* target, the nearest significantly downregulated genes (Fig. 5F) were located 2.9-4.1 Mb away in *E. berryi* and 2.9-4.2 Mb in *E. scolopes.* In *O. bimaculoides*, these genes are unannotated or on different chromosomes to the target loop (Fig. 5E), and most lacked strong homology to other metazoans, indicating they are likely coleoid-specific. The significantly downregulated genes showed high expression in *E. scolopes* adult neural tissues, as well as the mantle and gills (Fig. S27C) and at late stages during *E. scolopes* embryonic development (Fig. S27D). The conserved loop in *E. berryi*, along with the regions containing genes downregulated following knockout of the target intron, are shown in Fig. S30 for both *E. berryi* and *S. officinalis*.

Altogether, these findings support a model in which the targeted non-coding region acts as a putative enhancer within a conserved chromatin loop. Its disruption caused both local and long-range transcriptional and developmental effects. This is consistent with a meta-loop architecture, in which regulatory elements exert influence across compartment boundaries and over large genomic distances^50^. The downregulated genes span both conserved and lineage-specific targets, suggesting evolutionary flexibility in regulatory wiring despite structural conservation of the loop. The more limited expression changes observed within the conserved loop compared to the stronger transcriptional effects on genes beyond the loop imply that genes embedded within highly conserved and, likely, regulatorily entangled loop topologies are more buffered against changes to single CNEs, whereas those involved in more distal and less conserved regulatory interactions show stronger transcriptional responses when regulatory regions are disrupted.

Figure 5H illustrates this scenario, showing how chromatin loops can mediate long-range regulatory interactions that are constrained and evolvable at the same time. These results reinforce the concept of partially conserved regulatory loops, where conserved topologies harbor regulatory elements that may adapt their gene targets over time while retaining core functions in neural development. Supporting this, the corresponding loop in *O. bimaculoides* shows tighter co-localisation between enhancer and promoter, suggesting additional regulatory entanglement following chromosomal fusion and mixing in octopods.

## Conclusion

How compartments and regulatory connections change throughout evolution has only been studied in a few model systems^50,51^. Recent work highlights diverse patterns of compartmentalisation and regulatory architecture across animal lineages^9,11,19^. Coleoid cephalopods offer a powerful model for exploring how macroevolutionary processes shape and maintain regulatory landscapes, owing to their ancient and extensively reorganised and expanded genomes and striking phenotypic novelties.

By analysing 3D genome evolution across three coleoid species in the context of their unique evolutionary history, our study provides insight into the origin, maintenance and plasticity of chromatin interactions and their associated regulatory patterns. We show that in the expanded genomes of Decapodiformes, the regulatory interactions have more ancestral signatures. In the less expanded *O. bimaculoides* genome, our data on syntenic comparisons, compartment evolutionary history and loop sizes supports a model of secondary chromosomal fusion-with-mixing, resulting in more ‘entangled’ regulatory configurations^22^ and a more derived topological state for Octopodiformes. Furthermore, we show that while higher-order compartment structures are generally conserved, chromatin loops are dynamic and enriched in flexible regulatory features. Knockout of a putative regulatory site confirmed the functional relevance of long-range loop-mediated interactions with both conserved and novel genes.

Crucially, our results reveal a hierarchy of regulatory constraints across the 3D genome (summarised in Fig. S31), in which different classes of topological interactions, defined by their ancestral chromosomal state, clade specificity and 3D structure, show distinct levels of evolvability. This variation in constraint influences how genome rearrangement and expansion affect different parts of the genome. Rather than uniformly reshaping regulatory architecture, these processes selectively rewire, retain, or eliminate interactions based on their underlying level of constraint. Furthermore, mixing of multiple regulatory interactions can lead to entangled regulatory configurations, such as the particularly pronounced pattern observed in *O. bimaculoides*, which, depending on their complexity, may become increasingly resistant to reorganisation^22,34^. This dynamic interplay between topological flexibility and constraint gives rise to a distinct regulatory landscape in coleoids, shaped not only by the scale of genomic change, but also by the functional importance, regulatory context, and structural organisation of the affected regions.

Taken together, our findings highlight the importance of studying 3D genome evolution in coleoid lineages and show how genome architecture, differential evolutionary constraints, and large-scale chromosomal changes interact to drive regulatory innovation, thereby illustrating how spatial genome organisation both shapes and is shaped by evolution over deep timescales.

## Methods

### CNE analysis

Whole genome alignments were conducted using BLASTN v.2.16.0^52^ and using options -task megablast-perc_identity 0 -template_length 16 -penalty -2 -word_size 11 -evalue 1 -template_type coding_and_optimal^53^. To further filter the alignments open reading frame predictions were conducted using Transdecoder 5.7.1 (TransDecoder.LongOrfs -m10 parameter was used), only 50bp+ alignments were retained.

### Multi-locus topology (MLT) analysis Genome database

In total, 15 chromosome-scale cephalopod genome assemblies were used for comparisons. All genome assemblies are available via NCBI. There were 4 species within the Octopodiformes: *Eledone cirrhosa* (GCA_964016885.1), *Octopus bimaculoides* (GCA_001194135.2)^30^, *Octopus sinensis* (GCA_006345805.1)^54^, *Octopus vulgaris* (GCA_951406725.2)^55^. There were 10 species within the Decapodiformes: *Doryteuthis pealeii* (GCA_023376005.1), *Euprymna berryi*^56^, *Eumandya parva* (GCA_964059275.1), *Euprymna scolopes* (GCA_024364805.1)^30^, *Sepia lycidas* (GCA_963932145.1), *Sepia officinalis* (GCA_964300435.1), *Sepiola atlantica* (GCA_963556195.1), *Sepioteuthis lessoniana* (GCA_963585895.1), *Stigmatoteuthis arcturi* (GCA_964276865.1), *Thysanoteuthis rhombus* (GCA_963457665.1). There was 1 species within the Nautiloidea: *Argonauta hians* (GCA_123456789.1).

### Genome annotation

We annotated the genomes of the above species, except for the species with available annotations:

*E. scolopes, O. bimaculoides*, *Doryteuthis pealeii*, *Octopus sinensis* and *Octopus vulgaris*. Our strategy leveraged the existing cephalopod genome annotations as protein evidence during annotation. First, miniprot v.0.12-r237 with parameters --outn 10 --gff --trans was used to map the protein amino acid sequences from the following genome assemblies to the target genome: *Pecten maximus* (GCF_902652985.1), *Euprymna scolopes* (GCA_024364805.1), *Octopus vulgaris* (GCA_951406725.2)^55^, *Octopus bimaculoides* (GCA_001194135.2)^30^. The --trans option saves the protein sequence of the target genome to the .gff file. The script miniprot_gff_trans_to_proteinfa.py was used to extract protein fasta files from the .gff file. The resulting protein fasta sequences were clustered and reduced to representative sequences using a sequence identity threshold of 0.9 using cd-hit v.4.8.1^57^. The resulting fasta file and the miniprot gff file were used to generate filtered gff and chrom files using the script miniprot_filter_gff_from_fasta_gen_chrom.py.

### Ancestral ortholog identification

We inferred the identity and genome locations of the 2,361 bilaterian-cnidarian-sponge ancestral linkage group (BCnS ALG) orthologs^35^ in the cephalopods using the HMM-based approach using odp v0.3.0^58^.

### Multi-locus topology inference

We used the CNE and BCnS ALG genome location information as coordinates to generate similarity matrices as input for genome multi-locus topology matrices^34^. The following sets of loci were run through the MLT pipeline for the clades Coleoidea, Octopodiformes, Decapodiformes: (1) BCnS ALGs only, (2) O. bimaculoides CNEs only, (3) E. scolopes CNEs only, (4) BCnS ALGs + E. scolopes CNEs, (5) BCnS ALGs + O. bimaculoides CNEs. For the combined CNE + BCnS ALG MLT inference, the database files were simply concatenated to form a locus supermatrix. CNEs were defined as regions of at least 100 bp conserved between *E. scolopes* and *E. berryi* (for *E. scolopes* CNEs), and between *O. bimaculoides* and *O. vulgaris* (for *O. bimaculoides* CNEs). To be considered orthologous to the other species, each region also had to show at least 50 bp of sequence conservation.

### Micro-C sample collection and preparation

*E. scolopes* samples were provided by Schönbrunn Zoo in Vienna and dissected at the University of Vienna, Austria. *S. officinalis* specimens were obtained and dissected at the Laurent Lab at the Max Planck Institute for Brain Research in Frankfurt, Germany. *O. bimaculoides* specimens were collected and dissected at the Marine Biological Laboratory in Woods Hole, Massachusetts, and the Ragsdale Lab at the University of Chicago, Illinois, USA (Table S1). Following dissection, all samples were flash-frozen in liquid nitrogen and stored at -70°C.

### Micro-C library preparation and sequencing

For *E. scolopes*, eight pooled whole-embryo samples at developmental stages 20, 25, and 29 were used for Micro-C sequencing, with two replicates performed for the stage 29 samples. For *S. officinalis*, tissue-specific Micro-C was performed using pools of 10 or 15 embryos at stage 29, while whole-embryo Micro-C at the same stage was conducted on two separate samples: one from a single embryo and one from a pool of two embryos. For *O. bimaculoides*, three pooled samples at stage 20 were used for tissue-specific Micro-C sequencing, and a single stage 20 embryo was used for whole-embryo Micro-C. Furthermore, three pooled stage 29 embryos were used for Micro-C in *E. berryi*. In the Decapodiform species (*E. scolopes* and *S. officinalis*), stages 20 and 25 represent early and late organogenesis, respectively, while stage 29 corresponds to the final stage before hatching^44^. In the Octopodiform species (*O. bimaculoides*), stage 20 also represents the pre-hatching stage^59^. The tissue samples collected for *S. officinalis* included the central brain, optic lobes, eyes, internal organs, and arm crown (including tentacles). For *O. bimaculoides*, the sampled tissues were the central brain, eyes and optic lobes (combined), internal organs, and arm crown. Tissue-specific samples were all collected from the same individuals, except for the octopus arm crown, which was obtained from a different individual. Detailed sample information is provided in Table S1.

The Micro-C library was prepared using the Dovetail Micro-C Kit according to the manufacturer’s protocol. Briefly, the chromatin was fixed with disuccinimidyl glutarate (DSG) and formaldehyde in the nucleus. The cross-linked chromatin was then digested in situ with micrococcal nuclease (MNase).

Following digestion, the cells were lysed with SDS to extract the chromatin fragments and the chromatin fragments were bound to Chromatin Capture Beads. Next, the chromatin ends were repaired and ligated to a biotinylated bridge adapter followed by proximity ligation of adapter-containing ends. After proximity ligation, the crosslinks were reversed, the associated proteins were degraded, and the DNA was purified then converted into a sequencing library using Illumina-compatible adaptors. Biotin-containing fragments were isolated using streptavidin beads prior to PCR amplification. The libraries were sequenced on an Illumina platform at the Vienna BioCenter Core Facility, using the NovaSeq S4 XP and NovaSeqX 10B XP flow cells, both with PE150 read mode. One initial test sample, however, was sequenced on a NextSeq550 with PE75 medium read mode. The number of read pairs generated can be seen in Table S1.

### Processing and mapping of Micro-C data

Sequenced reads were trimmed to 50bp using Trimmomatic v.0.39^60^ to improve mapping rates and compensate for gaps in reference genome assemblies. For the downstream analyses focused on cross-species comparisons, reads for the two replicates of each of the Decapodiform species were concatenated to achieve approximately equal read coverage to that of *O. bimaculoides*, accounting for their larger genome sizes. Tissue-specific samples were downsampled to approximately 350M read pairs using seqtk v1.4 ^61^ if they fell above 5% of this number (Table S2). Reads were then mapped to the *E. scolopes* (GCA_024364805.1)^8^, *S. officinalis* (GCA_964300435.1)^62^, *O. bimaculoides* (GCA_001194135.2)^30^, and *E. berryi*^63^ genomes, softmasked genomes using Hic-Pro v.3.0.9^64^ using default parameters in the config file, with the exception of the ’Digestion Hi-C’ parameters, which were left blank. The number of mapped reads can be seen in Table S2. The allValidPairs files generated by HiC-Pro were filtered to retain only intrachromosomal interactions and exclude those involving scaffolds. The script hicpro2juicebox.sh from the Hic-Pro utilities was then used with JuicerTools v.1.22.01^65^ to convert the filtered allValidPairs file from the HiC-Pro output to a Juicer .hic file with KR normalisation. Micro-C contact maps were visualised in Juicebox v2.15^66^ for whole-chromosome square plots and in Plotgardener v1.8.3^67^ for smaller triangular plots.

### Ortholog classification and visualisation of syntenic topological regions

For *E. scolopes*, the gene annotation from Belcaid *et al*.^29^ was used and for *O. bimaculoides*, the NCBI annotation (GCF_001194135.2_ASM119413v2_genomic.gtf)^30^ was used in downstream analyses. The genome assemblies and gene annotations for the great scallop, *Pecten maximus* (xPecMax1.1, GCF_902652985.1)^68^ was retrieved from the NCBI RefSeq database. For the chambered nautilus, *Nautilus pompilius*, the genome assembly and gene annotation (GCA_047652355.1) from^69^ were used. Due to a lack of gene annotation for the *S. officinalis* genome assembly, *E. scolopes* proteins^29^ were mapped to the *S. officinalis* genome using Miniprot v0.12^70^ with default parameters and the -gff flag. The resulting GFF file was filtered to exclude scaffolds, retain only ’mRNA’ lines, and extract the highest-ranked hits (filtering for ’Rank=1’). This filtered dataset yielded 24,430 genes and was subsequently used as the *S. officinalis* gene annotation. One-to-one orthologs between other species were identified using BLASTP with the parameters -evalue 1E-2 -max_target_seqs 1 -outfmt 6, with *E. scolopes* proteins used as the reference database. Custom R scripts were used to generate dot plots and to produce the synteny overlays shown on triangle Micro-C contact maps.

### Classification of conserved and non-conserved intrachromosomal gene pairs and their ancestral states in P. maximus

Firstly, interchromosomal interactions were removed from .hic files, and KR-normalized sparse Micro-C matrices were then extracted for each chromosome using the dump command in JuicerTools, at 50 kb resolution for *O. bimaculoides* and 100 kb for the decapodiform species. A lower resolution was used to account for lower read coverage in the decapodiform species and to obtain the highest number of orthologous interactions across the three species. These were concatenated into one file for each genome and resolution, with the chromosome names for each interacting bin pair added to the first column. For each species, all genes located within or overlapping the start and end bins of every entry in the dumped sparse matrix were identified. If one or both bins had no genes in, they were excluded from downstream analyses. If multiple genes were present within a bin, all possible combinations of gene pairs were generated. Gene pairs where both genes were identical were removed. One-to-one orthologous gene pairs across all three coleoid species were identified, and their corresponding interaction frequencies were consolidated into a single file based on the gene pairs assigned in the *E. scolopes* 100 kb resolution matrix. The following gene pair analyses were performed only on genes with intrachromosomal orthologous pairs in the three coleoid species, thus having the potential to interact in all three coleoid species, as well as one-to-one orthologs in the outgroup species, *P. maximus*.

Gene pairs were classified as ’interacting’ if at least 10 KR-normalised reads were mapped between the bins they were associated with. To be considered ’interacting across the coleoids’, this threshold had to be met in all three species. Gene pairs were classified as ’interacting in Decapodiformes only’ if the threshold was met in *E. scolopes* and *S. officinalis*, but not in *O. bimaculoides*. Conversely, a gene pair was classified as ’interacting in *O. bimaculoide*s only’ if it met the threshold in *O. bimaculoides* but not in the other two species. Gene pairs classified as ’not in conserved coleoid interactions’ were genes in bin pairs that fell below this threshold in all coleoid species. The threshold of ≥ 10 KR-normalised mapped reads was chosen based on the observation that, across all species’ contact matrices, interactions between regions 10–20 Mb apart very rarely exceeded 9 mapped reads, while regions ≥ 20 Mb never reached this interaction frequency (Fig. S2). Below this threshold, interaction frequency appeared independent of genomic distance. For this reason, we considered ∼ 20 Mb the upper limit for topological interactions in the coleoids, and accordingly, highly interacting domains beyond this distance were usually not observed in contact maps. Within each interaction category, interaction frequencies were averaged for each gene pair that appeared multiple times in the dataset. This approach avoided compartment calling, which can be inconsistent across window sizes and algorithms, and instead provides a comparative framework for quantifying 3D synteny, defined here as the conservation of spatial gene proximity and chromatin interaction patterns across species, encompassing both multi-scale compartmentalisation and nested domain architecture.

Next, gene pairs were classified based on their ancestral chromosomal state in *P. maximus*, a scallop species with a high-quality reference genome and gene annotation. To validate its use as an outgroup, we confirmed that patterns of chromosomal origin were consistent between *P. maximus* and *N. pompilius*, supporting the suitability of *P. maximus* for identifying genomic changes associated with the large-scale genome rearrangement that occurred after their divergence from nautiloids. Firstly, gene pairs were grouped depending on whether the orthologs of each gene were located on the same chromosome or on different chromosomes in *P. maximus*, with the assumption that coleoid regulatory linkages between genes from different scallop chromosomes represent regulatory patterns that emerged after the ancient coleoid genome rearrangement event. If gene pairs were located on the same *P. maximus* chromosome, they were further classified into one of three distance categories: ≤5 Mb, >5 Mb and ≤15 Mb, or >15 Mb, with shorter distances being a proxy for a higher likelihood that genes were linked in their regulation in the ancestral mollusc species. Genomic distance between gene pairs was calculated as the start position of the downstream gene minus the end position of the upstream gene. Pairwise Wilcoxon rank-sum tests with Benjamini–Hochberg (BH) correction for multiple comparisons were used to test for significant differences between gene pair groups. Data processing and filtering in this and all subsequent sections were carried out using custom Python v.3.10.4 scripts and in R v.4.3.1^71^ with the packages dplyr v.1.1.4^72^, tidyr v.1.3.1^73^, and stringr v.1.5.1^74^. Statistical tests were also performed in R, and density plots, box plots, scatter plots, and bar plots were generated using ggplot2 v.3.5^75^ here and throughout.

### Gene expression analyses in E. scolopes

Publicly available TPM-normalised gene expression data for *E. scolopes* tissues^76^ and developmental stages^8^, mapped and processed as described in the original studies, was analysed to compare expression levels across gene pair categories defined by interaction status and *P. maximus* chromosome origin. Expression data was log-transformed with a pseudocount of 0.01 to stabilise variance for box plots and statistical testing. To ensure meaningful comparisons, all genes in the ’interacting across coleoids’, ’interacting Decapodiformes only’, ’interacting *O. bimaculoides* only’ were retained, while genes in the ’not in conserved coleoid interactions’ category were included only if they were exclusive to this category and not present in any interacting categories, including in comparisons involving ancestral *P. maximus* chromosomal origin. This filtering approach was applied consistently across all analyses on gene pairs that required single genes as input (e.g. calculation of tissue specificity, GO analyses). Venn diagrams of gene overlap between interaction categories after filtering were produced with the VennDiagram v.1.7.3^77^. Heatmaps were generated with hierarchical clustering of samples and genes using the R package Pheatmap v.1.0.12^78^ to identify patterns among genes in different interaction categories. Rows (tissues) with zero variance in expression were excluded from the heatmaps. Heatmaps were scaled by row (i.e., per gene) using the row-wise scaling function to emphasise relative differences in expression across tissues, except when direct comparisons between multiple heatmaps were required. Tissue specificity of *E. scolopes* gene expression was quantified using the Tau (τ) metric^79^, where values range from 0 (ubiquitous expression) to 1 (highly tissue-specific expression). Tau was calculated for each gene based on its TPM-normalised, log-transformed expression across all tissues, and the mean and median Tau for genes grouped by interaction status and chromosomal organisation was calculated. Differences in tissue specificity between groups were assessed using pairwise Wilcoxon tests with BH correction for multiple comparisons.

### Co-expression analyses

Gene pair co-expression was analysed across different interaction categories using the log-transformed TPM-normalised expression data in *E. scolopes*. Pearson’s correlation coefficients were calculated for each gene pair and loop across tissues, excluding cases where both genes in a pair or all genes within a loop exhibited zero variance. Correlation distributions for gene pair interaction categories were visualised using density plots and statistical differences between interaction categories were assessed using Wilcoxon tests with BH correction.

### Insulation score analyses

Insulation scores were calculated for each interaction matrix using FAN-C v.0.9.28^80^ with a 350 kb window chosen to capture smaller-scale chromatin interactions relevant to loop and domain boundaries. Scores were output in BED format for downstream computational analysis and BigWig format for visualisation in Plotgardener. Insulation scores were averaged between gene pairs for each species using a Python script. NA values were removed before downstream analysis to ensure only gene pairs with valid insulation scores were included. The distributions of these scores were visualised using density plots and statistical differences between interaction categories were assessed using T tests with BH correction.

Next, gene pairs with insulation scores ≥ 0.2 in all species were classified as being within continuous interaction domains, while those with scores ≤ -0.2 in all species were classified as spanning insulating boundaries in all species. Gene pairs with ≥ 0.2 in Decapodiformes but ≤ -0.2 in *O. bimaculoides* were classified as spanning insulating boundaries in *O. bimaculoides* only, whereas pairs with ≤ -0.2 in Decapodiformes but ≥ 0.2 in *O. bimaculoides* were classified as spanning insulating boundaries in Decapodiformes only. The proportion of gene pairs in each insulation score category across different interaction statuses was plotted as a stacked bar plot. Differential insulation scores for an *E. scolopes* loop across developmental stages were plotted using 50 kb iced matrices and abs.bed files from the HiC-Pro output using the R package GENOVA v.1.0.1^81^ at a 750 kb window.

### Quantification of intervening genes

To quantify coding content across gene pairs and chromatin loops, we intersected gene coordinates with gene pairs across the four interaction categories and loop intervals in *E. scolopes* and *O. bimaculoides*, the two species with available gene annotations using a custom Python script. For both datasets, intervening genes exceeding 1.5 Mb in length were excluded as likely misannotations. In R, we filtered for regions containing at least one intervening gene, excluding gene pairs or loop anchors that were immediately adjacent, in order to focus on regions with potential regulatory space. Median values were extracted for all measurements, and differences in gene coverage were assessed using Wilcoxon tests with BH correction, comparing between species, interaction categories and between gene pairs and loops.

### Repeat content analyses

Repeats were identified RepeatModeler v.2.0.6^82^ and RepeatMasker v.4.1.8^83^ using default parameters. For each gene pair, the intergenic region was defined as the genomic interval between the end of the upstream gene and the start of the downstream gene. Repeat features were intersected with intergenic regions using the intersect function in BEDTools v2.30.0, and the number of repeats of each type was counted. To account for differences in intergenic length, the repeat count was divided by the total number of intergenic base pairs to obtain a normalised repeat density for each gene pair. For each interaction category, these values were summed by repeat type and converted to within-category percentages for visualisation as stacked barplots in R. Only gene pairs containing at least one repeat were included in the analysis.

### Chromatin loop calling

Mustache v.1.0.1 ^84^ was used for calling loops and differential loops from KR normalised Juicer .hic files at 50 kb and 100 kb resolution, with default parameters. Loops were considered to be differential if FDR < 0.05. Differential loop analysis was performed on samples with comparable read coverage to minimise potential biases. For example, comparisons across developmental stages were restricted to a single sequencing replicate of stage 29 in *E. scolopes*, to match the data available for the other two stages. Loops detected at 50 kb and 100 kb resolution were merged, with loops in the 100 kb file considered duplicates and removed if they fell within a 50 kb window of those in the 50 kb file.

### Gene association, conservation, and chromosomal context of chromatin loops

Genes were considered associated with a loop if at least one gene overlapped the genomic bins corresponding to both the loop’s start and end positions. Loops sharing the same set of associated genes were merged, and the outermost start and end bins were used to define the final loop coordinates. Loop sizes were calculated as the distance between the start of the start bin and the end of the end bin. Loop size measurements were inspected for outliers potentially resulting from genome misassemblies, and removed if identified as such.

one orthologous gene across both the loop start and end coordinates, regardless of orientation. For each loop conserved in two or more species, we calculated the linear regression slope between genome size and loop size to test whether conserved loop size scales with genome size. A Wilcoxon test was applied to the distribution of per-loop slopes to test whether the median slope differed from zero, indicating a consistent scaling trend across conserved loops.

Loops were also classified as spanning different coleoid or *P. maximus* chromosomes if at least two associated genes had orthologs located on different chromosomes in the comparison species. To test whether loop size is influenced by ancestral gene spacing, we fitted linear models between loop size and the average intergenic distance of orthologous gene pairs located on the same chromosome in another species, using the coefficient of determination (R²) in R to quantify the relationship.

### Regulatory element enrichment analysis of chromatin loops

Because many loops did not have genes at their anchor points, we tested loop anchors across whole-embryo, developmental stage- and tissue-specific samples for significantly (hypergeometric test, BH-adjusted P < 0.5) enriched known transcription factor binding motifs using HOMER v.5.1^85^. Loop anchor regions were defined as the genomic intervals corresponding to the start and end bins of each loop contact point, and converted into BED format using a custom Python script. To enable motif discovery across broad loop anchors, we divided each loop anchor region into 500 bp non-overlapping windows using bedtools makewindows -w 500 -s 500. HOMER was then run with the findMotifsGenome.pl script and the -size given parameter.

Additionally, we tested whether CNEs were enriched in chromatin loop anchors across whole-embryo samples. BED files of CNE, loop anchor, and interloop coordinates were generated using UNIX tools and a custom Python script, then sorted using bedtools sort. Interloop spaces were defined as the genomic span between the end of the first anchor and the start of the second anchor. For each species, genomic background regions not overlapping anchors or interloop segments were defined using bedtools complement. To quantify CNE overlap, we used bedtools intersect -c to count the number of CNEs overlapping each region type (anchor, interloop, and their respective backgrounds). These counts were then normalised by region length (CNEs per kilobase) in R. Comparisons of CNE density between anchors and background, interloop and background, and interloop versus anchor were performed using BH-adjusted Wilcoxon tests.

### ATAC-seq enrichment analysis of chromatin loops

We next tested whether chromatin accessibility is enriched at chromatin loop features using publicly available ATAC-seq data from *E. scolopes* developmental stages 20, 25, and 29^8^, which were mapped and as described in the original publication. ATAC–seq signal tracks were normalised for sequencing depth using bigwigCompare from deepTools v3.5.6^86^, scaling stage 25 and 29 data to the mean signal of stage 20. Normalised bigWig files were converted to BED format with UCSC tools v486^87^, and regions with signal > 1 were retained as open chromatin peaks. Scaffold-level sequences not anchored to chromosomes were removed, and all files were sorted using a common chromosome size file.

BED files of loop, interloop and background regions were generated as in the CNE loop analysis. Overlaps between open chromatin peaks and loop anchors, non-anchor regions, interloop, and non-interloop regions were quantified with bedtools coverage. Significant differences in ATAC enrichment among region types were assessed per developmental stage using Wilcoxon tests with BH correction.

### Comparative analysis of chromatin loop overlap across developmental stages and tissues

Gene-associated loops were considered present in multiple developmental stages or tissues if they had identical gene sets. In analyses involving all loops, regardless of gene association, loops were classified as overlapping across stages or tissues if they shared the same coordinates, defined by the chromosome, the start bin of the loop’s start, and the start bin of the loop’s end. Venn diagrams were generated using VennDiagram in R, and the statistical significance of overlap enrichment and depletion was assessed using the SuperExactTest package^88^ with BH correction.

### Differential compartment analysis

To compare loop conservation with general compartment conservation, TADCompare v.1.12.1^89^ was used to classify and quantify different types of TAD changes across developmental stages and tissues. This analysis was conducted on dumped .hic matrices at 100 kb resolution for the Decapodiform species and 50 kb resolution for *O. bimaculoides*. The matrices were then converted to dense format in R and analysed using a 1 Mb window. Differential compartments shared across comparisons were visualised using UpSet v.1.4.0^90^. If a TAD contained an NA value in the ‘Differential’ classification column of the TADCompare output in at least one comparison file, it was removed from all comparison groups, as its classification as differential could not be reliably determined. Differential compartments were considered overlapping across comparisons if they had identical coordinates and were classified as differential, regardless of the specific change type. Differential compartments were considered unique to a comparison group if they were absent from all other comparisons or classified as ’non-differential’ in all other files.

### Gene ontology analysis across gene pairs and loop-associated genes

Gene Ontology (GO) analyses were performed for genes in gene pairs and tissue-specific chromatin loops to identify functional enrichment in gene pairs and loop-associated genes. InterProScan v.5.62–94.0 ^91^ with the *--goterms* flag was used to annotate protein domains. GO term databases were generated using a custom perl script to produce the necessary annotation files for downstream analyses.

Custom R annotation libraries were built using AnnotationForge v.1.44.0^92^, allowing GO enrichment to be performed in clusterProfiler v.4.10.1^93^. *O. bimaculoides* genes from different interaction, *P. maximus* and loop categories were extracted, and enrichment was tested against background gene sets using the enrichGO function. GO enrichment results were visualised as dot plots and bar plots using dotplot() and barplot() in clusterProfiler.

To further explore gene function, cadherins and zinc finger proteins were specifically examined due to their predicted role in cephalopod neural complexity. The proportion of genes with cadherin or zinc finger annotations was quantified for each gene pair interaction and loop anchor category to assess whether these gene families were enriched in regions of high chromatin organisation and these results were visualised as a barplot. To assess the statistical significance of these results, we performed Fisher’s exact tests using scipy.stats.fisher_exact^94^ in Python. Each gene list was compared against a background gene universe defined as all genes annotated in the respective species’ InterProScan file. Raw P values were BH-adjusted for multiple testing using statsmodels.stats.multitest.multipletests^95^, treating the zinc finger and cadherin test series independently (16 comparisons per group).

### CRISPR-Cas9 knockout of a putative regulatory region in a conserved loop-associated gene Target selection

To identify candidate regulatory elements for functional testing, we analysed gene expression in *E. scolopes* across a panel of tissues for genes located within conserved chromatin loops. One such loop, shared across *E. scolopes*, *E. berryi*, and *O. bimaculoides*, contains Quaking B and several neighboring genes such as SWR1-like, Centrosomal protein of 19 kDa, Futsch-like, and Gastric-lipase-like which are upregulated in neural tissues such as the central brain and optic lobes (Fig. S27A). Many of these genes are involved in neural development or regulatory topological processes, for example, Quaking B in RNA regulation and myelination^47–49^, SWR1-like in chromatin remodeling^96^, Futsch-like in neuronal structure and microtubule stability^97^, and Centrosomal protein of 19 kDa in centrosome function and cell polarity^98^. Notably, a conserved sequence is present in the same intron of Quaking B in all three species. Based on this combination of shared chromatin topology, brain-specific expression, and sequence conservation, we selected this intronic region for CRISPR-Cas9 knockout to test its regulatory function.

Although *E. scolopes* was the primary species of interest, CRISPR experiments were performed in *E. berryi*, a closely related species in which CRISPR-Cas9 protocols have been successfully optimised. Candidate guide RNAs were selected using the CRISPRscan tool^99^. Guides were prioritised based on high CRISPRscan scores, proximity to the target *E. berry*i intron sequence, and the presence of a canonical NGG PAM sequence. Potential off-target effects were minimised by selecting guides with minimal predicted off-target binding sites in the *E. berryi* reference genome^63^, using TBLASTN with default parameters (E-value threshold = 10).

To assess potential regulatory element activity within the candidate Quaking B knockout region (*E. berryi* chromosome 19: 27,035,572–27,036,708 bp), the sequence was divided into overlapping 200 bp windows with a 100 bp step size. Motif enrichment was then performed using HOMER as in the loop analyses.

### Embryo collection and culture

Adult *E. berryi* were kept in a breeding system at the MBL and embryos were collected as described in Ahuja et al. (2023). Briefly, mated females deposited egg clutches on provided structures (rocks or artificial substrates) during the night and freshly laid embryos were harvested in the morning. Embryos were kept at 24°C in 60 mm petri dishes (Fisher Scientific, FB0875713A) in filtered (0.22 µm Vacuum Filter System, Corning, 431205) natural sea water (FNSW) supplemented with Pen-Strep (1:100, Gibco, 10000 U/mL, 15140-122) until collection, with no more than 25 embryos per dish. Sea water was changed daily and dead embryos removed immediately.

### Injection

To prepare embryos for injections, the outer and inner jelly layers were removed under a dissection scope using fine forceps (Number 3 and Number 5 Dumoxel, Fine Science Tools) until chorion is exposed and accessible for injections. Embryos were placed cell side up in the trough of an injection plate. Injection plates were prepared by filling a 10cm petri dish (Fisher Scientific, FB0875712) holding a 20-gauge insulated wire with molten 1% Type II-A agarose (Sigma-Aldrich, A9918, medium EEO) in FNSW. Once the agarose solidified, the wire was removed, creating an injection trough approximately the size of an embryo. Injection needles were pulled on a P-2000 Laser-based Micropipette puller (settings: heat 700, filament 4, delay 140, pull 175; Sutter Instruments) using quartz capillaries (OD=1.0mm, ID=0.7mm, OL=10cm; Sutter Instruments, QF100-70-10) and then beveled at a 25° angle for 30 seconds on a BV-10 Microelectrode Beveler (Sutter instruments) to generate a 3-4 µm wide tip opening. Injection needles were then back filled with 1.5 µL injection mix and at the 2-cell stage each embryo was injected with 0.22 pL. For injection mixes lyophilised guide RNAs (1.5 nmol, Synthego Corporation) were resuspended in an injection buffer (10 mM Tris-HCl pH 8, 0.1 mM EDTA in nuclease-free water) to a concentration of 100 µM. The final injection mix was prepared to consist of 11µM CRISPR gRNA (each), 7 µM Cas9 protein (SpCas9 2NLS Nuclease, Synthego Corperation) and 0.9µM fluorescent tetramethylrhodamine-dextran (ThermoFisher, D18180 isothiocyanate, used as a tracer to identify successful injections. Injected embryos were removed from the trough and cultured as described above.

### Genotyping

Injected and uninjected control embryos were collected at late-organogenic stages and anesthetised for 20 min in 3.75% Magnesium chloride (Fisher Scientific, M33-500) in FNSW + Pen-Strep. Embryos were then individually flash-frozen in liquid nitrogen and stored at -80°C. For isolation of genomic DNA embryos were incubated in Digestion buffer (0.01M Tris pH 7.5, 0.2M EDTA, 0.4M NaCL and 1% SDS) and 1.2 mg/mL Proteinase K (Sigma-Aldrich, 3115887001) at 55°C until tissue was fully digested (∼ 3h). Natrium Chloride was added to a concentration of 1.25M and tissue debris pelleted through centrifugation. Supernatant was transferred to a new tube and DNA was ethanol-precipitated, air-dried and resuspended in nuclease-free water.

The genomic region spanning the gRNA target sites was PCR amplified using a forward primer with the sequence 5’-TTACATGCTTCGAGGAAAGACC-3’ and a reverse primer with the sequence 5’-AAGACAAGCTACAAACATGCCA-3’, with an expected band size of 1390 bp for wild type and 255 bp for a complete knockout. PCR products were run on a 2% agarose (Sigma-Aldrich, A9793) gel and resulting bands were gel purified using the Monarch DNA Gel Extraction kit (NEB, T1020) following Manufacturer’s instructions. Purified PCR products were analyzed using Sanger Sequencing services provided by Genewiz (South Plainfield, NJ).

### RNA sequencing

For RNA sequencing analysis, injected and uninjected control embryos were grown to stage 14 (epiboly) and stage 28 (late organogenesis) and anesthetised in 3.75% Magnesium chloride for 20 min. Embryos were individually flash frozen in liquid nitrogen. An initial crude homogenization was performed on dry, frozen tissue using a pellet pestle (DWK Life Sciences Kimble™ Kontes™ Pellet Pestle™, Fisher Scientific, K749521-1590). Tissue debris stuck to the pestle was transferred into digestion buffer and processed for genotyping as described above. TRIzol Reagent (Invitrogen, 15596026) was added to the remaining tissue, followed by full homogenization using a pellet pestle while keeping the tissue frozen on dry ice. RNA was extracted using standard TRIzol Reagent RNA isolation protocols. Briefly, chloroform was added to each sample, mixed by inverting and phases were separated through centrifugation. The aqueous phase was transferred to a new tube, RNA precipitated using ice-cold isopropanol and RNA pellet was resuspended in nuclease-free water. RNA samples were shipped to the University of Chicago Genomics Facility for library preparation and sequencing on the Illumina NovaSeqX platform.

Raw RNA-seq reads were trimmed with Trim Galore v0.6.10 (default settings and --clip_R1 5 --clip_R2 5). Mapping to the reference genome was performed using STAR v2.7.11b (--runMode alignReads)^100^. Read counts were obtained with featureCounts v2.1.1 against the available genome annotation of *E. berryi* (-g transcript_id -t exon -M -a Eberryi_GmvKQ_ah2p_ext3_000.wnm.cleanv2.gtf). The read count matrix was processed in R v4.5.0 using DESeq2 v1.48.1. Differential expression was tested using the Wald test, with Benjamini-Hochberg adjusted P values and log2 fold changes extracted from the DESeq2 results object. Variance stabilising transformation (VST) was applied to normalise counts prior to visualizations. PCA was performed on VST-transformed data separately for stage 14 and 28 embryos to assess expression separation between injected and control conditions, and to evaluate phenotypic clustering at stage 28. Heatmaps were generated from VST-normalised data for selected gene sets using the pheatmap package, and group-averaged expression values were computed across conditions and developmental stages. Genes with zero variance in expression across samples were removed from heatmaps.

## Acknowledgements

The computational results of this work were achieved using the Life Science Compute Cluster (LiSC) of the University of Vienna. We also acknowledge the Vienna BioCenter Core Facilities (VBCF) for performing the Micro-C sequencing. We thank the head aquarist Roland Halbauer and the team at the aqua facility of Vienna Zoo. We thank Fatih Sarigoel for advice on the computational analyses of this study. T.F.R., G.Y., D.T.S. and O.S. were supported by the ERC Horizon 2020: European Union Research and Innovation Programme (grant no. 945026). S.R. was supported by ERC grant CAMOUFLAGE, 101141501 to Gilles Laurent. C.W.R. and N.S. were supported by the NSF grant IOS-1354898. C.B.A. and J.S. acknowledge generous support through the MBL Early Career Fellows gift from Susan and David Hibbitt, NSF EDGE 2220587 and NIH R35GM147273. O.S., C.W.R., C.A. were supported by the Chicago-Vienna Strategic Partnership grant.

## Author contributions

T.F.R. led the research. T.F.R. and O.S. conceived and designed the research. J.S. and C.A. carried out and advised on the wet lab components of the CRISPR knockout experiment. G.Y. contributed to figure design and performed initial data analyses. N.S. provided and dissected *O. bimaculoides* samples, while S.R. provided and dissected *S. officinalis* samples. A.W. provided *E. scolopes* samples. D.T.S. conducted the multi-locus topology analysis. C.R. and T.C. provided expertise in the analysis of cephalopod and genome topology data. T.F.R. and O.S. carried out the remaining analyses. T.F.R. and O.S. wrote the manuscript with input from all authors.

## References

1. Simakov, O. et al. Deeply conserved synteny resolves early events in vertebrate evolution. Nat Ecol Evol 4, 820–830 (2020).

2. Zimmermann, B., Robert, N. S. M., Technau, U. & Simakov, O. Ancient animal genome architecture reflects cell type identities. *Nat*. Ecol. Evol. 3, 1289–1293 (2019).

3. Irimia, M. et al. Extensive conservation of ancient microsynteny across metazoans due to cis-regulatory constraints. Genome Res. 22, 2356–2367 (2012).

4. Simakov, O. et al. Insights into bilaterian evolution from three spiralian genomes. Nature 493, 526–531 (2013).

5. Kikuta, H. et al. Genomic regulatory blocks encompass multiple neighboring genes and maintain conserved synteny in vertebrates. Genome Res. 17, 545–555 (2007).

6. Engström, P. G., Ho Sui, S. J., Drivenes, O., Becker, T. S. & Lenhard, B. Genomic regulatory blocks underlie extensive microsynteny conservation in insects. Genome Res. 17, 1898–1908 (2007).

7. Robert, N. S. M. et al. Emergence of distinct syntenic density regimes is associated with early metazoan genomic transitions. BMC Genomics 23, 143 (2022).

8. Schmidbaur, H. et al. Emergence of novel cephalopod gene regulation and expression through large-scale genome reorganization. Nat. Commun. 13, 2172 (2022).

9. Acemel, R. D. & Lupiáñez, D. G. Evolution of 3D chromatin organization at different scales. Curr. Opin. Genet. Dev. 78, 102019 (2023).

10. Harmston, N. et al. Topologically associating domains are ancient features that coincide with Metazoan clusters of extreme noncoding conservation. Nat. Commun. 8, 441 (2017).

11. Rogers, T. F. & Simakov, O. Emerging questions on the mechanisms and dynamics of 3D genome evolution in spiralians. Brief. Funct. Genomics 22, 533–542 (2023).

12. Szalay, M.-F., Majchrzycka, B., Jerković, I., Cavalli, G. & Ibrahim, D. M. Evolution and function of chromatin domains across the tree of life. Nat. Struct. Mol. Biol. 1–14 (2024).

13. Smith, E. M., Lajoie, B. R., Jain, G. & Dekker, J. Invariant TAD boundaries constrain cell-type-specific looping interactions between promoters and distal elements around the CFTR locus. Am. J. Hum. Genet. 98, 185–201 (2016).

14. Álvarez-González, L. et al. 3D chromatin remodelling in the germ line modulates genome evolutionary plasticity. Nat. Commun. 13, 2608 (2022).

15. Torosin, N. S., Anand, A., Golla, T. R., Cao, W. & Ellison, C. E. 3D genome evolution and reorganization in the Drosophila melanogaster species group. PLoS Genet. 16, e1009229 (2020).

16. Farré, M., Robinson, T. J. & Ruiz-Herrera, A. An Integrative Breakage Model of genome architecture, reshuffling and evolution: The Integrative Breakage Model of genome evolution, a novel multidisciplinary hypothesis for the study of genome plasticity. Bioessays 37, 479–488 (2015).

17. Krefting, J., Andrade-Navarro, M. A. & Ibn-Salem, J. Evolutionary stability of topologically associating domains is associated with conserved gene regulation. BMC Biol. 16, 87 (2018).

18. Eres, I. E. & Gilad, Y. A TAD skeptic: Is 3D genome topology conserved? Trends Genet. 37, 216–223 (2021).

19. Kim, I. V. et al. Chromatin loops are an ancestral hallmark of the animal regulatory genome. Nature (2025) doi:10.1038/s41586-025-08960-w.

20. Vara, C. et al. The impact of chromosomal fusions on 3D genome folding and recombination in the germ line. Nat. Commun. 12, 2981 (2021).

21. Bird, A. Cohesin as an essential disruptor of chromosome organization. Mol. Cell (2025) doi:10.1016/j.molcel.2025.01.010.

22. Simakov, O. & Wagner, G. P. The application of irreversible genomic states to define and trace ancient cell type homologies. Evodevo 16, 5 (2025).

23. Tanner, A. R. et al. Molecular clocks indicate turnover and diversification of modern coleoid cephalopods during the Mesozoic Marine Revolution. Proc. Biol. Sci. 284, (2017).

24. Albertin, C. B. & Katz, P. S. Evolution of cephalopod nervous systems. Curr. Biol. 33, R1087–R1091 (2023).

25. Hanlon, R. T. & Messenger, J. B. Cephalopod Behaviour. (Cambridge University Press, 2018).

26. Hanlon, R. Cephalopod dynamic camouflage. Curr. Biol. 17, R400–4 (2007).

27. Kerwin, A. H. & Nyholm, S. V. Symbiotic bacteria associated with a bobtail squid reproductive system are detectable in the environment, and stable in the host and developing eggs. Environ. Microbiol. 19, 1463–1475 (2017).

28. Nyholm, S. V., Stewart, J. J., Ruby, E. G. & McFall-Ngai, M. J. Recognition between symbiotic Vibrio fischeri and the haemocytes of Euprymna scolopes. Environ. Microbiol. 11, 483–493 (2009).

29. Belcaid, M. et al. Symbiotic organs shaped by distinct modes of genome evolution in cephalopods. Proc. Natl. Acad. Sci. U. S. A. 116, 3030–3035 (2019).

30. Albertin, C. B. et al. Genome and transcriptome mechanisms driving cephalopod evolution. Nat. Commun. 13, 2427 (2022).

31. Albertin, C. B. et al. The octopus genome and the evolution of cephalopod neural and morphological novelties. Nature 524, 220–224 (2015).

32. Marino, A., Kizenko, A., Wong, W. Y., Ghiselli, F. & Simakov, O. Repeat age decomposition informs an ancient set of repeats associated with coleoid cephalopod divergence. Front. Genet. 13, 793734 (2022).

33. Yoshida, M.-A. et al. Chromosomal fusions and subsequent rearrangements shaped octopus genomes. bioRxiv 2025.05.16.652989 (2025) doi:10.1101/2025.05.16.652989.

34. Schultz, D. T., Blümel, A., Destanović, D., Sarigol, F. & Simakov, O. Topological mixing and irreversibility in animal chromosome evolution. bioRxiv 2024.07.29.605683 (2024) doi:10.1101/2024.07.29.605683.

35. Simakov, O. et al. Deeply conserved synteny and the evolution of metazoan chromosomes. Sci. Adv. 8, eabi5884 (2022).

36. Swygert, S. G. et al. Local chromatin fiber folding represses transcription and loop extrusion in quiescent cells. Elife 10, (2021).

37. Doni Jayavelu, N., Jajodia, A., Mishra, A. & Hawkins, R. D. Candidate silencer elements for the human and mouse genomes. Nat. Commun. 11, 1061 (2020).

38. Furlong, E. E. M. & Levine, M. Developmental enhancers and chromosome topology. Science 361, 1341–1345 (2018).

39. Schoenfelder, S. & Fraser, P. Long-range enhancer-promoter contacts in gene expression control. Nat. Rev. Genet. 20, 437–455 (2019).

40. Rao, S. S. P. et al. A 3D map of the human genome at kilobase resolution reveals principles of chromatin looping. Cell 159, 1665–1680 (2014).

41. Bing, X. et al. Chromosome structure in Drosophila is determined by boundary pairing not loop extrusion. Elife 13, (2024).

42. Castellanos-Martínez, S. & Gestal, C. Pathogens and immune response of cephalopods. J. Exp. Mar. Bio. Ecol. 447, 14–22 (2013).

43. Bond, M. L. et al. Chromatin loop dynamics during cellular differentiation are associated with changes to both anchor and internal regulatory features. Genome Res. 33, 1258–1268 (2023).

44. Lee, P. N., Callaerts, P. & de Couet, H. G. The embryonic development of the Hawaiian bobtail squid (Euprymna scolopes). Cold Spring Harb. Protoc. 2009, db.ip77 (2009).

45. Imarazene, B., Andouche, A., Bassaglia, Y., Lopez, P.-J. & Bonnaud-Ponticelli, L. Eye development in Sepia officinalis embryo: What the uncommon gene expression profiles tell us about eye evolution. Front. Physiol. 8, 613 (2017).

46. Liu, Y.-C., Liu, T.-H., Su, C.-H. & Chiao, C.-C. Neural organization of the optic lobe changes steadily from late embryonic stage to adulthood in cuttlefish sepia pharaonis. Front. Physiol. 8, 538 (2017).

47. Wu, J. I., Reed, R. B., Grabowski, P. J. & Artzt, K. Function of quaking in myelination: regulation of alternative splicing. Proc. Natl. Acad. Sci. U. S. A. 99, 4233–4238 (2002).

48. Sakers, K. et al. Loss of Quaking RNA binding protein disrupts the expression of genes associated with astrocyte maturation in mouse brain. Nat. Commun. 12, 1537 (2021).

49. Zhou, X. et al. Qki regulates myelinogenesis through Srebp2-dependent cholesterol biosynthesis. Elife 10, e60467 (2021).

50. Mohana, G. et al. Chromosome-level organization of the regulatory genome in the Drosophila nervous system. Cell 186, 3826–3844.e26 (2023).

51. Li, X. & Levine, M. What are tethering elements? Curr. Opin. Genet. Dev. 84, 102151 (2024).

52. Altschul, S. F., Gish, W., Miller, W., Myers, E. W. & Lipman, D. J. Basic local alignment search tool. J. Mol. Biol. 215, 403–410 (1990).

53. Simakov, O. et al. Hemichordate genomes and deuterostome origins. Nature 527, 459– 465 (2015).

54. Li, F. et al. Chromosome-level genome assembly of the East Asian common octopus (*Octopus sinensis*) using PacBio sequencing and Hi-C technology. Mol. Ecol. Resour. 20, 1572–1582 (2020).

55. Destanoviæ, D. et al. A chromosome-level reference genome for the common octopus, Octopus vulgaris (Cuvier, 1797). G3 (Bethesda) 13, (2023).

56. Gavriouchkina, D., et al. A single-cell atlas of bobtail squid visual and nervous system highlights molecular principles of convergent evolution. *bioRxiv* (2022) doi:10.1101/2022.05.26.490366.

57. Li, W. & Godzik, A. Cd-hit: a fast program for clustering and comparing large sets of protein or nucleotide sequences. Bioinformatics 22, 1658–1659 (2006).

58. Schultz, D. T. et al. Ancient gene linkages support ctenophores as sister to other animals. Nature 618, 110–117 (2023).

59. Shigeno, S., Parnaik, R., Albertin, C. B. & Ragsdale, C. W. Evidence for a cordal, not ganglionic, pattern of cephalopod brain neurogenesis. Zoological Lett. 1, 26 (2015).

60. Bolger, A. M., Lohse, M. & Usadel, B. Trimmomatic: a flexible trimmer for Illumina sequence data. Bioinformatics 30, 2114–2120 (2014).

61. Li, H. Seqtk. (Github, 2008).

62. National Center for Biotechnology Information (NCBI). Sepia officinalis genome assembly xcSepOffi3.1. NCBI Assembly Database. Accession GCA_964300435.1.

63. Gavriouchkina, D. et al. A single-cell atlas of the bobtail squid visual and nervous system highlights molecular principles of convergent evolution. *Nat*. Ecol. Evol. 1–18 (2025).

64. Servant, N. et al. HiC-Pro: an optimized and flexible pipeline for Hi-C data processing. Genome Biol. 16, 259 (2015).

65. Durand, N. C. et al. Juicer provides a one-click system for analyzing loop-resolution Hi-C experiments. Cell Syst. 3, 95–98 (2016).

66. Robinson, J. T. et al. Juicebox.Js provides a cloud-based visualization system for Hi-C data. Cell Syst. 6, 256–258.e1 (2018).

67. Kramer, N. E. et al. Plotgardener: cultivating precise multi-panel figures in R. Bioinformatics 38, 2042–2045 (2022).

68. National Center for Biotechnology Information (NCBI). Pecten maximus genome assembly xPecMax1.1. NCBI RefSeq Database. Accession GCF_902652985.1.

69. Huang, Z. et al. Genomic insights into the adaptation and evolution of the nautilus, an ancient but evolving ‘living fossil’. Mol. Ecol. Resour. 22, 15–27 (2022).

70. Li, H. Protein-to-genome alignment with miniprot. Bioinformatics 39, btad014 (2023).

71. R Core Team. R: A Language and Environment for Statistical Computing. Preprint at https://www.R-project.org/ (2023).

72. Wickham, H., François, R., Henry, L., Müller, K. & D, Vaughan. dplyr: A Grammar of Data Manipulation. R Package. Https://github.com/tidyverse/dplyr, Https://dplyr.tidyverse.org. (2023).

73. Wickham, H., Vaughan, D. & Girlich, M. tidyr: Tidy Messy Data. R Package. Https://github.com/tidyverse/tidyr, Https://tidyr.tidyverse.org. (2024).

74. *Wickham, H.* stringr: Simple, Consistent Wrappers for Common String Operations. R Package. Https://github.com/tidyverse/stringr, Https://stringr.tidyverse.org. (2023).

75. Wickham, H. ggplot2: Elegant Graphics for Data Analysis. (2016).

76. Rouressol, L. et al. Emergence of novel genomic regulatory regions associated with light-organ development in the bobtail squid. iScience 26, 107091 (2023).

77. Chen, H. & Boutros, P. C. VennDiagram: a package for the generation of highly-customizable Venn and Euler diagrams in R. BMC Bioinformatics 12, 35 (2011).

78. Kolde, R. pheatmap: Pretty Heatmaps. R Package Version 1.0.12, https://CRAN.R-Project.org/package=pheatmap. (2019).

79. Yanai, I. et al. Genome-wide midrange transcription profiles reveal expression level relationships in human tissue specification. Bioinformatics 21, 650–659 (2005).

80. Kruse, K., Hug, C. B. & Vaquerizas, J. M. FAN-C: a feature-rich framework for the analysis and visualisation of chromosome conformation capture data. Genome Biol. 21, 303 (2020).

81. van der Weide, R. H., et al. Hi-C analyses with GENOVA: a case study with cohesin variants. NAR Genom. Bioinform. 3, lqab040 (2021).

82. Flynn, J. M. et al. RepeatModeler2 for automated genomic discovery of transposable element families. Proc. Natl. Acad. Sci. U. S. A. 117, 9451–9457 (2020).

83. Smit, A. F. A., Hubley, R. & Green, P. RepeatMasker Open-4.0. Institute for Systems Biology, Seattle, WA. Available At: Http://www.repeatmasker.org. (2015).

84. Roayaei Ardakany, A., Gezer, H. T., Lonardi, S. & Ay, F. Mustache: multi-scale detection of chromatin loops from Hi-C and Micro-C maps using scale-space representation. Genome Biol. 21, 256 (2020).

85. Heinz, S. et al. Simple combinations of lineage-determining transcription factors prime cis-regulatory elements required for macrophage and B cell identities. Mol. Cell 38, 576–589 (2010).

86. Ramírez, F. et al. deepTools2: a next generation web server for deep-sequencing data analysis. Nucleic Acids Res. 44, W160–5 (2016).

87. Perez, G. et al. The UCSC Genome Browser database: 2025 update. Nucleic Acids Res. 53, D1243–D1249 (2025).

88. Wang, M., Zhao, Y. & Zhang, B. Efficient test and visualization of multi-set intersections. Sci. Rep. 5, 16923 (2015).

89. Cresswell, K. G. & Dozmorov, M. G. TADCompare: An R package for differential and temporal analysis of topologically associated domains. Front. Genet. 11, 158 (2020).

90. Conway, J. R., Lex, A. & Gehlenborg, N. UpSetR: an R package for the visualization of intersecting sets and their properties. Bioinformatics 33, 2938–2940 (2017).

91. Jones, P. et al. InterProScan 5: genome-scale protein function classification. Bioinformatics 30, 1236–1240 (2014).

92. Carlson, M. & Pagès, H. AnnotationForge: Tools for Building SQLite-Based Annotation Data Packages. https://bioconductor.org/packages/AnnotationForge. (2024).

93. Yu, G. Thirteen years of clusterProfiler. Innovation (Camb.) 5, 100722 (2024).

94. Virtanen, P. et al. SciPy 1.0: fundamental algorithms for scientific computing in Python. Nat. Methods 17, 261–272 (2020).

95. Seabold, S. & Perktold, J. Statsmodels: Econometric and statistical modeling with Python. Proceedings of the 9th Python in Science Conference (SciPy2010) 92– (2010).

96. Mizuguchi, G. et al. ATP-driven exchange of histone H2AZ variant catalyzed by SWR1 chromatin remodeling complex. Science 303, 343–348 (2004).

97. Roos, J., Hummel, T., Ng, N., Klämbt, C. & Davis, G. W. Drosophila Futsch regulates synaptic microtubule organization and is necessary for synaptic growth. Neuron 26, 371–382 (2000).

98. UniProt Consortium. UniProt: The universal protein knowledgebase in 2025. Nucleic Acids Res. 53, D609–D617 (2025).

99. Moreno-Mateos, M. A. et al. CRISPRscan: designing highly efficient sgRNAs for CRISPR-Cas9 targeting in vivo. Nat. Methods 12, 982–988 (2015).

100. Dobin, A. et al. STAR: ultrafast universal RNA-seq aligner. Bioinformatics 29, 15–21 (2013).

101. Sanchez, G. et al. Phylogenomics illuminates the evolution of bobtail and bottletail squid (order Sepiolida). Commun Biol 4, 819 (2021).

