## Supplementary figures for "Genome reorganisation and expansion shape 3D genome architecture and define a distinct regulatory landscape in coleoid cephalopods"

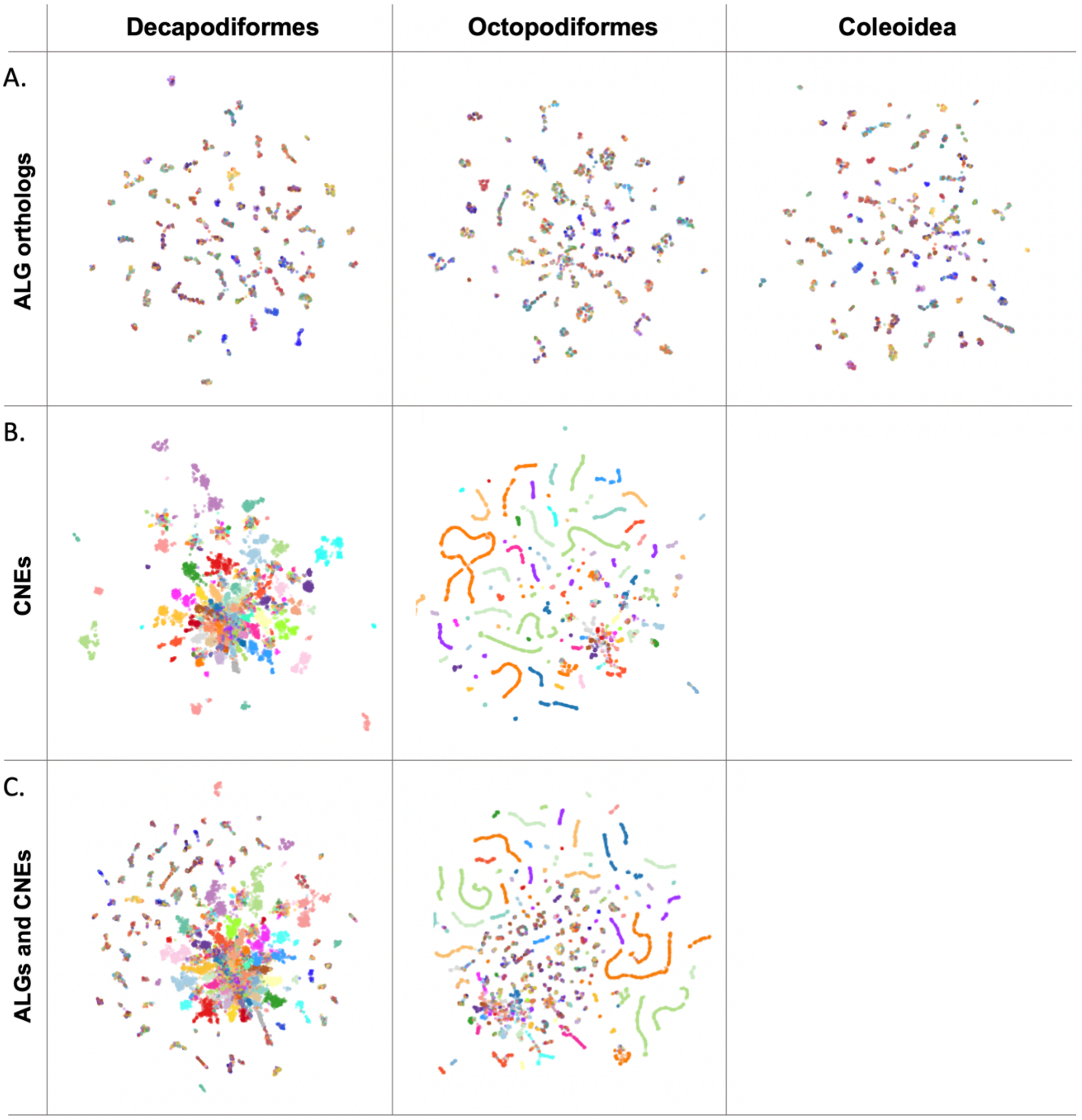


**Figure S1. Multi-locus topology of ALG orthologs and conserved non-coding elements (CNEs) across coleoid cephalopods**.

Each point represents a genomic locus, coloured by either: (A) ALG orthologs identified in BCnS (Simakov et al. 2022), or (B, C) by chromosome of origin: *E. scolopes* chromosomes for Decapodiform CNEs and *O. bimaculoides* chromosomes for Octopodiform CNEs. UMAP was performed using 15 nearest neighbours and a minimum distance of 0.75.

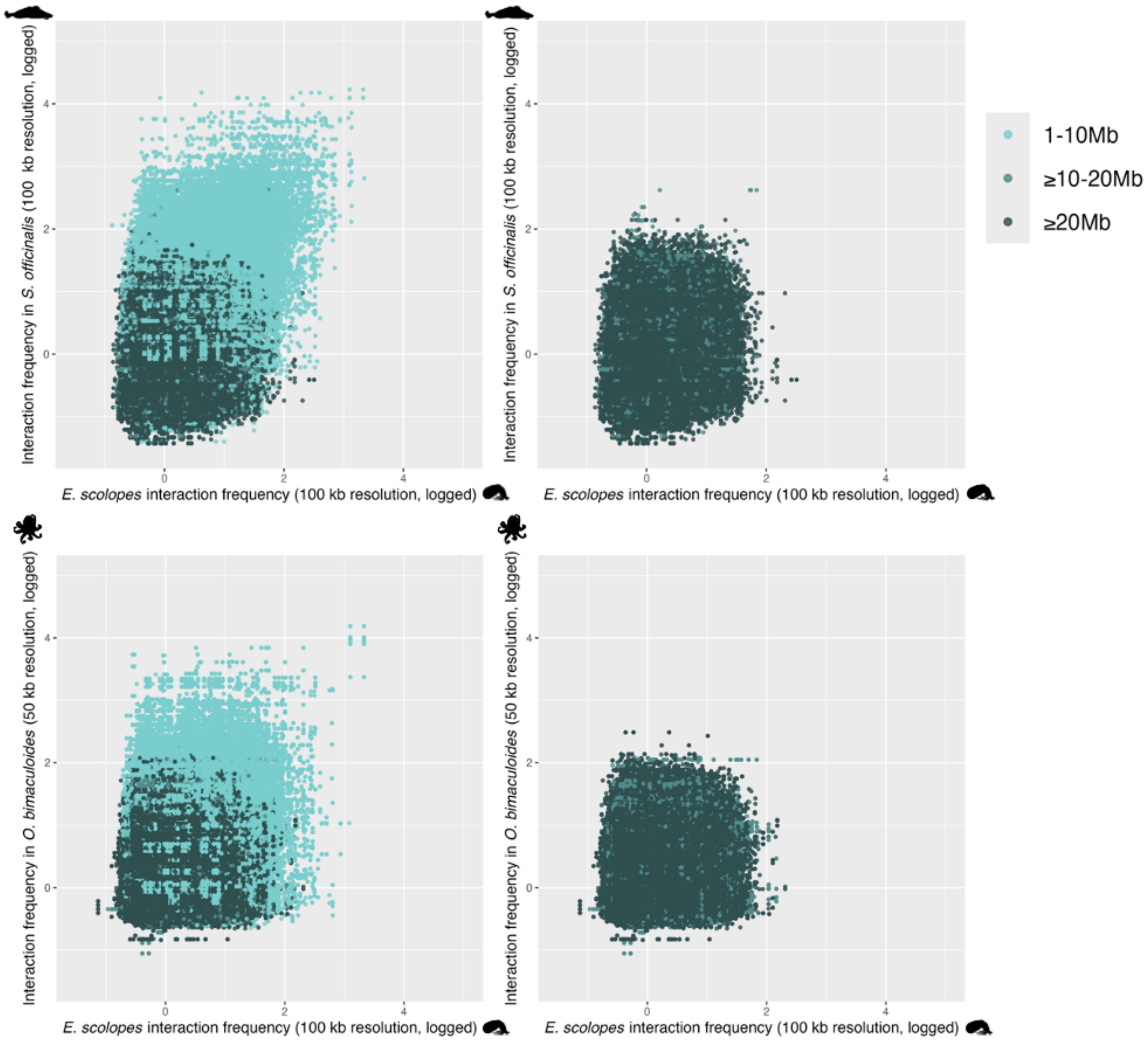


**Figure S2. Justification of interaction frequency threshold based on distance-dependent signal decay across species.**

Pairwise interaction frequencies (log-transformed) are plotted between *E. scolopes* (x-axis) and either *S. officinalis* or *O. bimaculoides* (y-axis) at 100 kb or 50 kb resolution, respectively, for orthologous gene pairs Points are coloured by genomic distance: 1–10 Mb (light blue), 10–20 Mb (medium blue), and ≥20 Mb (dark blue). Interactions involving regions ≥20 Mb apart rarely exceed 9 KR-normalised reads, providing a basis for setting the threshold of ≥10 reads to define interacting gene pairs. This threshold captures topological interactions primarily within 20 Mb and minimises inclusion of artefacts due to long-range noise or misassemblies.

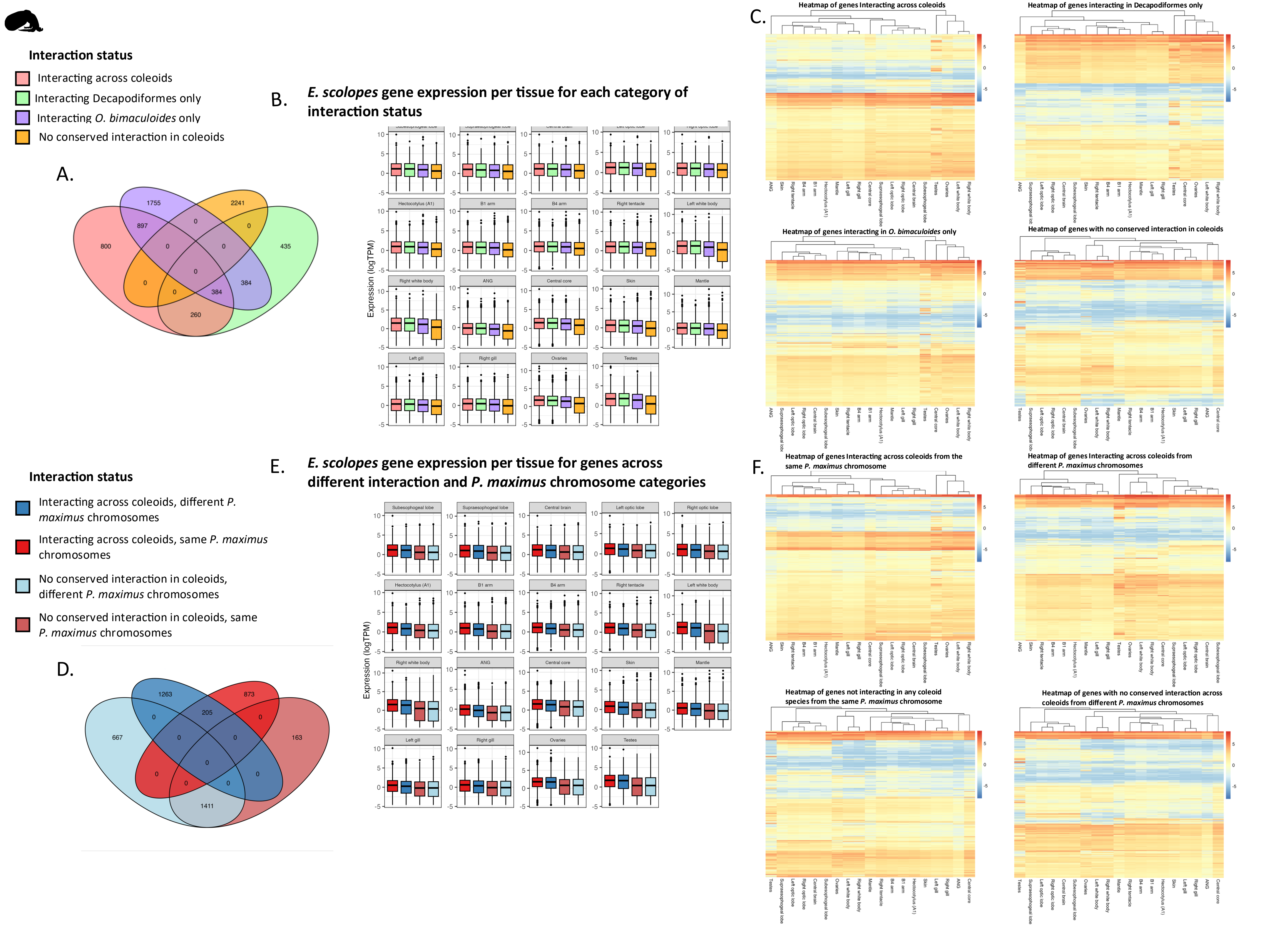
Interaction status
Interacting across coleoids
Interacting Decapodiformes only
Interacting O. bimaculoides only
No conserved interaction in coleoids
A.
B.
E. scolopes gene expression per tissue for each category of interaction status
Expression (logTPM)
C.
Heatmap of genes Interacting across coleoids
Heatmap of genes interacting in Decapodiformes only
Heatmap of genes interacting in O. bimaculoides only
Heatmap of genes with no conserved interaction in coleoids
Interaction status
Interacting across coleoids, different P. maximus chromosomes
Interacting across coleoids, same P. maximus chromosomes
No conserved interaction in coleoids, different P. maximus chromosomes
No conserved interaction in coleoids, same P. maximus chromosomes
D.
E.
E. scolopes gene expression per tissue for genes across different interaction and P. maximus chromosome categories
F.
Heatmap of genes Interacting across coleoids from the same P. maximus chromosome
Heatmap of genes Interacting across coleoids from different P. maximus chromosomes
Heatmap of genes not interacting in any coleoid species from the same P. maximus chromosome
Heatmap of genes with no conserved interaction across coleoids from different P. maximus chromosomes

**Figure S3. Expression patterns of *E. scolopes* genes across tissues in relation to interaction conservation and ancestral chromosomal organisation.**

(A) Venn diagram showing the overlap of genes across interaction categories included in gene expression analyses. Only genes unique to the 'no conserved interaction in coleoids' category were retained for comparisons in analyses of single genes (i.e. not paired).

(B) Boxplots showing log-transformed TPM expression of *E. scolopes* genes across tissues, grouped by interaction status: interacting in across coleoids (pink), Decapodiformes only (green), *O. bimaculoides* only (purple), or no conserved interaction in coleoids (orange). Wilcoxon test significance values comparing interaction categories within each tissue are provided in Table S3.

(C) Heatmaps of log-transformed TPM expression across *E. scolopes* tissues for genes in each interaction category.

(D) Venn diagram showing the overlap of genes across combined interaction and *P. maximus* chromosome status categories included in the expression analyses. Again, only genes unique to the 'no conserved interaction in coleoids' category were retained for comparisons, as in all analyses of single genes (i.e. not paired).

(E) Boxplots showing log-transformed TPM expression across *E. scolopes* tissues for genes interacting across coleoids or with no conserved interaction in coleoids, grouped by *P. maximus* chromosome status (same chromosome, interacting: dark red; same chromosome, non-interacting: light red; different chromosome, interacting: dark blue; different chromosome, non-interacting: light blue). Wilcoxon test significance values comparing interaction categories within each tissue are provided in Table S6.

(F) Heatmaps showing log-transformed TPM expression across *E. scolopes* tissues for genes interacting in all species or not interacting in any species, grouped by *P. maximus* chromosome status.

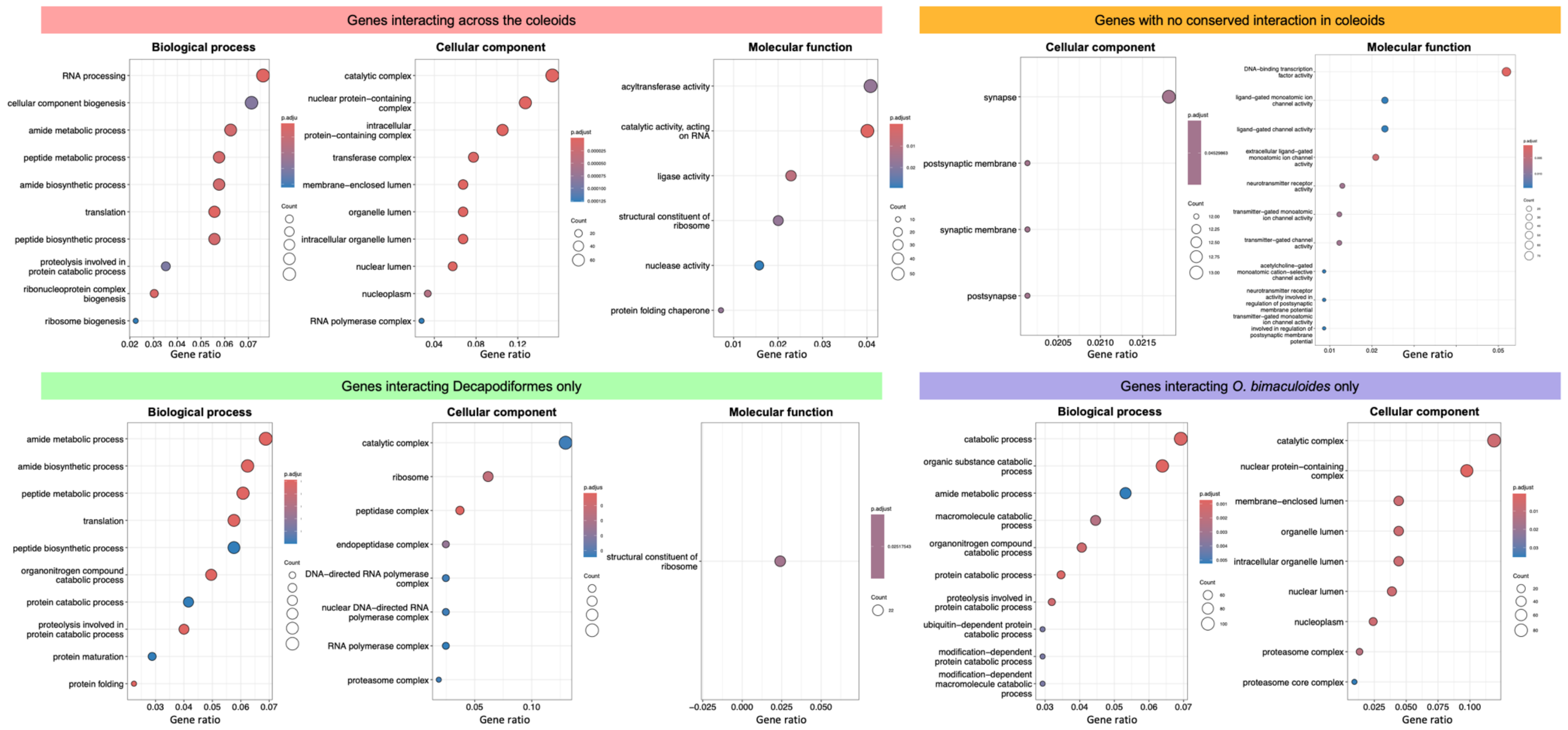


**Figure S4. Gene Ontology (GO) enrichment analysis for genes in each interaction category.**
Dot plots showing enriched GO terms (biological process, cellular component, molecular function) for gene pairs grouped by interaction status: Genes interacting in across the coleoids (pink), genes with no conserved interaction in coleoids (orange), genes interacting in Decapodiformes only (green), genes interacting only in *O. bimaculoides* (purple). For each GO category, enriched GO terms are shown along the y-axis. Dot size indicates the number of genes associated with each term, and colour reflects statistical significance (adjusted p-value). The gene ratio, shown on the x-axis, represents the proportion of input genes annotated with each GO term (number of genes in the term divided by total number of genes in the input list).

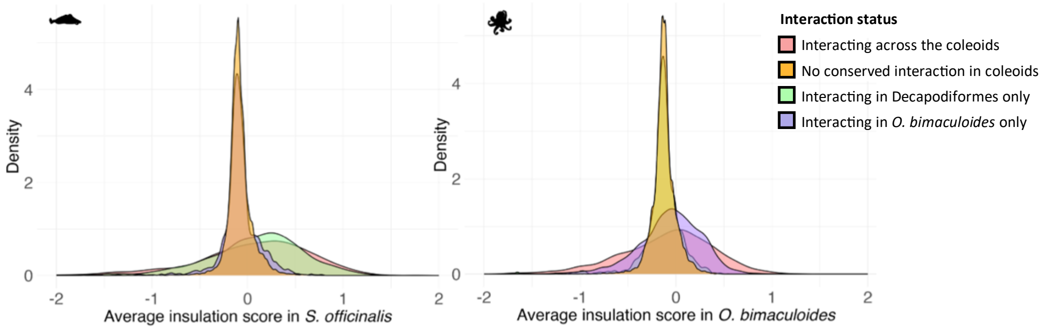


**Figure S5**. **Insulation score distributions for gene pairs grouped by interaction conservation.** Density plots of average insulation scores between gene pairs in *S. officinalis* (left) and *O. bimaculoides* (right), grouped by interaction status across species. Insulation scores were computed from Micro-C data at 100 kb resolution using a 350 kb window in *S. officinalis* and at 50 kb resolution with a 350 kb window in *O. bimaculoides*. All within-species pairwise comparisons were significant (BH-corrected T test, $P < 0.05$), except comparisons between the categories 'no conserved interaction in coleoids' vs. 'interacting in *O. bimaculoides* only' and 'interacting across the coleoids' vs. 'interacting in Decapodiformes only' in *S. officinalis* and 'interacting across the coleoids' vs. 'interacting in *O. bimaculoides* only', in *O. bimaculoides*.

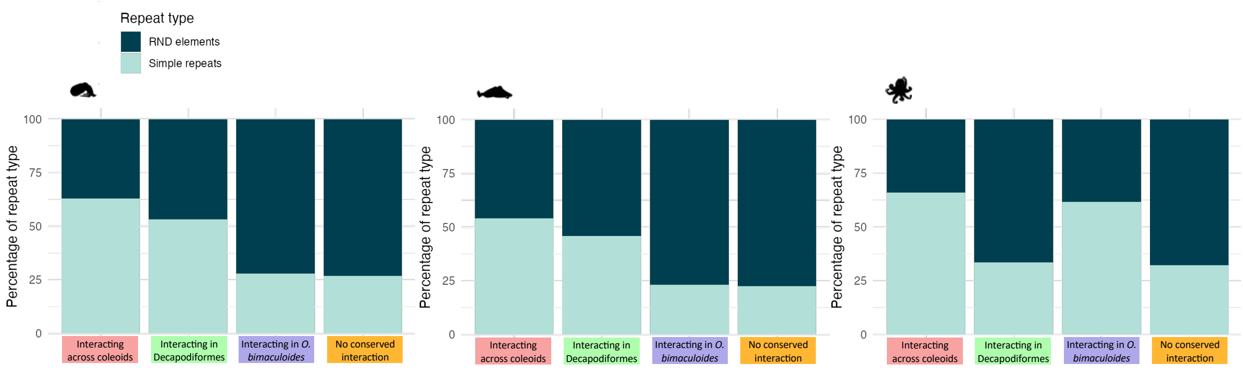


**Figure S6**. **Repeat content for gene pairs grouped by interaction conservation.**
Stacked bar plots showing the percentage of RND elements (complex repeats identified by RepeatModeler) and simple repeats found in the genomic intervals between gene pairs across three coleoid species (*E. scolopes*, *S. officinalis*, and *O. bimaculoides*). Gene pairs are grouped by interaction status. Across all species and categories, RND elements constitute the majority of repeat content. Differences in repeat composition in Decapodiformes and *O. bimaculoides* likely reflects variation in genome architecture associated with interaction conservation and lineage specificity.

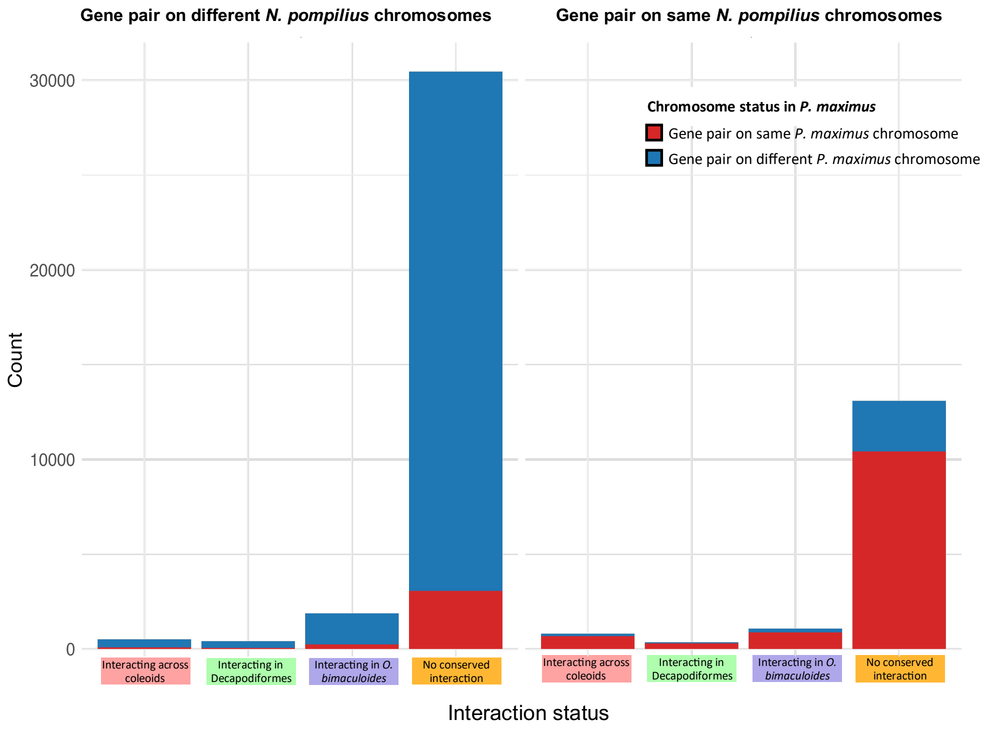


**Figure S7. Chromosomal arrangement of interacting gene pairs in *N. pompilius* and *P. maximus*.**
Barplots showing the number of gene pairs grouped by interaction status (x-axis), split by whether they are located on the same (right) or different (left) *N. pompilius* chromosomes. Bars are coloured by whether the gene pairs are on the same (red) or different (blue) *P. maximus* chromosomes, which is largely consistent with their chromosomal arrangement in *N. pompilius*.

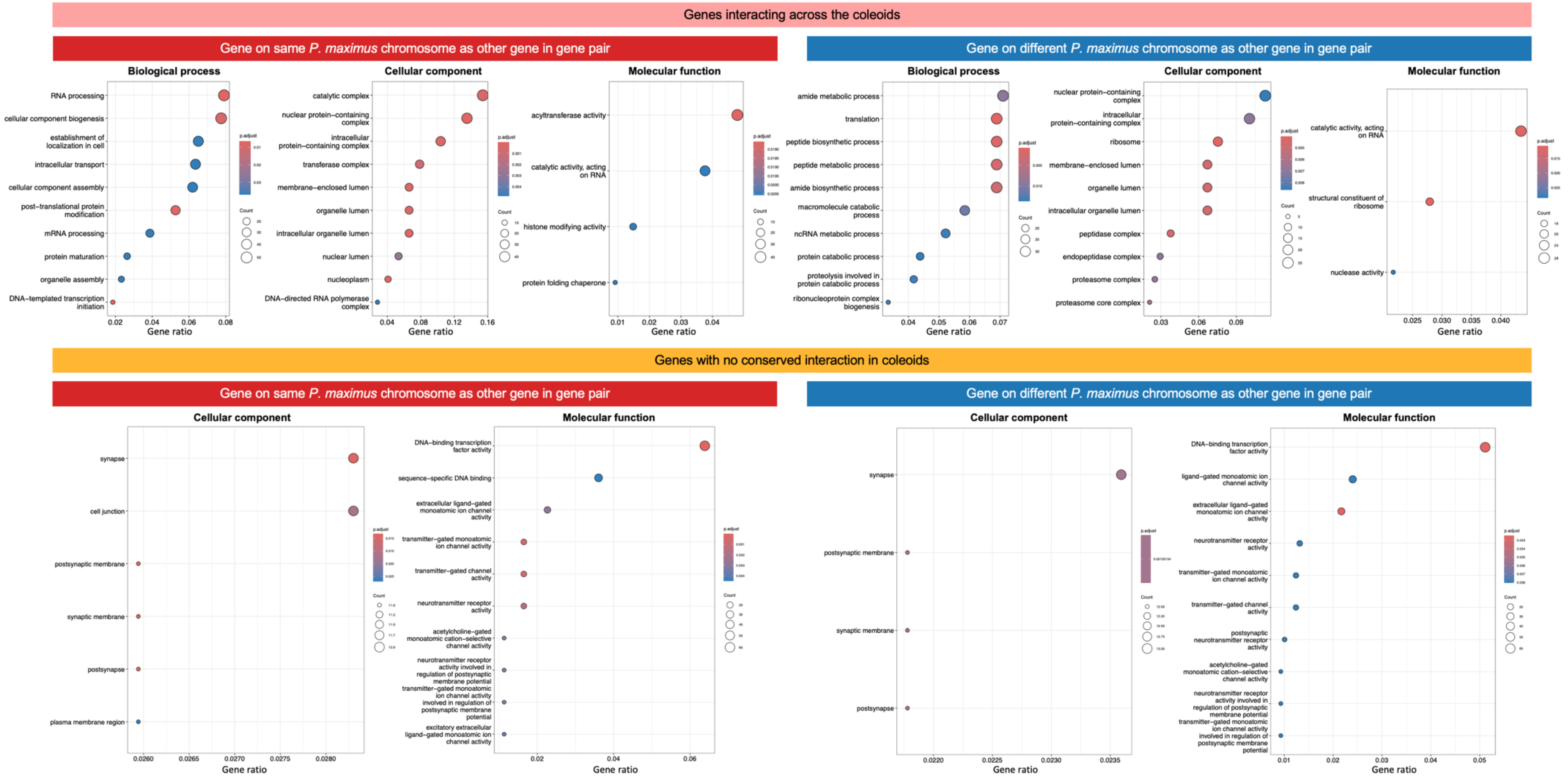


**Figure S8. Gene Ontology (GO) enrichment analysis for *E. scolopes* genes grouped by interaction status and chromosomal context in *P. maximus*.**
Dot plots show enriched GO terms (biological process, cellular component, and molecular function) for genes interacting across the coleoids (top panels; pink) and for genes with no conserved interaction in coleoids (bottom panels; orange). Genes are further separated by whether their gene pairs are located on the same (left; red) or different (right; blue) *P. maximus* chromosome as the other gene in the pair. For each group, enriched GO terms are displayed along the y-axis. Dot size represents the number of genes annotated with the term; colour reflects statistical significance (adjusted *p*-value); and the x-axis shows the gene ratio, that is, the proportion of input genes annotated with the given GO term.

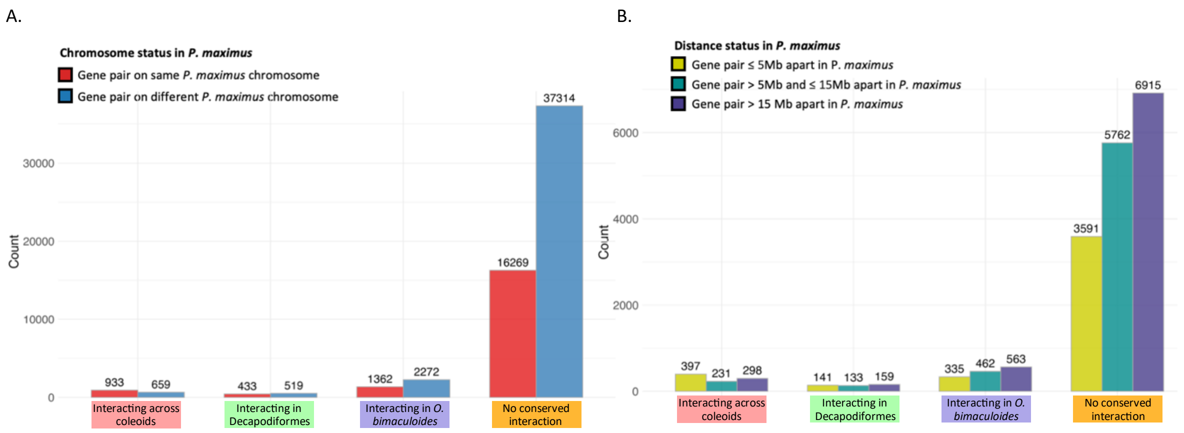


**Figure S9. Chromosomal and genomic distance context of gene pairs stratified by interaction status.** Bar plots showing the number of gene pairs in each interaction category, coloured by their chromosomal arrangement in *P. maximus*. Gene pairs are categorised as being located as (A) on the same (red) or different (blue) *P. maximus* chromosomes or (B) ≤ 5 Mb (yellow), > 5 Mb and ≤ 15 Mb (green), and > 15 Mb (purple) apart on the same *P. maximus* chromosomes.

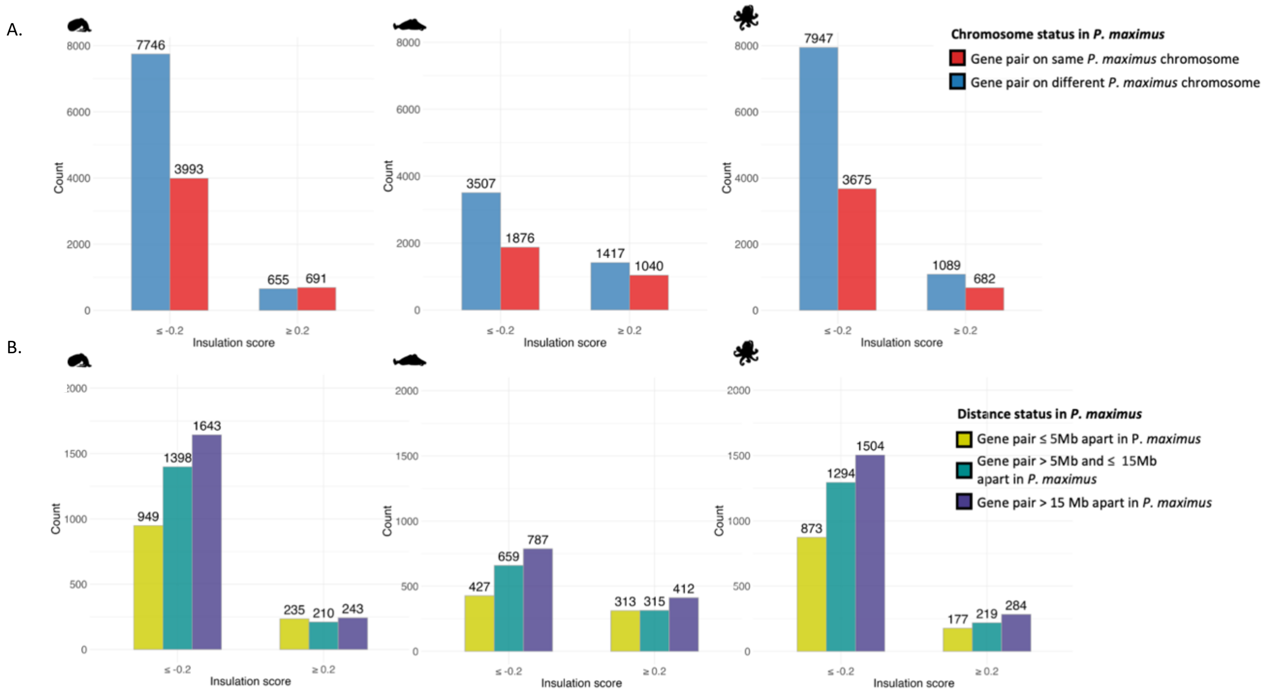


**Figure S10. Chromosomal and genomic distance context of gene pairs stratified by insulation score and species.**

(A) Bar plots showing the number of gene pairs with low (≤ −0.2) or high (≥ 0.2) average insulation scores in *E. scolopes*, *S. officinalis*, and *O. bimaculoides*, coloured by their chromosomal arrangement in *P. maximus*. Gene pairs are categorised as being located on the same (red) or different (blue) *P. maximus* chromosomes. Insulation scores were computed from Micro-C data at 100 kb resolution using a 350 kb window in *S. officinalis* and at 50 kb resolution with a 350 kb window in *O. bimaculoides*

(B) Bar plots showing gene pairs in each coleoid species located on the same *P. maximus* chromosome, grouped by their linear genomic distance in *P. maximus*, coloured by distance category: ≤ 5 Mb (yellow), > 5 Mb and ≤ 15 Mb (green), and > 15 Mb (purple).

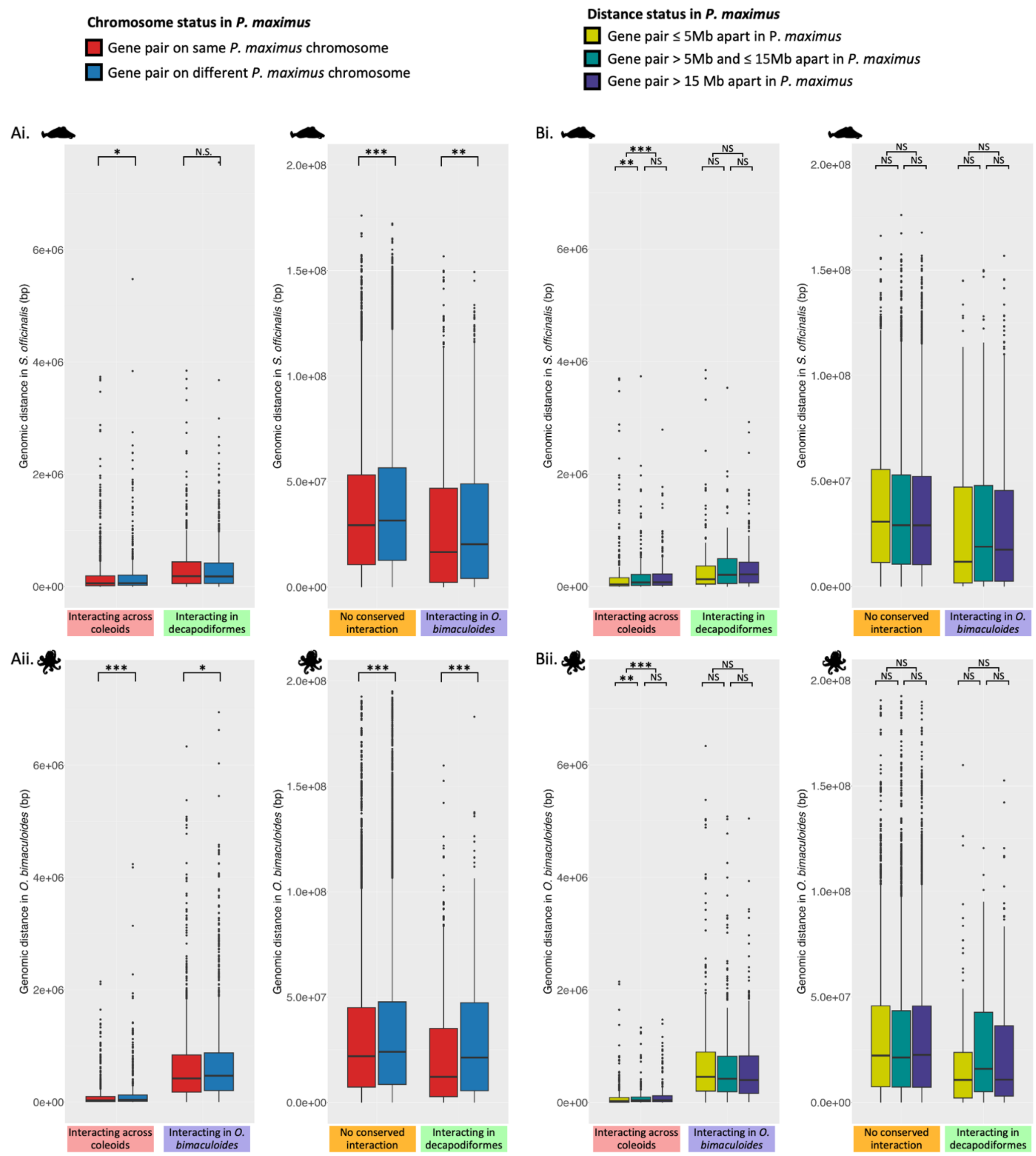


**Figure S11. Genomic distances between gene pairs in *S. officinalis* and O. *bimaculoides*, grouped by interaction status and ancestral genomic context in *P. maximus*.**
Boxplots show genomic distances between gene pairs in *S. officinalis* (top) *O. bimaculoides* (bottom) grouped by interaction status (x-axis). Panels in (A) are coloured by ancestral chromosome status in *P. maximus* (same chromosome: red; different chromosomes: blue). Panels in (B) are coloured by ancestral genomic distance in *P. maximus* (≤5 Mb: yellow; >5–15 Mb: turquoise; >15 Mb: purple). Boxplot significance values are based on BH corrected Wilcoxon rank-sum tests: ** = $p < 0.05$, ** = $p < 0.005$, *** = $p < 0.0005$.

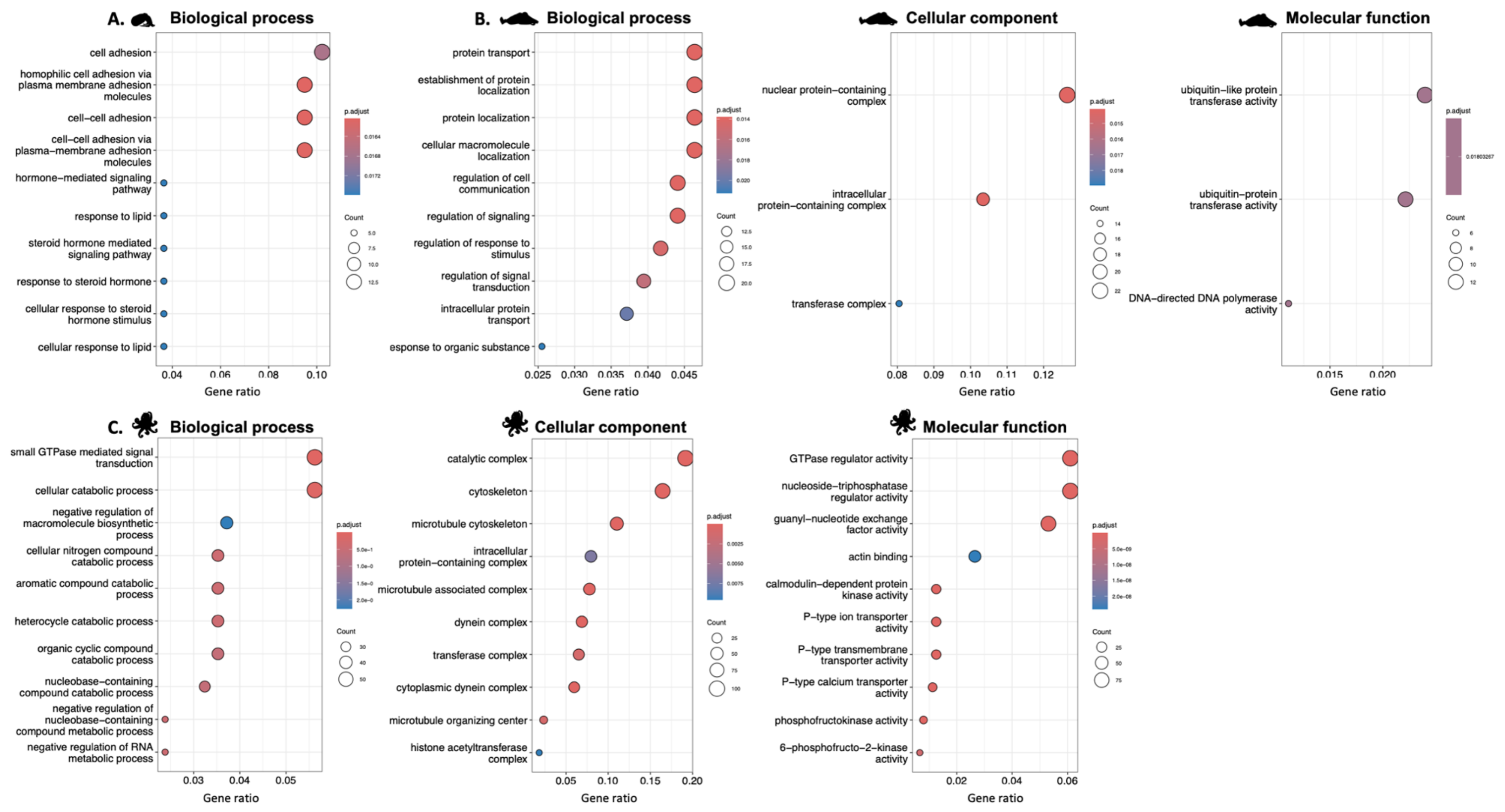


**Figure S12. GO term enrichment results for genes located in loop anchors across whole-embryo datasets in *E. scolopes*, *O. bimaculoides*, and *S. officinalis*.** (A) Enriched Gene Ontology (GO) terms for loop anchor genes in *E. scolopes*, (B) *O. bimaculoides*, and (C) *S. officinalis*, grouped by GO category: biological process, cellular component, and molecular function. Each dot represents an enriched GO term, dot size indicates the number of genes associated with each term, and colour reflects statistical significance (adjusted *p*-value). Gene ratio is defined as the number of genes in the query set annotated with a given term divided by the total number of query genes. Only genes located at loop anchors in loops detected at 100 kb and 50 kb resolution were included in the analysis.

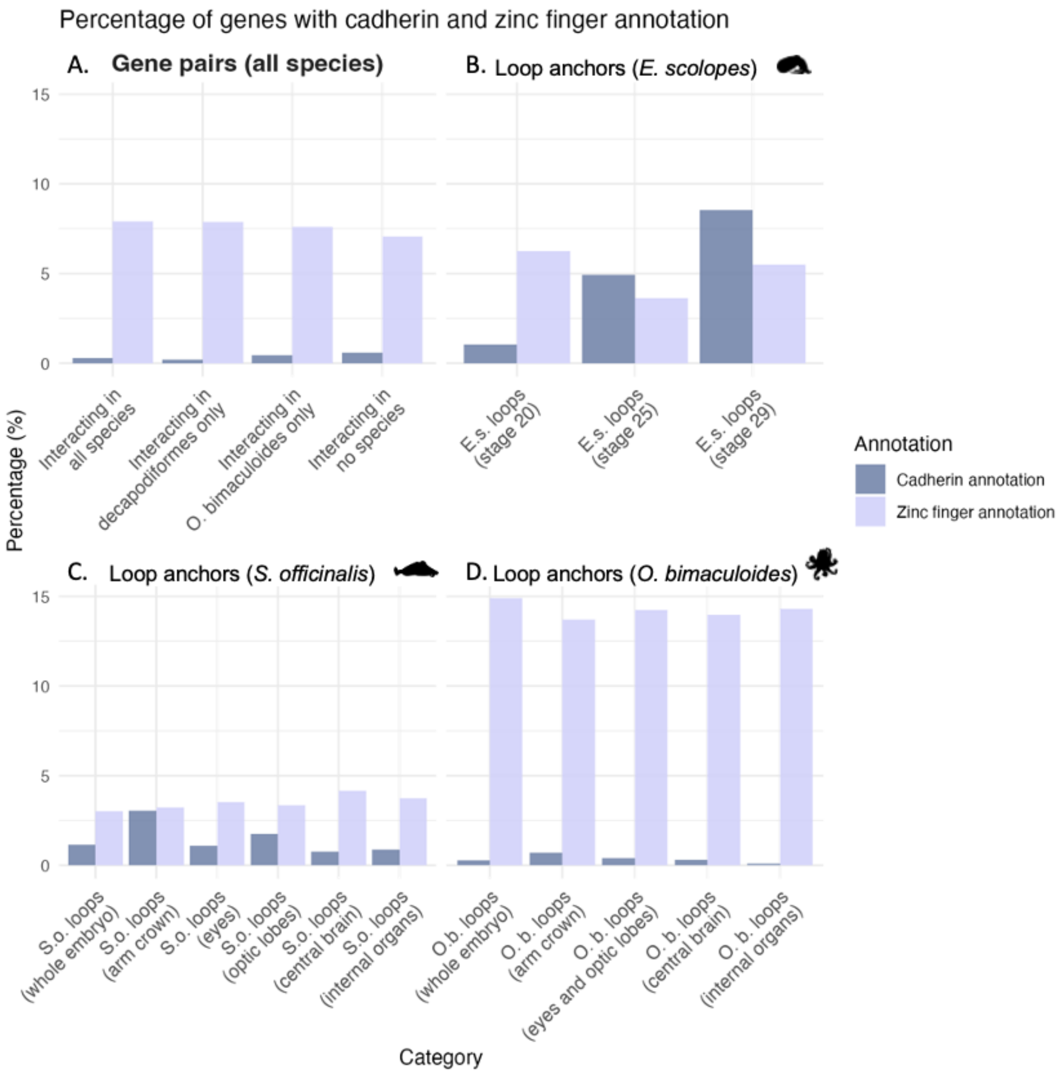


**Figure S13. Percentage of genes with cadherin and zinc finger annotations across interaction and loop anchor categories.**

Barplots showing the percentage of genes annotated with cadherin (purple) or zinc finger (light blue) domains across (A) gene pair categories and loop anchors in (B) *E. scolopes* (B) *S. officinalis* and (C) *O. bimaculoides*.

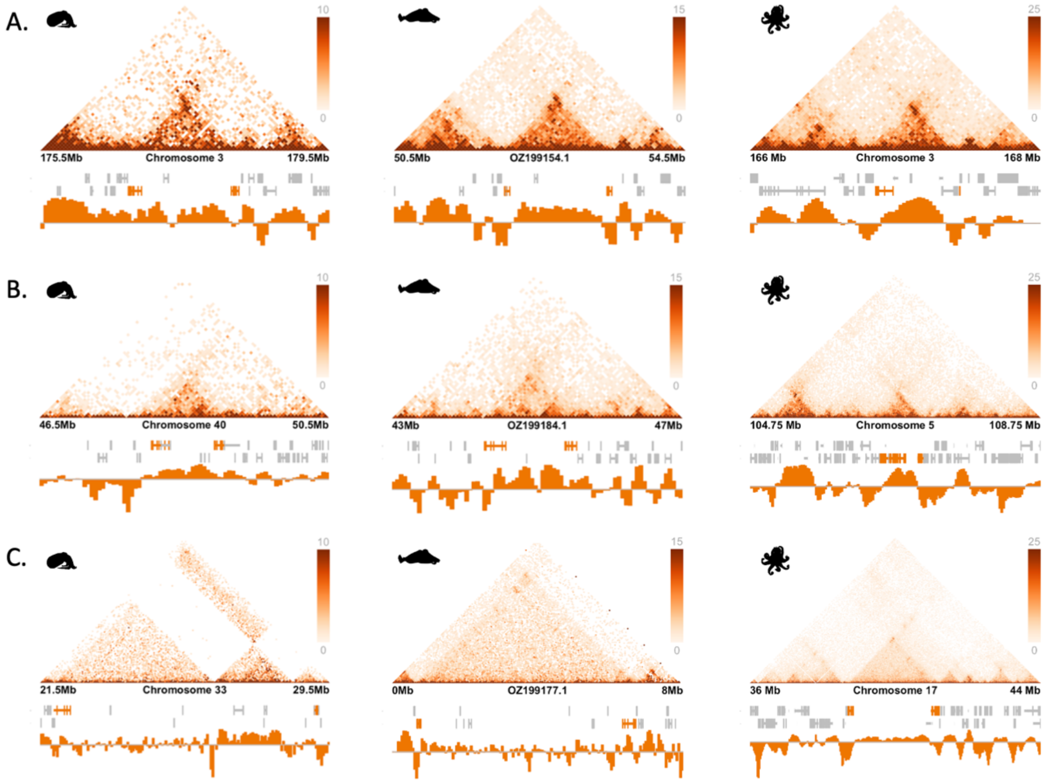


**Figure S14. Examples of loops conserved across *E. scolopes*, *S. officinalis*, and *O. bimaculoides*.**
Micro-C contact matrices and insulation score profiles are shown for three conserved loops (A–C), each detected in all three species, identified at 100 kb and 50 kb resolution. Loops are shown at 50 kb resolution for *E. scolopes* (left) and *S. officinalis* (middle), and at 25 kb resolution *for O. bimaculoides* (right). Orange bars represent genes that overlap with loop anchors. Grey bars represent other genes in the depicted region. Orange tracks show insulation scores calculated using a 350 kb window. Loops were considered conserved if they contained at least one orthologous gene at both the start and end positions, regardless of gene orientation. Note that the *E. scolopes* region in panel C shows a localised signal dropout between loop anchors, likely reflecting a small misassembly. Nonetheless, orthologous genes are present at both ends, supporting conservation of the loop across species.

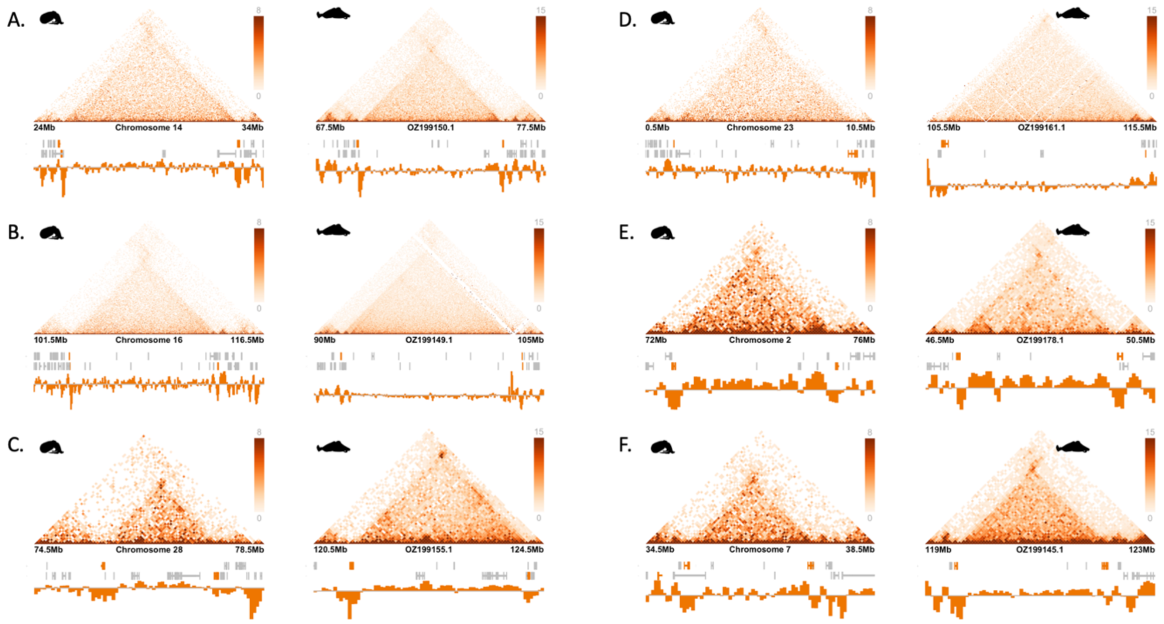


**Figure S15. Examples of loops conserved between *E. scolopes* and *S. officinalis* with disrupted anchor organisation in *O. bimaculoides*.**

Micro-C contact matrices and insulation score profiles are shown for six loops (A–F) conserved between *E. scolopes* (left) and *S. officinalis* (right), identified at 100 kb and 50 kb resolution. Loops are displayed at 50 kb resolution for both species. Orange bars indicate genes overlapping loop anchors; grey bars show other genes in the region. Orange tracks represent insulation scores calculated using a 350 kb window. Panels A–C show loops whose anchor orthologues are located on different chromosomes in *O. bimaculoides*. Panels D–F show loops whose anchor orthologues are located far apart on the same chromosome in *O. bimaculoides.*

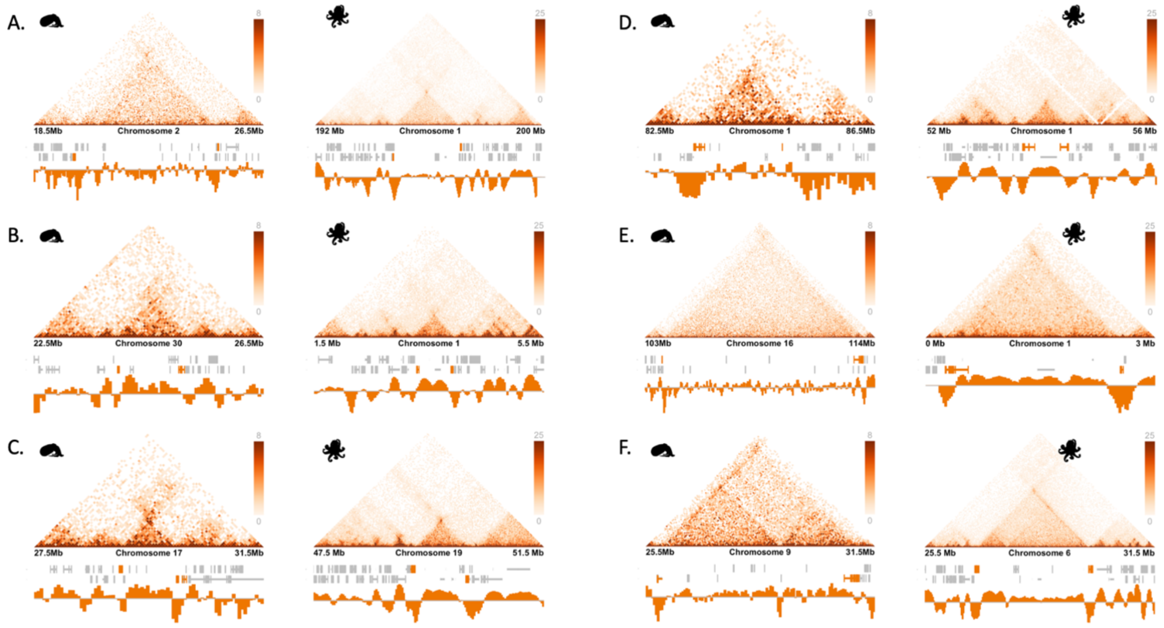


**Figure S16. All loops conserved between *E. scolopes* and *O. bimaculoides*.**
Micro-C contact matrices and insulation score profiles are shown for six loops (A–F) conserved between *E. scolopes* and *O. bimaculoides*, identified at 100 kb and 50 kb resolution. Loops are displayed at 50 kb resolution for *E. scolopes* and 25 kb resolution for *O. bimaculoides*. Orange bars indicate genes overlapping loop anchors; grey bars show other genes in the surrounding region. Insulation scores, calculated using a 350 kb window, are shown as orange tracks beneath each matrix. All conserved loops are shown in this figure, defined as those with at least one orthologous gene at both anchor positions, regardless of gene orientation.

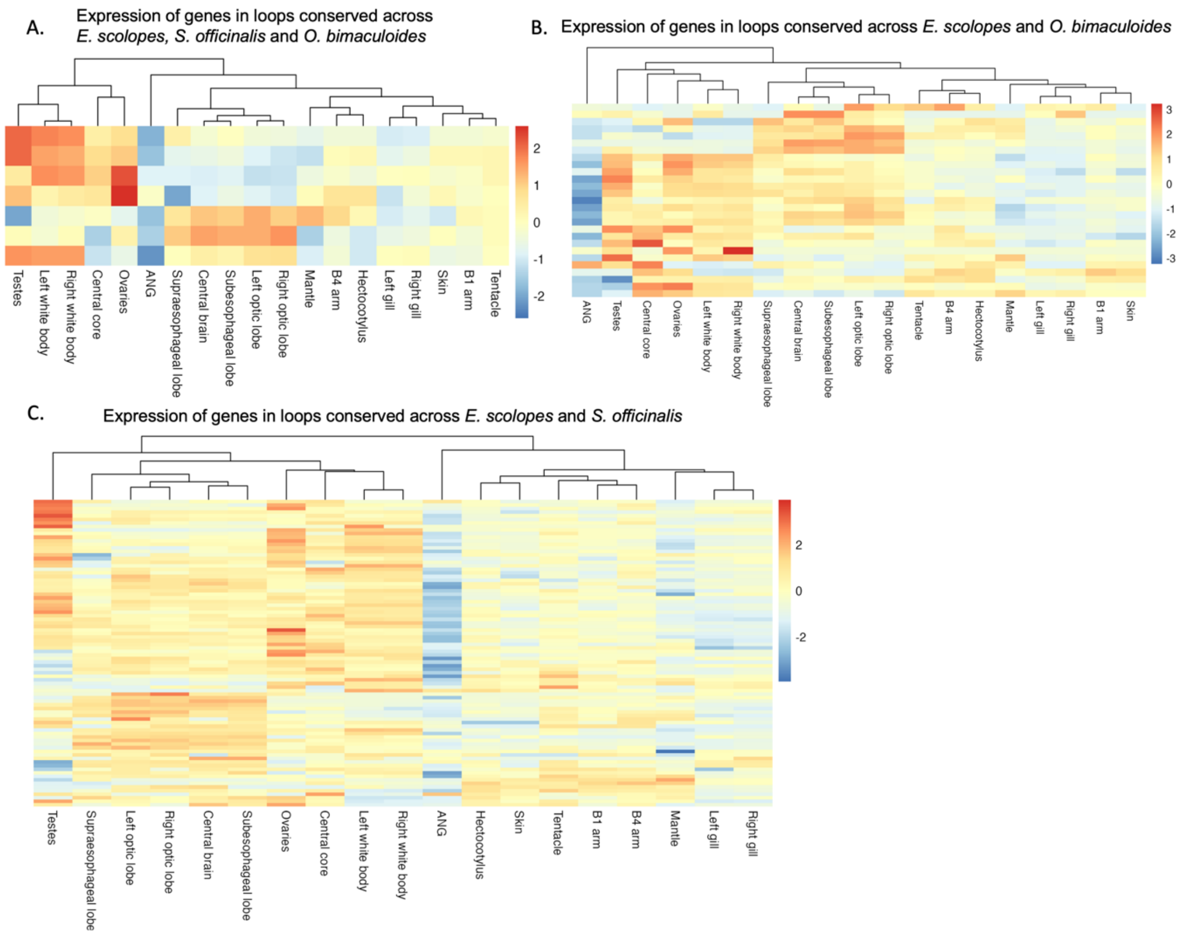


**Figure S17. Expression of genes located in conserved loops across *E. scolopes* tissues.**
Heatmaps showing log-transformed TPM expression of *E. scolopes* genes located in loops conserved across. (A) *E. scolopes*, *S. officinalis*, and *O. bimaculoides*; (B) *E. scolopes* and *O. bimaculoides*; (C) *E. scolopes* and *S. officinalis*. Each row represents a gene and each column a tissue. Expression values are scaled by row (Z-score) to highlight tissue-specific expression patterns. Hierarchical clustering was applied to both genes and tissues using Euclidean distance and complete linkage.

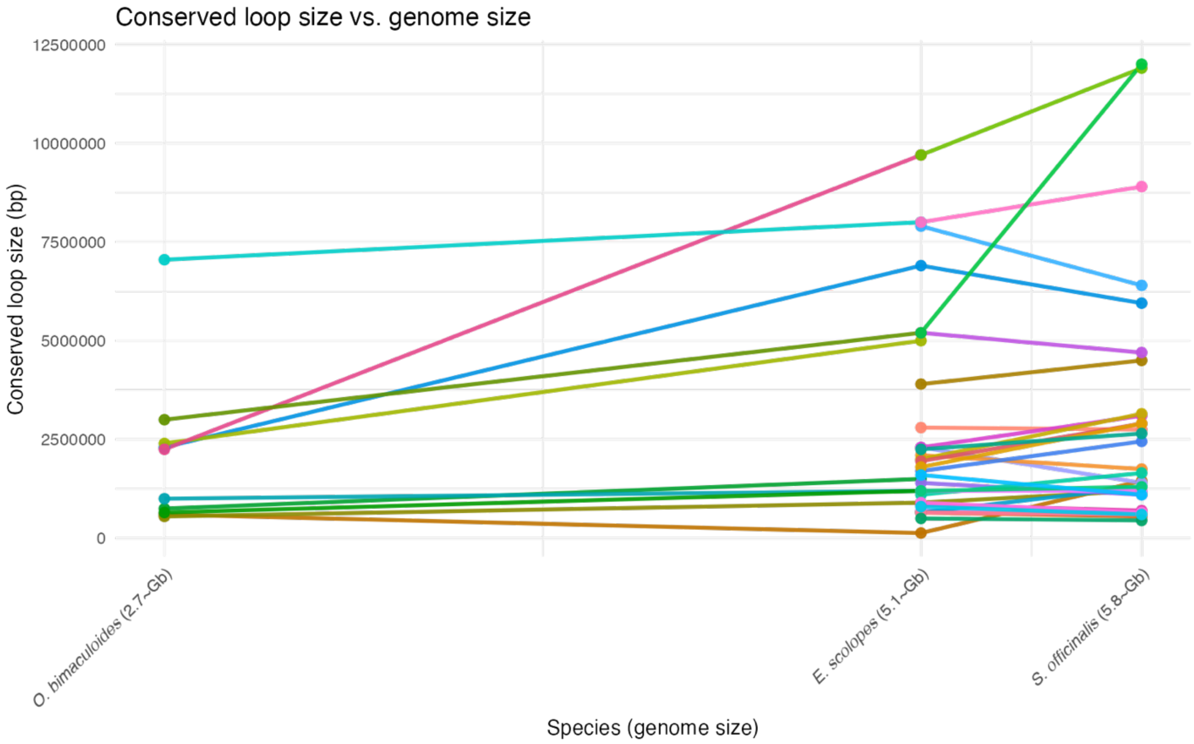


**Figure S18. Conserved chromatin loop sizes across species with varying genome sizes.**
Each line and colour represents a conserved loop shared between two or more coleoid species, with loop size plotted on the y-axis and species (ordered by genome size) on the x-axis. Genome sizes are indicated in parentheses on the x-axis labels.

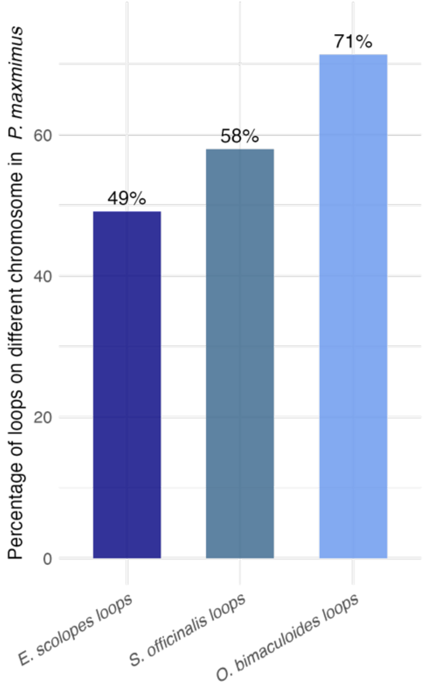


**Figure S19. Percentage of loops in each coleoid species with orthologous genes located on more than one chromosome in *P. maximus*.** Bar plot showing the proportion of chromatin loops in *E. scolopes*, *S. officinalis*, and *O. bimaculoides* in which the genes located at loop anchors map to different chromosomes in *P. maximus*. This serves as a proxy for the ACCRE, with higher percentages indicating greater disruption of ancestral synteny.

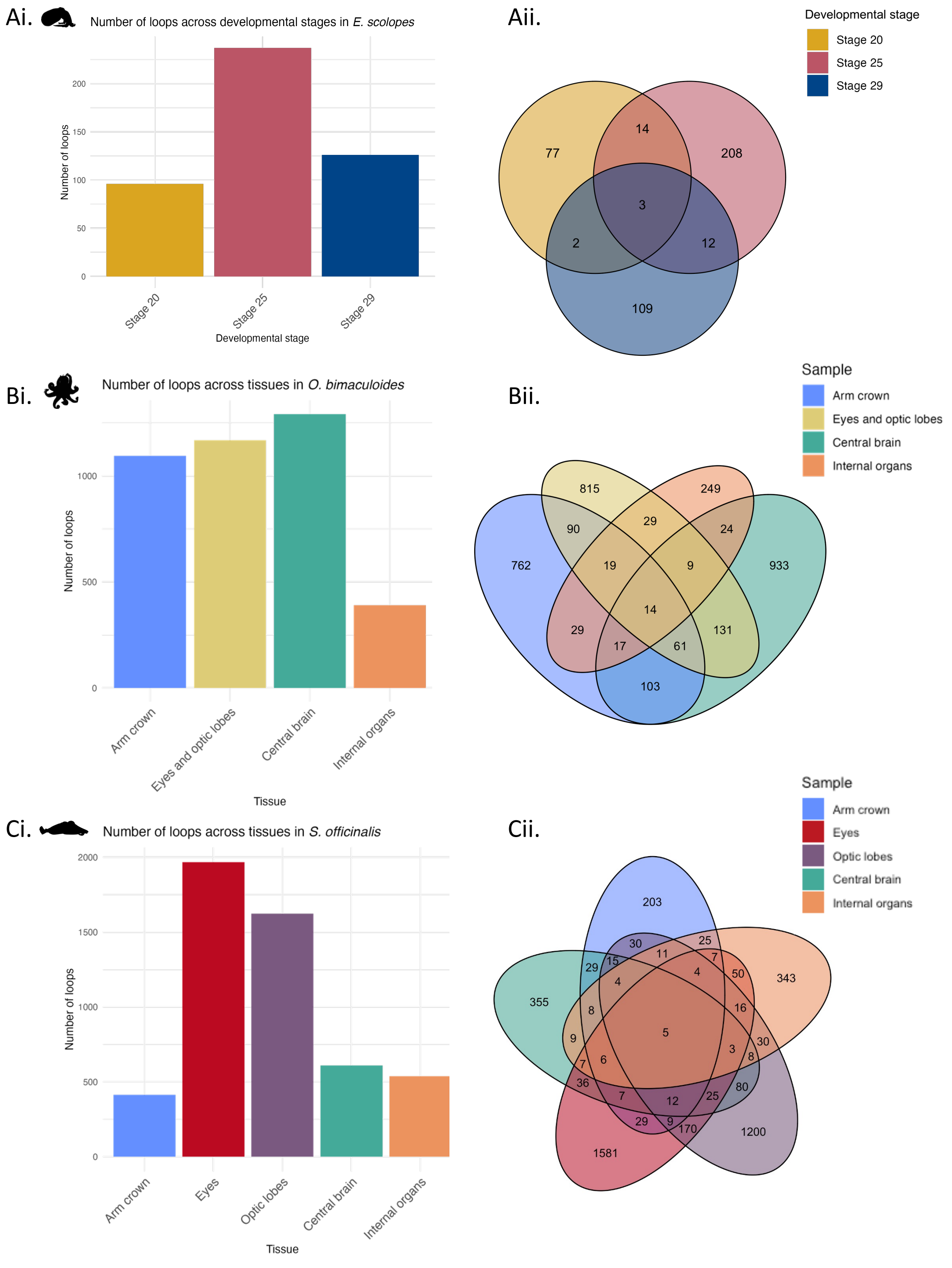


**Figure S20. Number and overlap of chromatin loops across developmental stages and tissues in *E. scolopes*, *O. bimaculoides*, and *S. officinalis*.** (Ai) Number of chromatin loops identified in *E. scolopes* across three developmental stages (stage 20, 25, and 29), and (Aii) Venn diagram showing the overlap of loops across these stages.

(Bi) Number of loops detected in four stage 20 tissues in *O. bimaculoides*: arm crown, eyes and optic lobes, central brain, and internal organs and (Bii) Venn diagram showing the overlap of loops across these tissues. (Ci) Number of loops detected in five stage 29 tissues in *S. officinalis*: arm crown, eyes, optic lobes, central brain, and internal organs and

(Cii) Venn diagram showing the overlap of loops across these tissues. Loops were detected for each species at 100 kb and 50 kb resolution.

Loops are shown here regardless of gene content at their anchors; i.e. loops may or may not contain genes at their anchor positions.

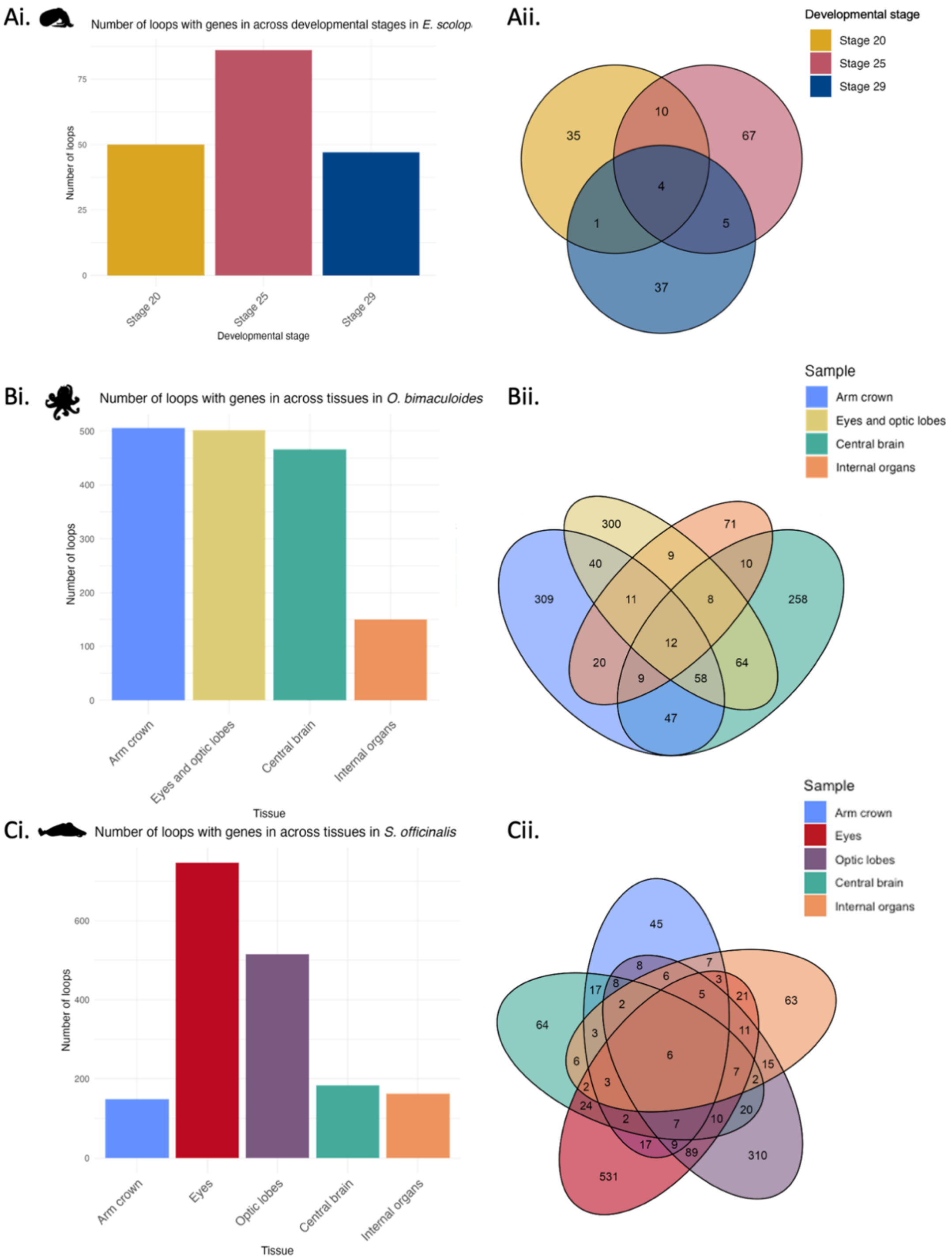


**Figure S21. Number and overlap of loops containing genes across developmental stages and tissues in *E. scolopes*, *O. bimaculoides*, and *S. officinalis*.** (Ai) Number of gene-containing chromatin loops identified in *E. scolopes* across three developmental stages (stage 20, 25, and 29), and (Aii) Venn diagram showing the overlap of loops across these stages.

(Bi) Number of gene-containing loops detected in four stage 20 tissues in *O. bimaculoides*: arm crown, eyes and optic lobes, central brain, and internal organs, and (Bii) Venn diagram showing the overlap of loops across these tissues.

(Ci) Number of gene-containing loops detected in five stage 29 tissues in *S. officinalis*: arm crown, eyes, optic lobes, central brain, and internal organs, and (Cii) Venn diagram showing the overlap of loops across these tissues.

Loops were detected for each species at 100 kb and 50 kb resolution. Only loops with at least one gene in both anchor regions were retained. Loops containing the exact same sets of genes at their anchors were considered duplicates and removed from the analysis.

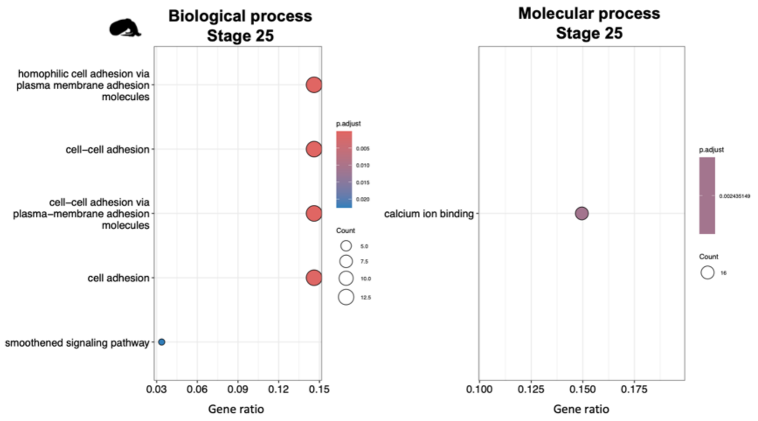


**Figure S22. GO term enrichment results for genes located in loop anchors in developmental stage Micro-C data of *E. scolopes*.** Grouped by GO category: biological process and molecular function. Each dot represents an enriched GO term, dot size indicates the number of genes associated with each term, and colour reflects statistical significance (adjusted *p*-value). Gene ratio is defined as the number of genes in the query set annotated with a given term divided by the total number of query genes. Only genes located at loop anchors in loops detected at 100 kb and 50 kb resolution were included in the analysis.

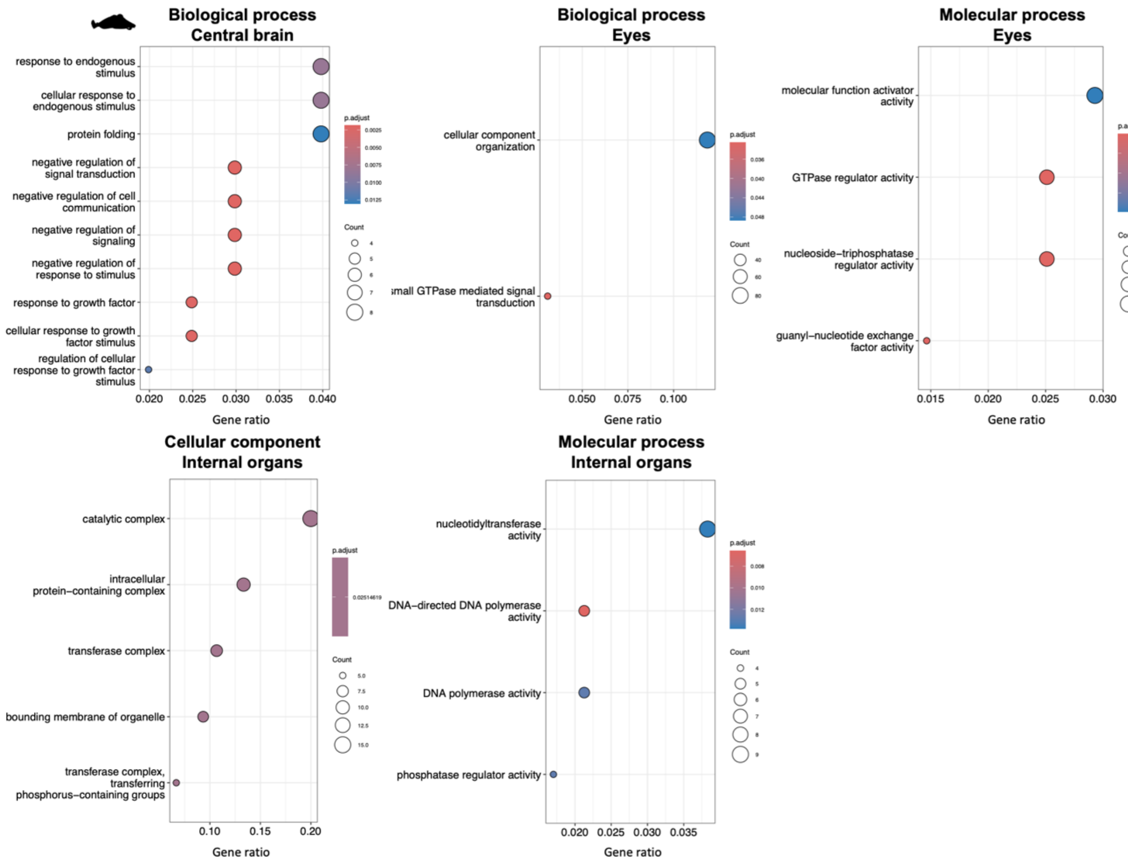


**Figure S23. GO term enrichment results for genes located in loop anchors in tissue-specific data for *S. officinalis.*** Grouped by GO category: biological process, cellular component, and molecular function. Each dot represents an enriched GO term, dot size indicates the number of genes associated with each term, and colour reflects statistical significance (adjusted *p*-value). Gene ratio is defined as the number of genes in the query set annotated with a given term divided by the total number of query genes. Only genes located at loop anchors in loops detected at 100 kb and 50 kb resolution were included in the analysis.

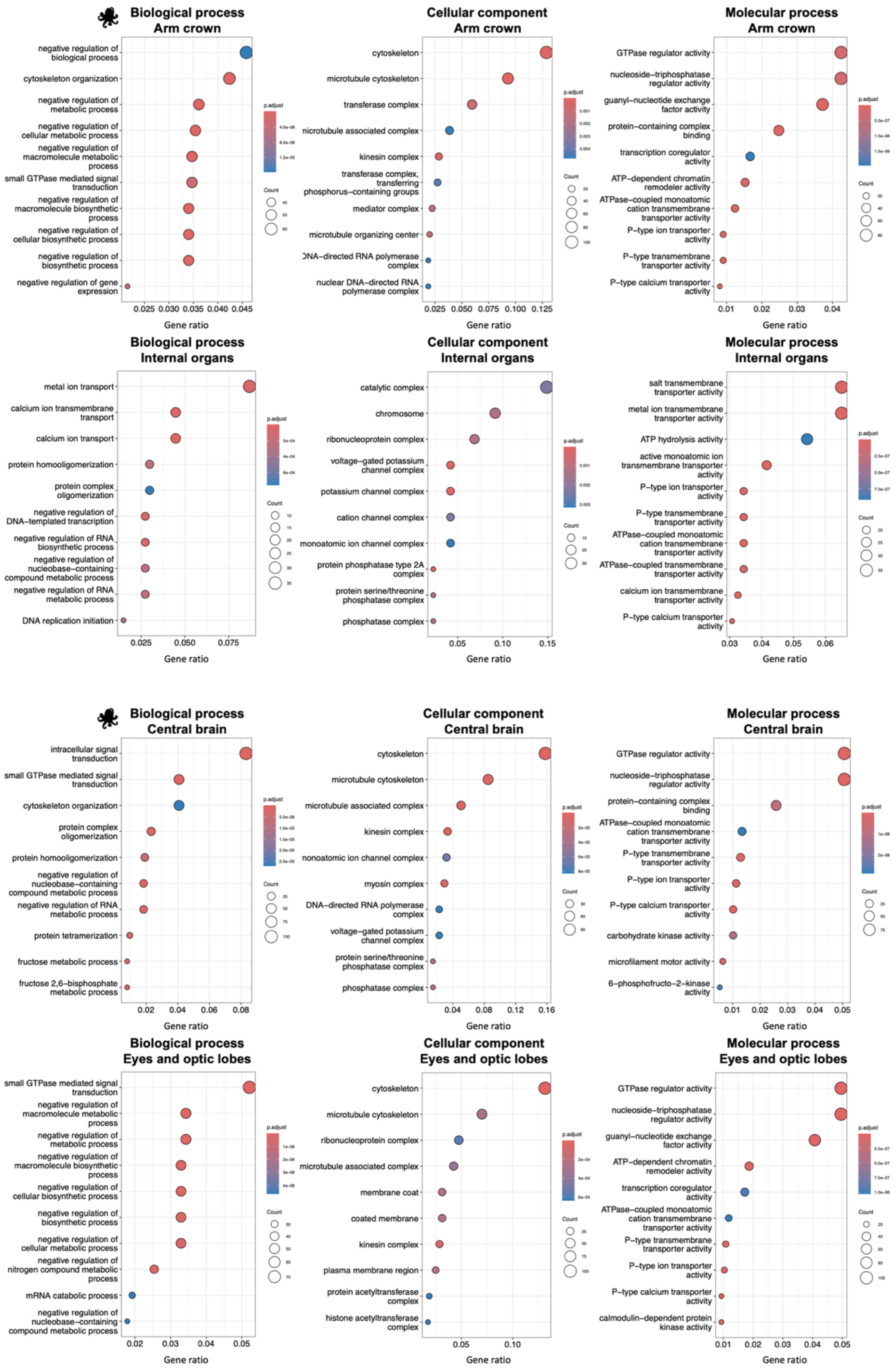


**Figure S24. GO term enrichment results for genes located in loop anchors in tissue-specific data for *O. bimaculoides.*** Grouped by GO category: biological process, cellular component, and molecular function. Each dot represents an enriched GO term, dot size indicates the number of genes associated with each term, and colour reflects statistical significance (adjusted *p*-value). Gene ratio is defined as the number of genes in the query set annotated with a given term divided by the total number of query genes. Only genes located at loop anchors in loops detected at 100 kb and 50 kb resolution were included in the analysis.

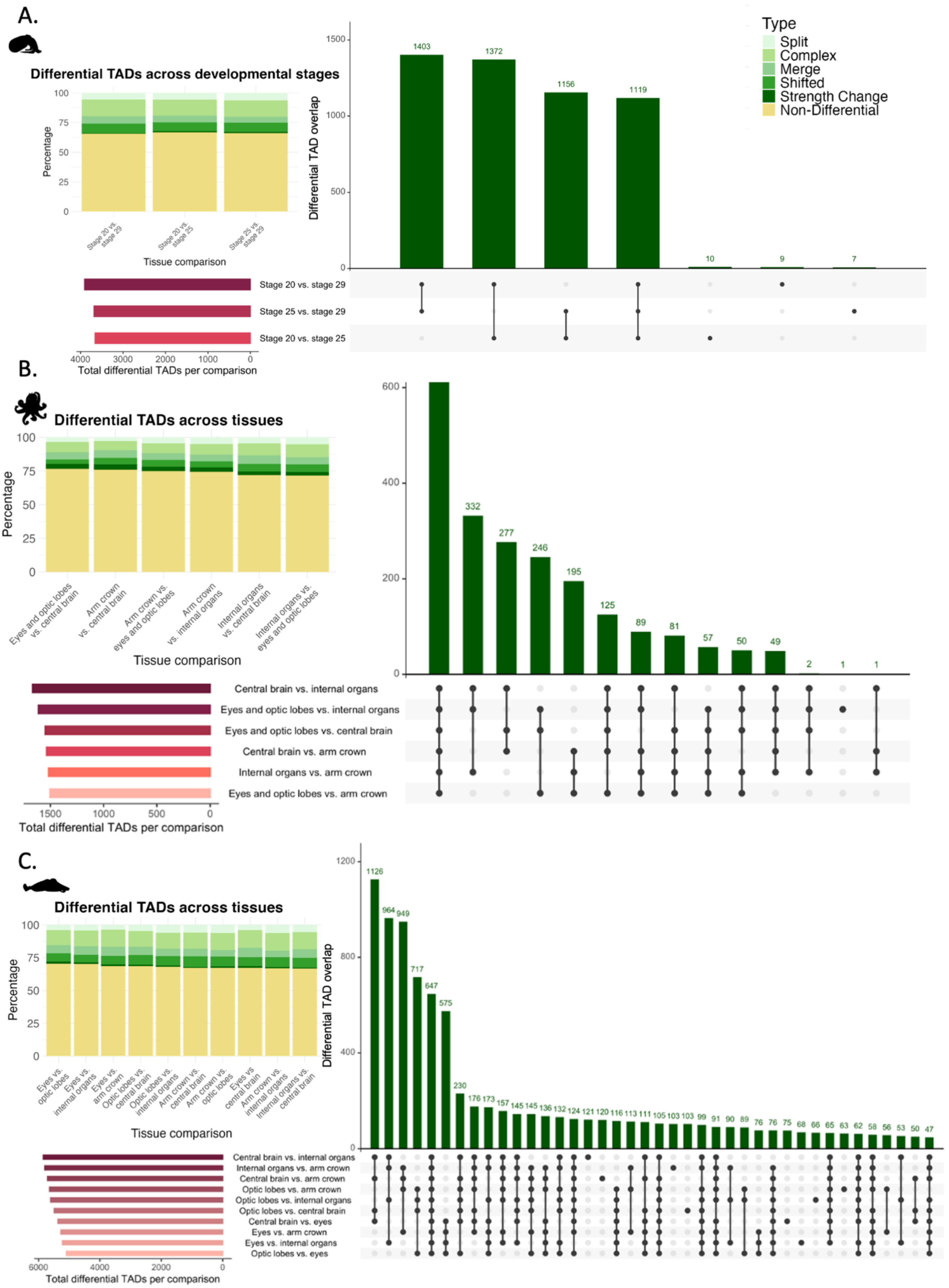
A.
Differential TADs across developmental stages
Percentage
Stage 20 vs. stage 29
Stage 20 vs. stage 25
Stage 25 vs. stage 29
Tissue comparison
Differential TAD overlap
1403
1372
1156
1119
10
9
7
Type
Split
Complex
Merge
Shifted
Strength Change
Non-Differential
Stage 20 vs. stage 29
Stage 25 vs. stage 29
Stage 20 vs. stage 25
Total differential TADs per comparison
B.
Differential TADs across tissues
Percentage
Eyes and optic lobes vs. central brain
Arm crown vs. central brain
Arm crown vs. eyes and optic lobes
Arm crown vs. internal organs
Internal organs vs. central brain
Internal organs vs. eyes and optic lobes
Tissue comparison
332
277
246
195
125
89
81
57
50
49
2
1
1
Central brain vs. internal organs
Eyes and optic lobes vs. internal organs
Eyes and optic lobes vs. central brain
Central brain vs. arm crown
Internal organs vs. arm crown
Eyes and optic lobes vs. arm crown
Total differential TADs per comparison
C.
Differential TADs across tissues
Percentage
Eyes vs. optic lobes
Eyes vs. internal organs
Eyes vs. arm crown
Optic lobes vs. central brain
Optic lobes vs. internal organs
Arm crown vs. central brain
Arm crown vs. optic lobes
Eyes vs central brain
Arm crown vs. internal organs
Internal organs vs. central brain
Tissue comparison
Differential TAD overlap
1126
964
949
717
647
575
230
176
173
157
145
145
136
132
124
121
120
116
113
111
105
103
103
99
91
90
89
76
76
75
68
66
65
63
62
58
56
53
50
47
Central brain vs. internal organs
Internal organs vs. arm crown
Central brain vs. arm crown
Optic lobes vs. arm crown
Optic lobes vs. internal organs
Optic lobes vs. central brain
Central brain vs. eyes
Eyes vs. arm crown
Eyes vs. internal organs
Optic lobes vs. eyes
Total differential TADs per comparison

**Figure S25. Differential compartments across developmental stages and tissues in *E. scolopes*, *O. bimaculoides*, and *S. officinalis*.** (A) Differential compartments identified across developmental stages in *E. scolopes*, (B) across stage 20 tissues in *O. bimaculoides*, and (C) across stage 29 tissues in *S. officinalis*. Left: stacked barplots showing the proportion of differential compartments per comparison by type of change (Split, Merge, Shifted, Complex, Strength Change), with non-differential compartments shown in grey. Middle: total number of differential compartments detected per pairwise comparison. Right: UpSet plots showing shared and unique differential compartments across comparisons. Differential compartments were identified using TADCompare and analysed at 100 kb resolution for *E. scolopes* and *S. officinalis*, and 50 kb resolution for *O. bimaculoides*, using a 1 Mb sliding window. If a compartment contained an NA in the 'Differential' classification in any comparison, it was excluded from all comparisons. compartments were considered overlapping if they shared exact coordinates and were classified as differential in each file. compartments classified as differential in one comparison and non-differential in all others were counted as unique to that comparison.

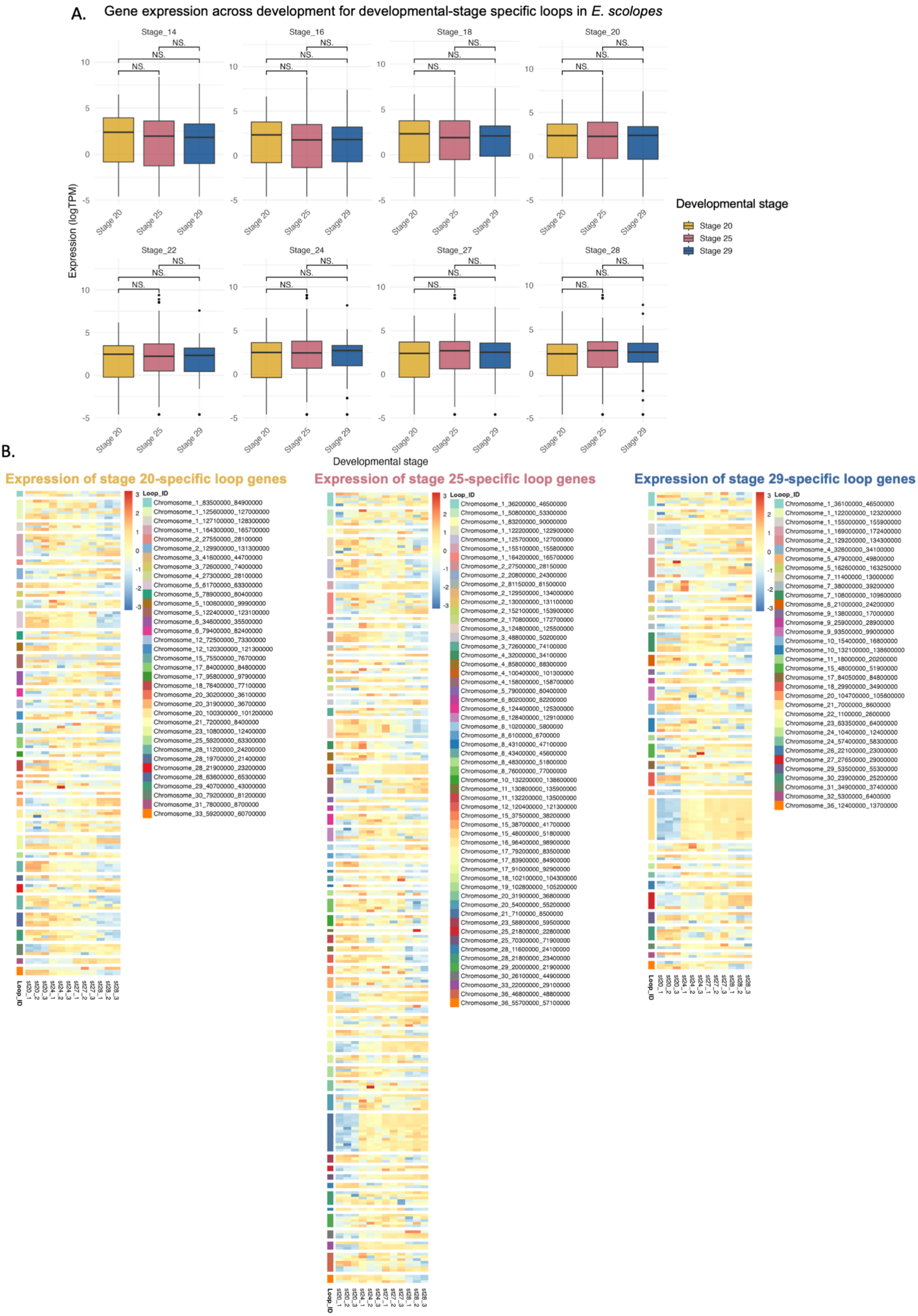
A.
Gene expression across development for developmental-stage specific loops in *E. scolopes*
Stage_14
Stage_16
Stage_18
Stage_20
Stage_22
Stage_24
Stage_27
Stage_28
NS.
Expression (logTPM)
Stage 20
Stage 25
Stage 29
Developmental stage
B.
Expression of stage 20-specific loop genes
Expression of stage 25-specific loop genes
Expression of stage 29-specific loop genes
Loop_ID

**Figure S26. Gene expression patterns across development for *E. scolopes* genes located in developmental stage-specific loops**.
(A) Boxplots showing log-transformed TPM expression across three developmental stages (stage 20, 25, and 29) for genes located in loops specific to each stage. Each panel shows the expression of stage-specific loop genes (indicated along the x-axis) across all three stages, highlighting changes in expression across development. Statistical comparisons were performed using Wilcoxon rank-sum tests; 'NS' indicates non-significant comparisons. Genes found in loops shared across multiple developmental stages were excluded to ensure stage specificity.
(B) Heatmaps showing log-transformed TPM expression of genes located in stage-specific loops across developmental stages: genes in stage 20-specific loops (left), stage 25-specific loops (middle), and stage 29-specific loops (right). Rows represent individual genes, and columns represent three biological replicates per developmental stage. Each heatmap is scaled by row to highlight stage-specific expression patterns. Genes with zero expression (TPM = 0) and no variation across all samples were excluded, and only loops with more than one remaining expressed gene were retained. As in panel A, loops with gene sets shared across multiple developmental stages were excluded.
Loops were detected at 100 kb and 50 kb resolution.

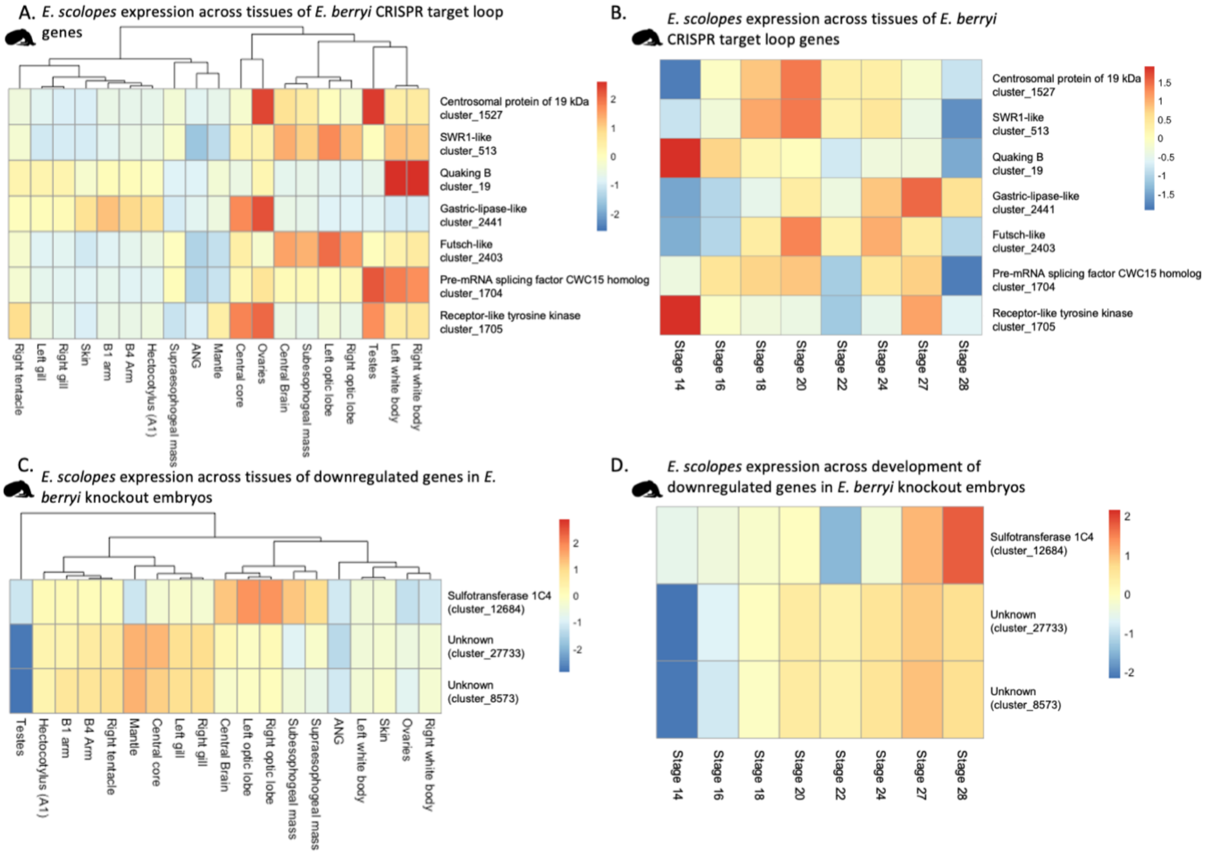


**Figure S27. Expression of *E. scolopes* genes associated with CRISPR-targeted loops and transcriptional response in *E. berryi* knockout embryos.**

(A) Expression across development of *E. scolopes* orthologues located in loops targeted by CRISPR-Cas9 in *E. berryi*.

(B) Expression across tissues of *E. scolopes* orthologues of *E. berryi* CRISPR target genes.

(C) Tissue-wide expression in *E. scolopes* of orthologues of genes significantly downregulated (Wald test, BH correction) in *E. berryi* CRISPR knockout embryos.

(D) Expression across developmental stages of the same significantly downregulated (Wald test, BH correction) ortholog set in *E. scolopes*. Heatmaps display log-transformed TPM values across tissues or developmental stages. Rows represent genes, columns represent biological samples. Expression values are row-scaled to highlight expression patterns across conditions.

Tissues were hierarchically clustered in panels (A) and (C), while developmental stages in panel (B) and (D) were shown in chronological order without clustering to preserve temporal progression. Note: One *E. berryi* gene (EB05908.1) is absent from panels C and D because it is unannotated in the *E. scolopes* gene annotation used in this study, despite having a clear BLAST hit in the region shown in Fig. 5E. This gene is also annotated as G5765 in the *E. scolopes* gene annotation from Rogers et al. (2024).

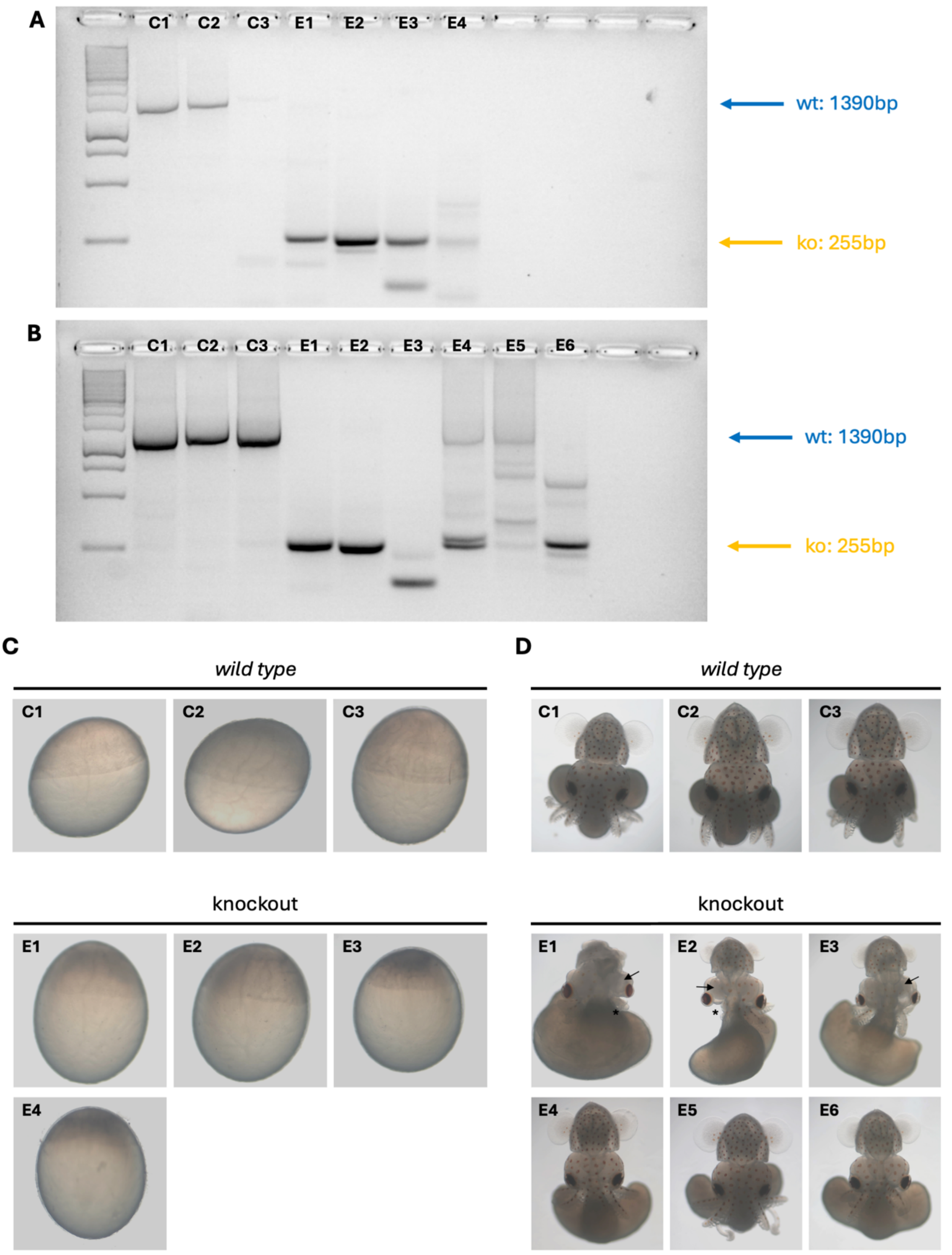


**Figure S28. Phenotypic and genotypic analysis of crispant embryos.**
(A) Agarose gel displaying results of genotyping PCR of embryos depicted in (C). Expected wild type band is 1390 bp (blue arrow) and expected knockout band is 255 bp (orange arrow).
(B) Agarose gel displaying results of genotyping PCR of embryos depicted in (D). Expected wild type band is 1390 bp (blue arrow) and expected knockout band is 255 bp (orange arrow).
(C) Wild type and knockout embryos that were used for RNA sequencing analysis at developmental stage 14.
(D) Wild type and knockout embryos that were used for RNA sequencing analysis at developmental stage 28. Embryos E1-E3 showed significant developmental phenotypes, including smaller optic lobes (arrow) and smaller or missing arms (star), while Embryos E4-E6 showed very slight (developmental delay) to no phenotype after gRNA injection.

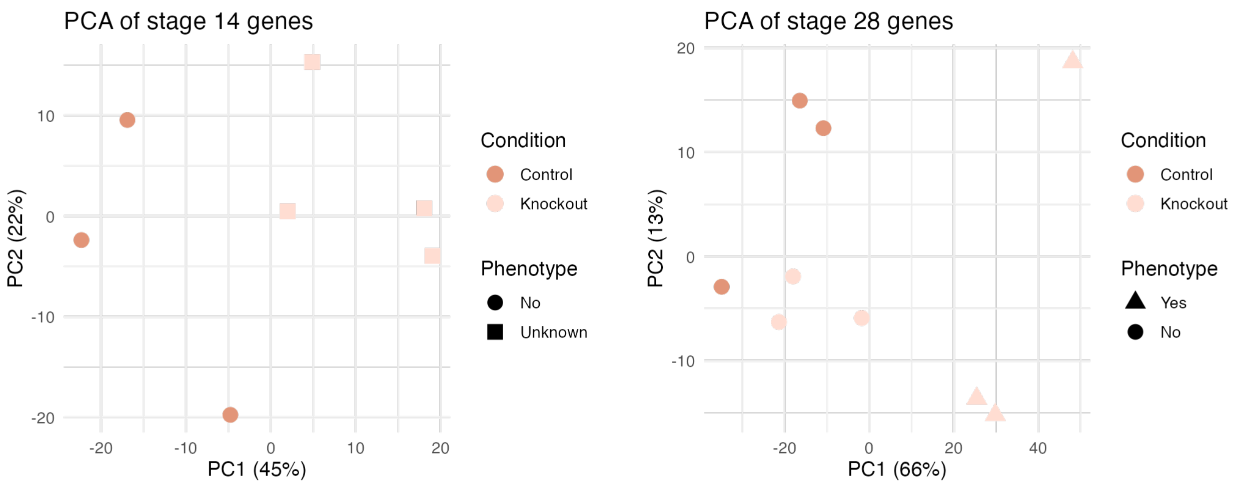


**Figure S29. Principal component analysis (PCA) of *E. berryi* CRISPR embryos at stages 14 and 29.** PCA plots showing transcriptome profiles of *E. scolopes* embryos injected with CRISPR-Cas9 constructs compared to uninjected controls. Analyses were performed separately for stage 14 (left) and stage 29 (right) embryos. Each point represents a single embryo, coloured by experimental condition (control or injected) and shaped by phenotypic outcome (present, absent, or unknown). PCA was performed on variance-stabilised (VST) RNA-seq expression values across all genes expressed at each developmental stage. Axes indicate the percentage of variance explained by each principal component.

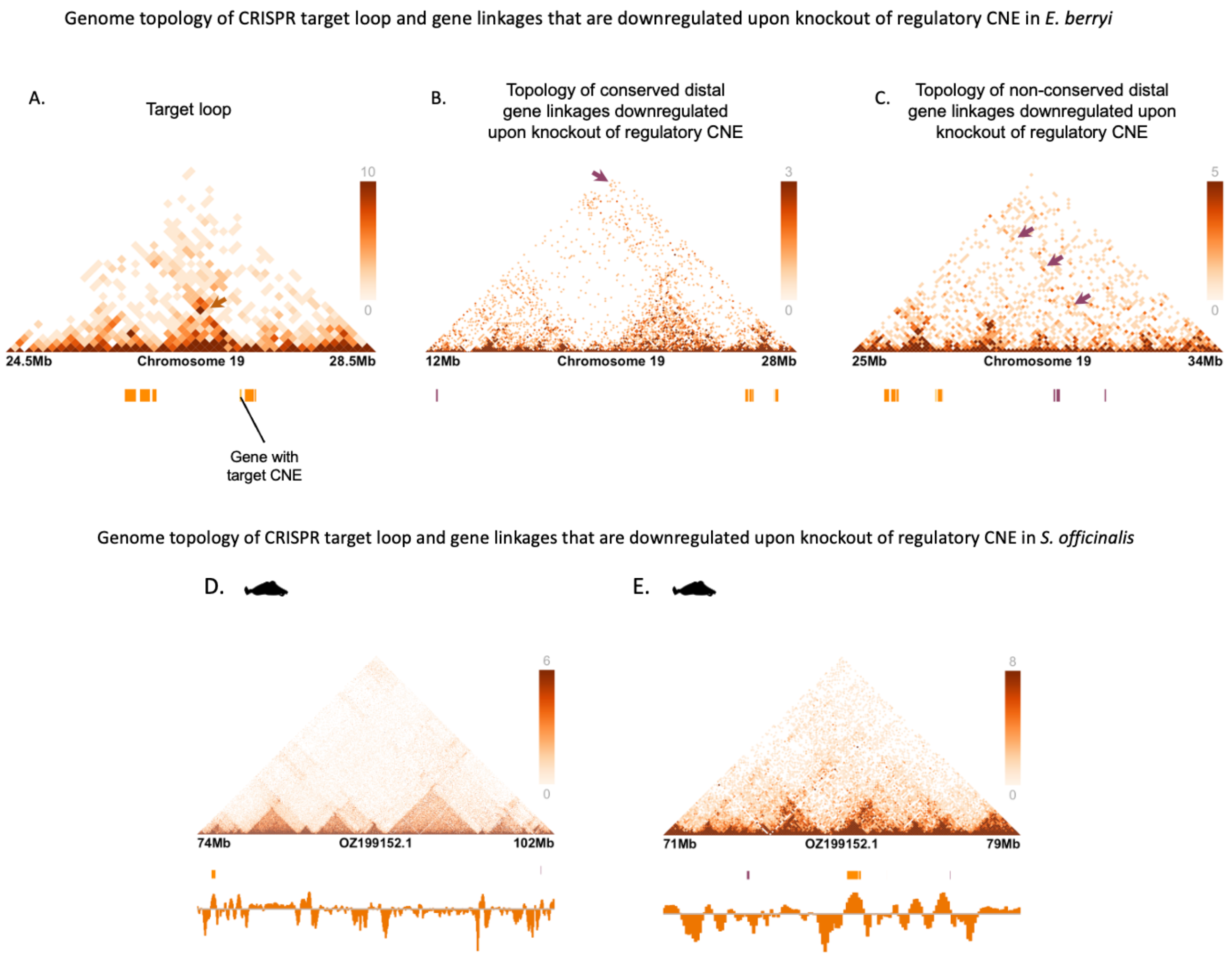


**Figure S30. Genome topology of CRISPR target loop and distal downregulated gene linkages in *E. berryi* and *S. officinalis*.**

(A) Micro-C contact map showing the CRISPR-targeted loop on chromosome 19 in *E. berryi*. The gene containing the targeted putative regulatory non-coding sequence is highlighted in yellow and marked below the map.

(B) *E. berryi* topology of distal gene linkages downregulated upon knockout of the target intron, which is conserved in *E. scolopes* and *O. bimaculoides* (orthologous to the contacts shown in Fig. 5D).

(C) Same as (B), but showing downregulated gene linkages in *E. berryi* that are not conserved in *O. bimaculoides* (but still orthologous to *E. scolopes*, as shown in Fig. 5E).

(D) Micro-C contact map of the corresponding CRISPR target loop in *S. officinalis*. Note that not all genes could be annotated in this species; only cluster_1527/EB05998, cluster_2403/EB06550, cluster_1704/EB06007, and cluster_1705/EB06081 were confidently mapped.

(E) *S. officinalis* topology of distal gene linkages orthologous to downregulated genes in *E. berryi*, conserved across *E. scolopes* and *O. bimaculoides* (as in Fig. 5D). Note that the loop appears only partially conserved in *S. officinalis* due to limitations in genome annotation: *E. scolopes* genes were mapped onto the genome and only top-ranked mRNA hits were retained to create the gene annotation, therefore, some genes are missing. Furthermore, the target gene Quaking B itself is located on an unplaced scaffold in *S. officinalis.*

Contact maps for *E. berryi* are shown at 100 kb resolution, and for *S. officinalis* at 50 kb resolution. Genes within the CRISPR target loop are highlighted in orange below the contact map, with the gene with the target intron shown in yellow. Genes that were downregulated in *E. berryi* knockout embryos are marked in purple. Background genes were removed from the contact map gene annotations to improve clarity.

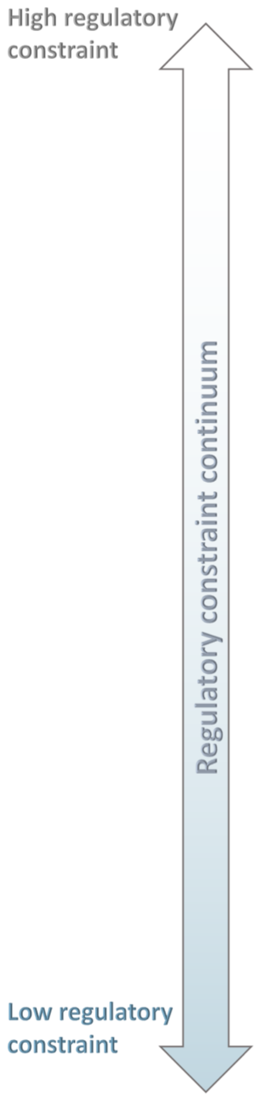


| 3D chromatin structure | Interaction type | Evolutionary trajectory | Features | Function |
|---|---|---|---|---|
| 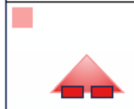 | **Interacting gene pairs conserved across all coleoids and *P. maximus*** | Emerged >600 MYA and conserved since early molluscan ancestor | Very genomically close (~56 kb), constraint against repeat-driven expansion, twice as likely to be located within a compartment across all species than spanning a compartment boundary, highest co-expression (0.474), highest expression, broadly expressed (τ = 0.546), in areas of high gene density (35% coverage in *E. scolopes*, 44% coverage in *O. bimaculoides*) | Enriched for housekeeping functions including ribosome biogenesis, RNA processing, translation, and peptide biosynthesis |
| 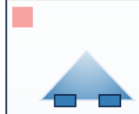 | **Interacting gene pairs conserved across the coleoids only** | Emerged after or due to ancient genome rearrangement event ~450–270 MYA | Genomically close (~70 kb), constraint against repeat-driven expansion, slightly more likely to be located across a compartment boundary than within a compartment, high co-expression (0.405), slightly less highly expressed In some tissues as conserved coleoid interactions from the same ancestral chromosome, broadly expressed (τ = 0.556), high gene (42% coverage in *E. scolopes*, 46% coverage in *O. bimaculoides*) | Involved in broad regulatory functions; enriched for ubiquitin-dependent protein catabolism, proteasome activity, and catalytic complexes; expression highest in brain and reproductive tissue |
| 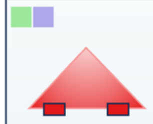 | **Interacting gene pairs conserved in Decapodiformes and/or *O. bimaculoides* and *P. maximus* but lost in the other coleoid lineage** | Suggests lineage-specific retention or loss of ancestral interactions; emerged >600–100 MYA, with asymmetric loss | Moderately spaced (~170 kb), within compartments in the species they are conserved, on boundaries or unstructured in species where lost, broadly expressed (τ = 0.591), high co-expression (0.455 in Decapodiformes), moderate intervening gene density (43% coverage in *E. scolopes*, 46% coverage in *O. bimaculoides*) | Reflect decaying or rewired ancestral interactions; in Decapodiformes: enriched for protein degradation and proteasome-related functions, in *O. bimaculoides*; enriched for biosynthesis, protein folding, translation |
| 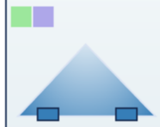 | **Interacting gene pairs present only in Decapodiformes or only in *O. bimaculoides*** | Emerged <270 MYA, potentially post-rearrangement | Moderately spaced (~172 kb, similar/slightly further than above), within compartments in the species they are conserved, on boundaries or unstructured in species where lost, lower insulation, intermediate co-expression (0.377, Decapodiformes), broadly expressed (τ = 0.567), moderate intervening gene density (42% coverage in *E. scolopes*, 46% coverage *in O. bimaculoides*) | Reflect lineage-specific regulatory innovations; Decapodiformes: modest enrichment for regulatory maintenance; expression in *O. bimaculoides*; enriched for cytoskeletal organisation, GTPase signalling, and neurodevelopment |
| 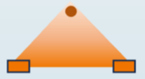 | **Chromatin loops conserved across coleoid species** | Emerged ~450-270 MY–present, not conserved in ancestral molluscs, rare, impacted heavily by species-specific genome expansions | Rare, scale with genome size, often contain conserved internal topologies (e.g., volcano, double loops), high expression in tissues associated with coleoid innovations e.g. optic lobes, central brain, white body | Rare but functionally constrained structures; enriched for chromatin regulation, transcription, and epigenetic remodelling; may support stable regulatory interactions underlying morphogenesis and cell identity |
| 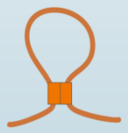 | **Species-specific chromatin loops** | Emerged <100 MYA; structure heavily influenced by clade-specific rearrangements (e.g. chromosomal fusions in Octopodiformes) | Generally large but variable loop sizes, especially in *O. bimaculoides* (*E. scolopes* = 1.6Mb, *S. offiicinalis* = 1.3 Mb, *O. bimaculoides* = 2.9 Mb), often span genes from different ancestral chromosomes suggesting they are formed via chromosomal fusions, depleted of intervening genes (19% coverage in *E. scolopes*, 23% coverage *in O. bimaculoides*), but CNEs, ATAC-seq peaks, and TF binding motifs enriched at anchor points | Mediate clade-specific regulatory programs in tissue- and developmental contexts; enriched for signal transduction, intracellular transport, and transcriptional regulation; likely involved in coleoid-specific traits such as neural and arm crown development; exhibit functional plasticity |
| | **Context-specific chromatin loops** | Very dynamic, differential across species, tissues, and developmental stages | Common, rarely reused across developmental stages or tissues, TF binding motifs enriched at boundaries, including tissue-specific motifs, complex expression patterns; upregulation and downregulation of anchor point genes in the same loop is common, low co-expression (Pearson's $r$ = 0.044, 0.203, 0.200 for *E. scolopes* stages 20, 25, and 29, respectively) | Enable fine-scale, stage- or tissue-specific regulation; enriched for cell–cell adhesion, hormonal signalling, and neural processes; associated with organogenesis and functional maturation; exhibit high regulatory plasticity |

**Figure S31. Hierarchy of regulatory constraint across chromatin interactions in coleoid cephalopods.**
Schematic representation of regulatory constraint levels across different classes of topological interactions in the coleoids. The continuum reflects a hierarchy of stability shaped by the interplay between ancestral chromosomal state, clade specificity, and type of 3D structure. Differential constraint modulates the extent to which large-scale genomic processes, such as rearrangements and expansions, reshape regulatory architecture, ultimately contributing to the emergence of the distinct gene expression patterns and phenotypic innovations in the coleoids. Interaction classes are colour-coded as in previous figures: pink indicates interactions shared across all coleoids; green and purple indicate Decapodiformes- and *O. bimaculoides*-specific interactions, respectively; red denotes gene pairs located on the same *P. maximus* chromosome, and blue indicates those on different *P. maximus* chromosomes.
