## Supplementary notes for "Genome reorganisation and expansion shape 3D genome architecture and define a distinct regulatory landscape in coleoid cephalopods"

**Supplementary Note 1. Braiding entropy**

While fusion-with-mixing has been reported in the literature, measurements of mixing are not clearly established. As more genomes become available we can begin to measure not just positional entropy (shuffling along a genomic coordinate), but also mixing of the homology assignments ('lines') between multiple genomes or species. For this we applied concepts from topological theory to investigate the degree and entropy of braided structures (i.e., mixing of homologous coding and non-coding element order, irrespective of the exact base pair position) on chromosomes[1]. For the example shown in Fig. 2C, the homologous region between *S. officinalis* and *E. scolopes* (the Decapodiformes) showed a braiding entropy of < 0.01 (indicating almost no mixing), whereas between the more distantly related species *E. scolopes* and *O. bimaculoides* (Coleoidea), high entropy of 0.96 was detected. We quantified the degree of mixing for three different sizes of homologous regions (1 Mb, 10 Mb, and 50 Mb) and found the smallest window had very little mixing signal difference (median entropy of < 0.001 for all species combinations). The largest difference in mixing between these two phylogenetic nodes was observed for 10 Mb windows and comprised an entropy of 0.6 between *E. scolopes* and *O. bimaculoides*, compared to < 0.001 between *E. scolopes* and *S. officinalis*. The time of coleoid cephalopod separation was thus sufficient to achieve a relatively high mixed state, whereas decapodiform species showed almost no entropy increase, despite multiple apparent inversions on Oxford dotplots (Fig. 2A). For larger regions, such as 50 Mb, we observed higher entropy of mixing: 1.53 between *E. scolopes* and *O. bimaculoides* and 0.84 between *E. scolopes* and *S. officinali*s. Together, this suggests that within Decapodiformes the positioning of coding and non-coding elements is not as highly mixed or entropic, especially for smaller windows, whereas between Decapodiformes and Octopodiformes, the states are highly mixed.

To test the degree of mixing within homologous chromosome or sub-chromosomal regions with respect to their genome organisation, we partitioned the mixing results above into categories of regions that span gene pairs from different interaction categories, according to their Micro-C strength: (1) interacting across coleoids, (2) *O. bimaculoides*-only interactions (octopodiform-only), (3) decapodiform-only interactions, or (4) gene pairs not in conserved coleoid interactions. Between *E. scolopes* and *O. bimaculoides*, the median degree of braid entropy for 10 Mb homologous regions that contain at least one conserved coleoid interacting pair bin was 0.609, for regions with decapodiform-only interactions 0.529, for regions with octopodiform-only interactions 0.429, compared to 0.306 for 10 Mb regions that had no conserved interaction in coleoids (Wilcoxon test, $P = 0.054$). Similar to the results for all windows, the braiding entropy scaled with the homology window size, with smaller windows having less entropy than larger ones. For 1 Mb homologous windows, windows with interactions conserved across the coleoids showed entropy of 0.001, 0 for decapodiform-only, 0.001 for octopus-only, and 0 for windows with no conserved interactions. For 50 Mb homologous windows, the entropy was 1.283, 1.426, and 0.985 for windows with conserved coleoid interactions, decapodiform-only interactions, and octopodiform-only interactions, respectively. No 50 Mb windows had exclusively 'gene pairs not in conserved coleoid interactions'. This indicates that genomic topological linkages as measured by conserved interactions facilitate stronger mixing within the homologous regions. This complements the notion of frequent syntenic breaks at topological domain boundaries[2,3] with the indication of a stronger mixing signal within domains and suggests that a certain minimal size of a compartment is required (i.e., more than 1 Mb) for the mixing to manifest itself evolutionarily. The smaller mixing entropy for octopodiform-only interactions suggests that these regions had experienced less mixing compared to *E. scolopes*, whereas higher entropy of decapodiform-only interactions suggests that these other regions have undergone more mixing in the context of their stronger interaction environments in *E. scolopes*. This higher degree of mixing may thus reflect a more ancient origin of decapodiform interactions, allowing more time for mixing.

**Supplementary Note 2. Interaction conservation and ancestral genome organisation shape tissue-wide gene expression and reflect regulatory constraints**

To investigate whether chromatin interactions influence gene expression, we analysed TPM-normalised expression levels of genes across *E. scolopes* tissues and interaction categories (Fig. S3, Table S3). For this analysis, all genes in the 'interacting across the coleoids', 'interacting Decapodiformes only', 'interacting *O. bimaculoides* only' were retained, while genes in the 'not in conserved coleoid interactions' category were included only if they were exclusive to this category and not present in any interacting categories, preventing overlaps and ensuring clear comparisons.

Within each interaction category, variation in gene expression in *E. scolopes* was substantial, with broad interquartile ranges indicating tissue-specific differences in expression levels. However, across interaction categories, gene expression distributions were largely similar across tissues (Fig. S3B, C). Genes interacting in all species showed the highest median expression across *E. scolopes* tissues, followed by genes interacting in *E. scolopes* only, then genes interacting in *O. bimaculoides* only, with genes not in conserved coleoid interactions exhibiting the lowest median expression. These patterns suggest that genes in conserved interactions may be located in transcriptionally active regions that promote consistent and higher expression levels across tissues. Notably, genes interacting in all species had significantly higher expression (Wilcoxon test, BH-adjusted) in several tissues with coleoid innovations (Table S3) such as the subesophageal lobe, supraesophageal lobe, central brain, light organ, optic lobes, hectocotylus and accessory nidamental gland (ANG). Similarly, genes interacting in Decapodiformes only also had significantly higher expression compared to genes not in conserved coleoid interactions in some of these coleoid-specific tissues, including the subesophageal lobe, supraesophageal lobe, central brain, central core, optic lobes, hectocotylus, and ANG. This suggests that conserved chromatin interactions may contribute to the evolution of complex regulatory networks underpinning coleoid-specific adaptations. In contrast, genes interacting only in *O. bimaculoides* exhibited significantly higher expression than genes not in conserved coleoid interactions in fewer tissues, including the central core, left optic lobe, right optic lobe, and ANG. However, these effects were weaker and less consistent across tissues compared to genes interacting in all species, reinforcing the idea that conserved interactions play a role in regulatory evolution of coleoid novel traits.

To further assess the tissue specificity of *E. scolopes* gene expression across interaction categories, we calculated the tissue specificity index ($\tau$). Genes in the ‘interacting in all species’ category had the lowest $\tau$ values (median $\tau = 0.551$), followed by those ‘interacting in Decapodiformes only’ ($\tau = 0.580$), indicating broader expression across tissues (Table S4Ai). In contrast, genes interacting only in *O. bimaculoides* ($\tau = 0.606$) and genes not in conserved coleoid interactions ($\tau = 0.665$) exhibited significantly higher tissue specificity (Wilcoxon test, BH-adjusted) (Table S4Aii). Statistical comparisons confirmed that genes interacting in all species had significantly lower $\tau$ values than those in the ‘interacting in *O. bimaculoides* only’ ($P = 0.014$) and ‘genes not in conserved coleoid interactions’ (P =

0.0004) categories. However, differences between ‘interacting in Decapodiformes only’ and ‘interacting in all species’ were not significant, suggesting that conservation of interactions within Decapodiformes may also be linked to broader expression patterns.

These results are consistent with recent large-scale comparative transcriptomic studies showing that genes with broad expression patterns across tissues are under stronger evolutionary constraint, while more tissue-specific genes are less conserved at both the sequence and regulatory levels[4,5]. Broadly expressed genes, a proxy for pleiotropy, are less likely to undergo rapid changes in expression or sequence, likely due to their roles in multiple tissues and biological processes[4,5] Furthermore, highly expressed genes are likewise associated with stronger purifying selection, leading to slower evolutionary rates, likely due to the higher cost of transcriptional and translational errors[6,7]. Collectively, these findings support the idea that the most conserved interacting gene pairs are subject to particularly strong selective constraints, likely due to their broad expression and pleiotropic function.

To assess whether ancestral genomic organisation influences gene expression in *E. scolopes*, we compared co-expression patterns between genes whose orthologs were located on the same or different *P. maximus* chromosomes. Gene pairs interacting across the coleoids and located on the same *P. maximus* chromosome showed significantly higher *E. scolopes* gene expression correlation than those on different chromosomes (Wilcoxon test, BH-adjusted P = 0.0094), suggesting that shared chromosomal context may contribute to regulatory coordination and constraint. A similar trend was observed for gene pairs interacting only in Decapodiformes, though this difference was marginally non-significant after correction (BH-adjusted P = 0.0574). In contrast, there was no significant difference in co-expression for gene pairs not in conserved coleoid interactions or those interacting only in *O. bimaculoides* (i.e not interacting in the species with gene expression data, *E. scolopes*).

Furthermore, when we compared expression levels across tissues (Fig. S3D-F), we found that for genes in gene pairs interacting in all species, those located on the same *P. maximus* chromosome exhibited higher median expression than those on different chromosomes (Fig. S3D), with significant differences in a range of tissues, some of which could be classed as coleoid novelties, such as the ANG, B1 arm, B4 arm, central brain, light organ, hectocotylus, optic lobes, right tentacle, right white body and skin (Table S6). There were no significant differences in expression level between genes not in conserved coleoid interactions with different chromosomal origins across tissues. This suggests that interacting gene pairs that have remained syntenic since the ancestral molluscan genome may promote higher expression and be under stronger regulatory constraints due to their transcriptional importance than genes from different *P. maximus* chromosomes, whereas genes not in conserved coleoid interactions are under similar relaxed constraints, regardless of chromosomal origin. On the other hand, tissue specificity (τ) did not differ significantly between same and different chromosomal origin across any of the four interaction categories, suggesting that ancestral chromosomal context is not a major determinant of expression breadth. Together, these findings indicate that while ancestral synteny correlates with higher expression and co-expression in conserved interacting pairs, interaction status more strongly reflects underlying pleiotropic constraints and evolutionary pressures.

Hierarchical clustering of gene expression patterns were very similar (Fig. S3C, F), with no strong separation based on interaction status or chromosomal origin. While some variation in expression levels is evident from the heatmaps, genes across all groups exhibited broadly comparable expression distributions. This suggests that while conserved interactions may contribute to overall expression levels (as indicated by the boxplots), they do not necessarily drive distinct expression patterns across tissues.

No significant differences in intervening gene coverage were observed between gene pairs located on the same versus different *P. maximus* chromosomes for any of the interacting categories in either species. In contrast, gene pairs not in conserved interactions in coleoids showed significant differences in both *E. scolopes* (Wilcoxon test, BH-adjusted $P = 1.95 \times 10^{-5}$) and *O. bimaculoides* (BH-adjusted $P = 7.70 \times 10^{-4}$), potentially reflecting that relaxed evolutionary constraints (given the absence of a conserved strong interaction signal) in ancestral pairs that have had more time to accumulate intervening genes.

Overall, these findings suggest that conserved chromatin interactions are associated with higher and more broadly expressed gene expression patterns across tissues compared to genes that do not interact with other genes. This indicates that interacting genes may be highly conserved housekeeping genes, maintaining essential functions across multiple tissues[8]. In contrast, genes not in conserved coleoid interactions may have more specialised roles and/or evolve under relaxed regulatory constraints. Furthermore, chromosomal origin within *P. maximus* significantly impacts gene expression levels and co-expression but does not influence tissue specificity or intervening gene coverage for interacting gene pairs, reinforcing the distinct role of ancestral genome organisation in shaping regulatory interactions. These results provide further evidence that large-scale genomic processes play a critical role in the evolution of complex gene regulatory patterns in the coleoids.

**Supplementary Note 3. Functional enrichment patterns reflect interaction conservation and ancestral chromosomal context of gene pairs**

To investigate potential functional differences between gene interaction categories, we performed GO enrichment analyses across biological processes, molecular functions, and cellular components for genes in each interaction group (Fig. S4). As with the expression analyses, all genes in the 'interacting across the coleoids', 'interacting Decapodiformes only', 'interacting *O. bimaculoides* only' were retained, while genes in the 'not in conserved coleoid interactions' category were included only if they were exclusive to this category. Genes interacting in all species were strongly enriched for fundamental biological processes such as ribosome biogenesis, RNA processing, peptide biosynthesis, and translation, as well as structural components like the ribosome and ribonucleoprotein complex, suggesting a core regulatory role for these genes across coleoids, consistent with the results of the gene expression analyses.

Genes interacting in Decapodiformes only were primarily enriched for protein catabolic processes, particularly those dependent on ubiquitin and proteasome function, including enrichment in components of the proteasome complex and catalytic complexes. This points to a potential role in lineage-specific regulation of protein degradation and turnover. In contrast, genes interacting only in *O. bimaculoides* showed enrichment for both catabolic and biosynthetic processes, including protein folding, translation, and peptide metabolic processes, as well as peptidase activity and RNA polymerase complexes. This may reflect species-specific adaptation or regulatory rewiring.

Genes that were not in conserved coleoid interactions were notably enriched for neural signalling-related functions, including neurotransmitter receptor activity, ligand-gated ion channel activity, and components of the synaptic membrane and postsynaptic compartments. These genes were also associated with DNA-binding transcription factor (TF) activity, suggesting a regulatory role possibly decoupled from deeply conserved chromatin architecture.

When comparing genes interacting in all species based on their ancestral chromosomal origin, we observed notable differences in enriched GO terms (Fig. S8). Genes interacting across the coleoids whose orthologs were located on the same *P. maximus* chromosome were strongly enriched for housekeeping and biosynthetic functions, including translation, peptide biosynthesis, and ribosomal structure. In contrast, genes interacting across the coleoids but located on different *P. maximus* chromosomes showed enrichment for processes such as chromatin organisation, DNA replication, and histone modification. These results indicate that even within highly conserved interaction groups, ancestral chromosomal origin can influence gene function, with genes in ancestral interactions more associated with core cellular processes and those emerging post genome rearrangement potentially contributing to regulatory adaptation in coleoids. For genes not in conserved coleoid interactions, GO results were more similar when comparing those with same vs different *P. maximus* chromosome origin (Fig. S8). Taken together, these results suggest that ancestral chromosomal proximity plays a role in maintaining the regulatory and functional coherence of conserved interactions between gene pairs, whereas genes not in conserved coleoid interactions may evolve under fewer structural constraints and show more functional heterogeneity.

To complement the GO analysis, we also assessed the enrichment of specific protein domain annotations associated with cephalopod nervous system development[9] across gene pair categories. No significant enrichment of zinc finger or cadherin genes was detected in any gene pair group (Fig. S13), suggesting that these domain families are not strongly associated with conserved chromatin interaction status. Instead, their distribution may reflect lineage-specific expansions more closely tied to species-specific or novel aspects of genome architecture, rather than ancient, conserved regulatory interactions and core biological functions.

**Supplementary Note 4. Loop-associated genes are enriched for regulatory and signalling functions and transcription binding factor motifs across species and contexts**

To assess whether genes located at chromatin loop anchors are associated with particular functional roles, we performed GO enrichment analyses across all species, developmental stages and tissues (Fig. S12, S22, S23, S24). Despite differences in loop distribution and usage across categories, a consistent pattern emerged: loop-associated genes are enriched for regulatory, signalling, and cellular coordination processes, consistent with the proposed role of chromatin loops in enabling dynamic, context-dependent gene expression.

In *E. scolopes* whole-embryo data, loop-associated genes were enriched for processes related to cell adhesion and hormone-mediated signalling, including responses to steroids and lipids (Fig. S12A). Analysis of earlier embryonic stages further supported this, with stage 25 loop genes showing strong enrichment for cell–cell adhesion, as well as calcium ion binding (Fig. S22), consistent with the onset of tissue maturation and increased intercellular coordination during late organogenesis[10]. In *S. officinalis*, whole-embryo loop genes showed enrichment for processes such as protein transport, regulation of signal transduction and cell communication, as well as components of nuclear protein-containing complexes and ubiquitin transferase activity (Fig. S12B), suggesting loop-mediated control over intracellular trafficking and post-translational regulation. In *O. bimaculoides*, loop genes in whole embryos were strongly associated with cytoskeletal organisation, GTPase signalling, and calmodulin-dependent kinase activity, with enrichment of dynein, microtubule-associated, and actin-binding proteins (Fig. S12C), suggesting that chromatin loops may support structural remodeling and rapid signalling in complex, highly active tissues such as the coleoid nervous system.

Tissue-level GO enrichment analyses in *S. officinalis* (Fig. S23) and *O. bimaculoides* (Fig. S24) reinforced whole-embryo patterns and revealed additional tissue-specific specialisations. In *S. officinalis*, central brain loops were consistent with the regulatory and signalling enrichments seen at the whole-embryo level, while the eyes and internal organs showed more restricted enrichment, including cellular organisation and replication-related terms. In *O. bimaculoides*, loop genes in neural tissues (central brain, eyes and optic lobes, and arm crown) were consistently enriched for functions similar to those associated with whole-embryo samples, supporting a role for loops in neural structure and fast signal transduction. On the other hand, internal organ loops were instead enriched for DNA and RNA polymerase activity, ion transport, and protein-containing complexes, reflecting core cellular functions. These findings suggest that chromatin loops in neural tissues are functionally geared toward plasticity and dynamic regulation, whereas those in internal organs are associated with essential cellular maintenance.

Taken together, these findings suggest that chromatin loops frequently bring together genes involved in plastic, condition-sensitive regulatory functions, such as signal transduction, intracellular transport, and tissue-specific organisation. These functional profiles stand in contrast to those of genes in gene pairs grouped by interaction category. While genes located in loops, which are rarely conserved, were enriched for dynamic regulatory processes, genes involved in conserved interactions across all species were primarily associated with core biosynthetic and housekeeping functions. This dichotomy supports a model in which conserved gene interactions maintain essential, stable cellular processes under strong regulatory constraint, whereas chromatin loops act as flexible, evolutionarily adaptable regulatory elements that facilitate precise control of gene expression in response to developmental, spatial, or environmental cues.

Furthermore, we found that across all species, developmental stages, and tissues, zinc finger-containing genes comprised ~7–14% of loop-associated gene sets, while cadherin genes generally remained below 2%, with notable exceptions (Fig. S13). In *E. scolopes*, zinc finger gene representation peaked in stage 20 loops, whereas cadherin gene content increased steadily through development, reaching ~8.5% at stage 29. In *S. officinalis*, zinc finger gene proportions were lower (3–5%), but consistently higher than cadherins. In *O. bimaculoides*, the proportion of loop-associated zinc finger genes were notably high (~14%) across all tissues, consistent with a known lineage-specific expansion of this gene family in Octopodiformes[11], and not necessarily driven by chromatin interaction patterns.

No significant enrichment of zinc finger genes was detected across loop datasets in any species compared to the rest of the genome (Fisher's exact test with BH correction). However, despite the overall low abundance of cadherin genes, enrichment analysis revealed significant overrepresentation in several categories. Cadherin genes were significantly enriched in the *E. scolopes* stage 25 and stage 29 loop sets (Fisher's exact test, BH-adjusted P = 2.22e-04 and P = 1.22e-06, respectively), and also in the *S. officinalis* arm crown loop set (BH-adjusted P = 0.0273), indicating stage- or tissue-specific functional relevance. These findings are consistent with stage 25 and 29 representing key periods of nervous system development in *E. scolopes*[10]. The *O. bimaculoides* arm crown loops also showed the highest proportion of cadherin genes across tissues, consistent with the pattern found in *S. officinalis*, although the enrichment was not statistically significant. The high proportion of cadherin genes in the *S. officinalis* and *O. bimaculoides* arm crown loops likely reflects the neural character of the arm tissues in both species, which support complex sensorimotor functions. Given the conserved role of cadherins in nervous system development across metazoans[12] and the expansion of protocadherins in the Octopodiformes[11] and Decapodiformes[13,14], their enrichment in loop-associated regions of neural tissues likely reflects their involvement in establishing and maintaining complex neural architecture in the coleoids.

Developmental stage and tissue-specific motif enrichment at loop anchors mostly mirrored patterns from whole-embryo analyses (main text), including consistent significant (hypergeometric test, BH-adjusted $P < 0.05$) detection of low-complexity and repeat-associated sequences, and widespread enrichment of Forkhead, bHLH, MYB, Homeobox, and CTCF-like motifs across species. Building on this shared regulatory foundation, tissue-specific analyses revealed additional transcription factors reflecting context-dependent regulatory activity. In *O. bimaculoides*, the eyes and optic lobes were enriched for neural regulators such as Barhl2, Vax2, and NeuroG2, while central brain, internal organs, and the arm crown displayed more general motifs including Tcf3 and GA-rich sequences. In *S. officinalis*, all tissues showed additional enrichment for developmental regulators such as Vax2 and Dlx1. These findings suggest that while some regulatory motifs shape a conserved regulatory backbone, dynamic recruitment of context-specific transcription factors likely contributes to the spatial and temporal control of gene expression.

**Supplementary Note 5. Regulatory features of loops and comparison to compartment structures**

We quantified CNE enrichment in loop anchors and interloop regions across *E. scolopes*, *O. bimaculoides*, and *S. officinalis*. Loop anchors showed significant enrichment for conserved CNEs in all three species compared to background genomic regions (Wilcoxon test, BH-adjusted $P = 6.6 \times 10^{-4}$ for *E. scolopes* CNEs conserved in *O. bimaculoides*, $4.0 \times 10^{-10}$ for *O. bimaculoides* CNEs conserved in *E. scolopes*, and $1.2 \times 10^{-13}$ for *S. officinalis* CNEs conserved in *E. scolopes*). Interloop regions also exhibited significant CNE enrichment compared to the genomic background, although to a lesser extent in *E. scolopes* and *O. bimaculoides* (BH-adjusted $P = 5.4 \times 10^{-3}$ and $2.7 \times 10^{-2}$, respectively), and very strongly in *S. officinalis* (BH-adjusted $P = 1.6 \times 10^{-61}$), potentially reflecting the shorter evolutionary distance between decapodiform–decapodiform CNEs relative to octopodiform–decapodiform CNEs. Direct comparisons of loop anchors and interloop regions showed that CNE density was significantly higher in loop anchors in *E. scolopes* and *O. bimaculoides* (BH-adjusted $P = 1.6 \times 10^{-2}$ and $4.4 \times 10^{-8}$, respectively), and marginally higher in *S. officinalis* (BH-adjusted $P = 5.4 \times 10^{-2}$). Together, our results suggest that chromatin loop topology, especially the anchors, which form the spatial proximity-based interactions between distal genomic regions, plays a regulatory role in coleoid genomes.

To further assess how chromatin loops relate to gene content, we compared the proportion of loop regions occupied by genes to genome-wide coverage across all gene pairs in all four interaction categories as a reference set. In both *E. scolopes* and *O. bimaculoides* whole-embryo samples, loop regions exhibited lower median gene coverage than genome-wide gene pairs. *E. scolopes* loops showed 19.1% median gene coverage coverage, while *O. bimaculoides* loops had median gene coverage of 23.3%. These values contrast with genome-wide gene pairs, where median gene coverage ranged from 33.0% to 46.2% (Table S5). These differences were statistically significant between loops and gene pairs (Wilcoxon test, BH-adjusted $p < 0.0001$), confirming that loops are consistently more gene- sparse. The higher gene content in *O. bimaculoides* is consistent with its more compact genome, which has a smaller total size despite having a similar number of annotated genes to *E. scolopes*[11,13].

Next, we quantified normalised ATAC peak enrichment in loop anchors, interloop regions, and the genomic background at stages 20, 25, and 29. At all three developmental stages, loop anchor regions showed significantly greater chromatin accessibility than background regions (Wilcoxon test, BH-adjusted $P = 0.0092$ at stage 20, 0.0092 at stage 25, and $9.0 \times 10^{-8}$ at stage 29). At stage 29, interloop regions were also significantly enriched for ATAC peaks ($P = 0.010$) compared to background regions, though this enrichment was not observed at stages 20 or 25. Pairwise comparisons further revealed that loop anchors had significantly higher accessibility than interloop regions at all stages (BH-adjusted $P = 0.035$ at stage 20, 0.022 at stage 25, and 0.0063 at stage 29).

Median normalised fractional chromatin accessibility at loop anchors increased across development, from 0.559 at stage 20 to 0.630 at stage 25 and 0.634 at stage 29. Background regions, including interloop space, showed a similar upward trend, indicating a general increase in chromatin accessibility during embryogenesis. Despite this global shift, loop anchors exhibited greater variability, suggesting that only a subset of anchors become highly accessible during development.

To test whether loops indeed represent more dynamic regulatory structures than higher-order chromatin features, we performed compartment calling and differential compartment analysis across developmental stages and tissues. In contrast to loops, compartments appeared far more conserved; approximately 65–75% of compartments were shared across tissue comparisons (Fig. S25A). Most differential events were classified as complex rearrangements, where existing compartments were neither split nor merged but reorganised into entirely new configurations[15], potentially enabling localised expression shifts while preserving broader architectural domains. Nevertheless, the general patterns of tissue-specific differences observed in loops and compartments were broadly consistent, suggesting that both levels of genome architecture contribute to tissue-specific regulation, albeit with differing degrees of conservation. UpSet plots indicated that the same set of compartments were recurrently identified as differential across tissues, suggesting a shared group of domains with heightened regulatory plasticity. However, this pattern may partly reflect our analytical approach, which classifies compartments absent in a tissue as differential (Fig. S25B). Taken together, these results indicate that chromatin loops represent plastic, context-dependent regulatory features of genome organisation, potentially facilitating fine-scale control of gene expression. Furthermore, they may evade the evolutionary constraints experienced by whole compartments, which serve as more stable, higher-order structures.

**Supplementary Note 6. Example of dynamic chromatin loop across development and associated gene expression**

To illustrate a developmentally dynamic chromatin loop in *E. scolopes*, we highlight an interaction on chromosome 5 that is present at stage 20 but absent by stage 25 and 29 (Fig. 4D). At stage 20, the loop coincides with elevated chromatin accessibility (ATAC-seq peaks), local insulation at loop borders (Fig. 4E) and peak gene expression (Fig. 4F), suggesting it may coordinate the activation of this regulatory module during the onset of tissue differentiation. The loop brings together six genes; Prefoldin, two subunits of Chromatin Assembly Factor 1 (CAF-1), the snRNA-activating complex (SNAPc), Sas10/Utp3/C1D, and Sirtuin, all *o*f which are functionally linked to chromatin assembly, cytoskeletal organisation, and RNA processing, processes essential for early embryogenesis and organogenesis. Each of these genes has well-established developmental roles: Prefoldin facilitates the folding of cytoskeletal proteins, such as actin and tubulin, which are essential for morphogenesis and cell division[16]; CAF-1 subunits organise chromatin and deposit histones to preserve epigenetic states during rapid cell proliferation[17]; SNAPc regulates snRNA transcription, necessary for pre-mRNA splicing during zygotic genome activation[18]. Sas10/Utp3/C1D belong to the Sas10/C1D protein family, which is involved in small subunit rRNA processing and ribosome biogenesis[19,20]; and Sirtuin modulates chromatin accessibility and epigenetic silencing during asymmetric divisions, which are crucial for patterning and lineage specification[21,22]. Together, their activity likely orchestrates key transitions during early organogenesis. By stage 25, the disappearance of this loop is accompanied by a reduction in both insulation and expression, consistent with a shift from specification to more stable lineage commitment. Notably, while SNAPc (cluster_11478) and one CAF-1 subunit (cluster_18817) are syntenic in the ancestral *P. maximus* genome, the remaining genes are located far apart on the same or across different chromosomes. This suggests that the looped configuration is a derived feature, likely facilitated by genome rearrangements in the coleoid ancestor. This is consistent with the idea that large-scale genomic changes in the coleoid lineage have enabled the emergence of complex traits such as the unique coleoid developmental program through novel 3D regulatory architectures.

**Supplementary Note 7. Transcription factor binding motif enrichment analysis in the CRISPR-targeted knockout region**

To investigate the potential regulatory role of the ~1 kb intronic sequence deleted by CRISPR-Cas9 (Fig. 5B), we performed motif enrichment analysis using HOMER on the knockout region (Table S11). After correction, two motifs were found to be significantly enriched: a Hox-like motif (Hoxb4) and ATHB25 (ZFHD) (BH-corrected $P < 0.05$). Because ATHB25 is plant-derived, it was excluded from biological interpretation. In total, eight plant/non-metazoan motifs were filtered. Thus, Hoxb4 is the sole significant metazoan signal, consistent with input from homeodomain factors involved in body patterning and neural development[23].

Because the target set comprised tiled 200 bp windows across the ~ 1 kb locus, per-motif counts were low and statistical power was limited; accordingly, we did not expect many post-correction discoveries compared to genome-wide analyses. Therefore, although not significant after correction, the presence of additional metazoan motifs in the knockout locus is still informative about potential regulatory inputs. We detected sites corresponding to Lhx6 (LIM-homeobox), IRF3 and a composite bZIP:IRF motif, as well as general regulatory factors (NF1/CTF, Ap4/Tfap4, Hnf1); selected bHLH signals (e.g., Ap4, Ptf1a) were also observed at nominal levels. Functionally, these factors are consistent with roles in neural or developmental regulation: Lhx6 is implicated in neuronal differentiation[24]; Ap4/Tfap4 and other bHLH proteins modulate proliferation–differentiation programs[30]; and IRF-family and bZIP:IRF composite sites can interface immune and activity-dependent signalling with neural gene regulation[25,26].

MYNN (Myoneurin) motifs were also detected within the deleted sequence and just upstream of it. Notably, the upstream site lies in an intronic region conserved in *O. bimaculoides* (Fig. 5B) within a repetitive interval not targetable by guide RNAs. The presence of these MYNN motifs, together with their evolutionary conservation, suggests that this region may form part of a broader enhancer module, with the deleted sequence representing one functional component within it. MYNN itself is not well characterised in neural contexts, but it encodes a BTB/POZ and C2H2 zinc finger transcription factor[27], a family broadly involved in gene regulation. Notably, related family members such as MYT1 and MYT1L, sometimes collectively referred to as neural zinc finger (NZF) proteins, play well-established roles in neuronal differentiation and neural circuit development[28,29]. In cephalopods, zinc finger transcription factors have also been implicated in neural complexity[11,13], further supporting their functional relevance in this clade.

Taken together, these enriched motifs support the hypothesis that the targeted intronic sequence functions as a putative neural regulatory element such as an enhancer. Although the precise TFs operating in cephalopods remain to be defined, the conserved presence of neural- and development-associated motifs implies shared regulatory logic across bilaterians. Given the phenotypic consequences of the knockout, such as reduced brain size, malformed optic lobes, and delayed development, we infer that the deleted region normally integrates transcription factor inputs critical for neural gene regulation. These results suggest the intronic sequence acts as a regulatory hub, contributing to the expression of Quaking B and potentially additional genes within the conserved chromatin loop.

### References

1. Thiffeault, J.-L. & Budisic, M. Braidlab: A software package for braids and loops. *arXiv [math.GT]* (2014).
2. Farré, M., Robinson, T. J. & Ruiz-Herrera, A. An Integrative Breakage Model of genome architecture, reshuffling and evolution: The Integrative Breakage Model of genome evolution, a novel multidisciplinary hypothesis for the study of genome plasticity. *Bioessays* **37**, 479–488 (2015).
3. Liao, Y., Zhang, X., Chakraborty, M. & Emerson, J. J. Topologically associating domains and their role in the evolution of genome structure and function in Drosophila. *Genome Res.* **31**, 397–410 (2021).
4. Mantica, F. *et al.* Evolution of tissue-specific expression of ancestral genes across vertebrates and insects. *Nat. Ecol. Evol.* **8**, 1140–1153 (2024).
5. Jiang, W. & Chen, L. Tissue specificity of gene expression evolves across mammal species. *J. Comput. Biol.* **29**, 880–891 (2022).
6. Drummond, D. A., Bloom, J. D., Adami, C., Wilke, C. O. & Arnold, F. H. Why highly expressed proteins evolve slowly. *Proc. Natl. Acad. Sci. U. S. A.* **102**, 14338–14343 (2005).
7. Pál, C., Papp, B. & Hurst, L. D. Highly expressed genes in yeast evolve slowly. *Genetics* **158**, 927–931 (2001).
8. Eisenberg, E. & Levanon, E. Y. Human housekeeping genes, revisited. *Trends Genet.* **29**, 569–574 (2013).
9. Albertin, C. B. & Katz, P. S. Evolution of cephalopod nervous systems. *Curr. Biol.* **33**, R1087–R1091 (2023).
10. Lee, P. N., Callaerts, P. & de Couet, H. G. The embryonic development of the Hawaiian bobtail squid (Euprymna scolopes). *Cold Spring Harb. Protoc.* **2009**, db.ip77 (2009).
11. Albertin, C. B. *et al.* The octopus genome and the evolution of cephalopod neural and morphological novelties. *Nature* **524**, 220–224 (2015).
12. Hulpiau, P. & van Roy, F. Molecular evolution of the cadherin superfamily. *Int. J. Biochem. Cell Biol.* **41**, 349–369 (2009).
13. Belcaid, M. *et al.* Symbiotic organs shaped by distinct modes of genome evolution in cephalopods. *Proc. Natl. Acad. Sci. U. S. A.* **116**, 3030–3035 (2019).
14. Rencken, S. *et al.* Chromosome-scale genome assembly of the European common cuttlefish Sepia officinalis. *Genomics* (2025).
15. Cresswell, K. G. & Dozmorov, M. G. TADCompare: An R package for differential and temporal analysis of topologically associated domains. *Front. Genet.* **11**, 158 (2020).

16. Liang, J. *et al.* The functions and mechanisms of prefoldin complex and prefoldin-subunits. *Cell Biosci.* **10**, 87 (2020).
17. Liu, C.-P. *et al.* Structural insights into histone binding and nucleosome assembly by chromatin assembly factor-1. *Science* **381**, eadd8673 (2023).
18. Sun, J. *et al.* Structural basis of human SNAPc recognizing proximal sequence element of snRNA promoter. *Nat. Commun.* **13**, 6871 (2022).
19. Mitchell, P. Rrp47 and the function of the Sas10/C1D domain. *Biochem. Soc. Trans.* **38**, 1088–1092 (2010).
20. Chen, Y.-J. C., Wang, H.-J. & Jauh, G.-Y. Dual role of a SAS10/C1D family protein in ribosomal RNA gene expression and processing is essential for reproduction in Arabidopsis thaliana. *PLoS Genet.* **12**, e1006408 (2016).
21. Jing, H. & Lin, H. Sirtuins in epigenetic regulation. *Chem. Rev.* **115**, 2350–2375 (2015).
22. Fang, Y., Tang, S. & Li, X. Sirtuins in metabolic and epigenetic regulation of stem cells. *Trends Endocrinol. Metab.* **30**, 177–188 (2019).
23. Morgan, R., Pettengell, R. & Sohal, J. The double life of HOXB4. *FEBS Lett.* **578**, 1–4 (2004).
24. Kim, D. W. *et al.* Gene regulatory networks controlling differentiation, survival, and diversification of hypothalamic Lhx6-expressing GABAergic neurons. *Commun. Biol.* **4**, 95 (2021).
25. Joshi, R. *et al.* IRF3 regulates neuroinflammatory responses and the expression of genes associated with Alzheimer's disease. *J. Neuroinflammation* **21**, 212 (2024).
26. Rodríguez-Martínez, J. A., Reinke, A. W., Bhimsaria, D., Keating, A. E. & Ansari, A. Z. Combinatorial bZIP dimers display complex DNA-binding specificity landscapes. *Elife* **6**, (2017).
27. GeneCards Human Gene Database. MYNN Gene - GeneCards. https://www.genecards.org/cgi-bin/carddisp.pl?gene=MYNN.
28. Hudson, L. D., Romm, E., Berndt, J. A. & Nielsen, J. A. A tool for examining the role of the zinc finger myelin transcription factor 1 (Myt1) in neural development: Myt1 knock-in mice. *Transgenic Res.* **20**, 951–961 (2011).
29. Chen, J., Yen, A., Florian, C. P. & Dougherty, J. D. MYT1L in the making: emerging insights on functions of a neurodevelopmental disorder gene. *Transl. Psychiatry* **12**, 292 (2022).
30. Wong, M. M.-K., Joyson, S. M., Hermeking, H. & Chiu, S. K. Transcription factor AP4 mediates cell fate decisions: To divide, age, or die. *Cancers (Basel)* **13**, 676 (2021).
