## Supplementary table figure legends for "Genome reorganisation and expansion shape 3D genome architecture and define a distinct regulatory landscape in coleoid cephalopods"

**Table S1. Summary of all Micro-C sequencing libraries used in this study.**
The “Sample ID” refers to the unique identifier for each library. “Tissue” indicates the dissected material used for Micro-C; “whole embryo” refers to undissected samples. The developmental stage corresponds to the embryonic stage at the time of collection. The “Number of pooled embryos” specifies how many embryos were combined per library; libraries based on single embryos are indicated with a value of 1. The “Source” column lists the institution or laboratory that provided the material. “Readmode” indicates the sequencing platform and chemistry used. The “Number of read pairs” represents the total raw read pairs generated per sample. Where applicable, the “Notes” column indicates whether read pairs were downsampled prior to downstream analysis.

**Table S2. Sequencing depth and mapping summary for Micro-C libraries used in this study.**
Summary of sequencing depth and mapping efficiency for all Micro-C libraries used in this study. Columns indicate total number of read pairs generated, followed by the number of mapped reads, number of mapped reads, and the final proportion of usable reads retained after mapping. Asterisks (*) denote samples that were downsampled prior to mapping, with the values shown reflecting the post-downsampling read count. Notes specify any exceptions in sample usage across specific analyses. Sample 409493 was used for all *E. scolopes* stage 29 analyses unless otherwise indicated.

**Table S3. Differential tissue-specific expression in *E. scolopes* across gene pair categories.**
Results of statistical tests comparing *E. scolopes* gene expression across different gene interaction categories in various tissues. Columns indicate the interaction status of each gene pair, the *E. scolopes* tissue examined, raw P values from Wilcoxon rank-sum tests, and adjusted P values after BH correction.

**Table S4. Tissue specificity of gene expression (τ) across interaction categories and chromosomal origin.**
Summary of tissue specificity values (τ) for *E. scolopes* genes across chromatin interaction categories and *P. maximus* chromosomal origins.
(A) Mean and median τ values grouped by interaction status.
(B) Results of Wilcoxon rank-sum tests comparing τ values between groups. Adjusted P values were calculated using BH correction.

**Table S5. Intervening gene coverage across gene pairs in *E. scolopes* and *O. bimaculoides*, grouped by interaction status and lineage conservation.**
As with the corresponding loop analysis, this analysis focused only on gene pairs containing at least one intervening gene.
(A) Median percentage of genomic space between interacting gene pairs covered by annotated genes for coleoid species with available gene annotations, *E. scolopes* and *O. bimaculoides.* Values are reported for the four interaction categories of gene pairs.
(B) Wilcoxon rank-sum test results comparing intervening gene coverage between interaction categories within each species. Reported are unadjusted and BH-adjusted P values.

**Table S6. Differential tissue-specific expression in *E. scolopes* across gene pair categories and *P. maximus* chromosomal origin.**
Results of statistical tests comparing *E. scolopes* gene expression across different gene interaction categories and *P. maximus* chromosome statuses in various tissues. Columns indicate the interaction status of each gene pair, the *E. scolopes* tissue examined, raw P values from Wilcoxon tests, and adjusted P values after BH correction.

**Table S7. Median genomic distances between gene pairs across interaction statuses, by *P. maximus* chromosomal origin categories and *P. maximus* distance bins for the three focal coleoid species.**
Median genomic distances (in base pairs) are shown for the four interaction categories of gene pairs in *E. scolopes*, *S. officinalis*, and *O. bimaculoides*. Gene pairs are grouped into either (A) being located on the same or on two different *P. maximus* chromosomes, with the latter being a proxy for coleoid gene pairs whose interactions may have emerged due to interchromosomal genome rearrangements; and (B) three distance categories based on their genomic separation in *P. maximus*: ≤ 5 Mb, > 5 Mb and ≤ 15 Mb, and > 15 Mb. Longer distances in *P. maximus* serve as a proxy for gene pairs in coleoids that are more likely to have become colocalised through intrachromosomal genome rearrangements.

**Table S8. Enriched known DNA motifs in loop anchor-associated sequences in *E. scolopes*.**
This table presents TF binding motifs significantly enriched in genomic regions at loop anchors in whole embryo *E. scolopes s*amples. Motif enrichment analysis was performed using the HOMER motif discovery suite, comparing motif occurrences in loop anchor-associated sequences (target set) to the *E. scolopes* reference genome as the background. The table includes the motif name (indicating the transcription factor or protein family), consensus binding sequence, unadjusted P value and log-transformed P value for enrichment, and the BH-adjusted P value. Also shown are the number and percentage of target sequences containing each motif, as well as the number and percentage of background genomic sequences with the motif. Only motifs with a significant enrichment ((hypergeometric test, BH-adjusted P < 0.05) are included.

**Table S9. Enriched known DNA motifs in loop anchor-associated sequences in *S. officinalis*.**
This table presents TF binding motifs significantly enriched in genomic regions at loop anchors in whole embryo *S. officinalis* samples. Motif enrichment analysis was performed using the HOMER motif discovery suite, comparing motif occurrences in loop anchor-associated sequences (target set) to the *S. officinalis* reference genome as the background. The table includes the motif name (indicating the transcription factor or protein family), consensus binding sequence, unadjusted P value and log-transformed P value for enrichment, and the BH-adjusted P value. Also shown are the number and percentage of target sequences containing each motif, as well as the number and percentage of background genomic sequences with the motif. Only motifs with a significant enrichment (hypergeometric test, BH-adjusted P < 0.05) are included.

**Table S10. Enriched known DNA motifs in loop anchor-associated sequences in *O. bimaculoides*.**
This table presents transcription factor binding motifs significantly enriched in genomic regions at loop anchors in whole embryo *O. bimaculoides* samples. Motif enrichment analysis was performed using the HOMER motif discovery suite, comparing motif occurrences in loop anchor-associated sequences (target set) to the *O. bimaculoides* reference genome as the background. The table includes the motif name (indicating the transcription factor or protein family), consensus binding sequence, unadjusted P value and log-transformed P value for enrichment, and the BH-adjusted P value. Also shown are the number and percentage of target sequences containing each motif, as well as the number and percentage of background genomic sequences with the motif. Only motifs with a significant enrichment (hypergeometric test, BH-adjusted P < 0.05) are included.

**Table S11. Enriched known DNA motifs in CRISPR-cas9 knockout region in *E. berryi*.**
This table presents transcription factor binding motifs enriched in the CRISPR knockout region (Fig. 5B) in *E. berryi*. Motif enrichment analysis was performed using the HOMER motif discovery suite, comparing motif occurrences in the knockout region to the *E. berryi* reference genome as the background. The table includes the motif name (indicating the transcription factor or protein family), consensus binding sequence, unadjusted P value and log-transformed P value for enrichment, and

the BH-adjusted P value. Also shown are the number and percentage of target sequences containing each motif, as well as the number and percentage of background genomic sequences with the motif.
